# A definition schema for life level and some associated models

**DOI:** 10.1101/2023.08.02.551611

**Authors:** Attila Losonczi

## Abstract

We present a mathematical definition schema for measuring life level that is to give options to describe how much a (generalized) organism is alive. It is just a definition schema, not a definition. To make it a real definition configuration is required by applications.

We devote a long section to various examples where we demonstrate how a ready made definition can work and what ways it can provide reasonable values for life levels.

In the second part of the paper we outline many possible ways how one can partially derive life-level values from some simpler principles using probabilistic approach and various other methods. All of those models are still going to stay on a general level however they will suggest some more specific methods.

From technical point of view, we also present various kinds of operations which produce new life level functions, life level systems from already existing ones, and also highlight connections among systems / functions. Some operations also extend the usability of the original life level function, others add nice (expectable) features. We also examine the properties of such operations and the relations among them. Then we apply those techniques for some biologically meaningful extensions / restrictions of life level functions and systems.

We also deal with various averages on already defined life level functions and see how they can be applied, and we also investigate metrics (biological distance functions) that are synchronized with a life level function.

We also build a basic theory about measuring the vitality of sub-organisms regarding a containing (generalized) organism, and we identify some links between this notion and the life level concept.

## 1 Introduction

In his famous book, *The problems of biology* ([28]), John Maynard Smith was dealing with the definition of life in the first chapter of the book, emphasizing the utmost importance of the problem. Before and after, there were and still there are many attempts to find one single and final definition for life, where “*life*” is meant in the original, usual sense. Those attempts sometimes followed different paths, different attitudes and sometimes they were more or less chasing different goals.

In this paper we are not going to solve the original problem, instead, we will present a significantly different viewpoint about the base of that problem; in some sense, we will rephrase, alter and refine the original question, and we will find answer(s) that question.

The following introduction is divided into six subsections. We suggest that the reader should read at least the first three ones in order to get a rough picture of the content of the paper. The first one contains some very basic ideas, the second one is a more or less detailed description of the aim and scope of the paper, while the third one sketches how the definition schema is build. In the introduction, everything is explained without mathematics, therefore here, the reader should not expect high rigor and precision.

### 1.1 Some preliminary thoughts on the paper

#### The two main aims of the paper

1. To present a mathematical definition schema for measuring life level of (generalized) organisms.
2. To provide some associated models that fulfill the requirements of the definition schema and provide reasonable life level values.

A little bit of explanation:

1. What we will present is just a definition schema, not a definition. It is a kind of framework that all such definitions should follow. In that sense, one may also say that **the definition schema is an axiomatic definition of life level**. Therefore, configuration is required in order to get a real definition. This kind of configuration can be made in several ways, hence many such definitions can exist. A ready made definition can be used for measuring life level of assorted (generalized) organisms in many aspects.
2. The models represent possible ways of configuration of the definition schema. Each of them is based on some basic principles, and the life level values are derived from them.

#### An important secondary aim of the paper

Furthermore we want to build a reasonable mathematical formalism that is capable to describe biological systems from aliveness perspective, a mathematical framework where various such related questions can be formalized precisely and answered accurately.

#### Some warnings against possible expectations on the paper

Let us emphasize again: this paper does not answer the question what life is. It does not want to. It is going to build a theory that is applicable in a different level.

Also essential to mention right at the beginning that the definition schema we will provide is purely mathematical. Mathematical in the sense that we project the real world to mathematical notions and then *there* we make definitions and create statements.

What we currently present is a descriptive theory and a few general models with some modest statements and some connections among the new concepts.

#### A few words on the original life concept versus the current model

The most important part of this introduction is that we emphasize here that we are not going to replace the usual notion of “*life*” with a new one in any sense. We do not think that the usual notion is outdated and/or needs amendment. We respect the quest of biologists for finding a generally accepted definition for “*life*”. We are not going to challenge that in any way. Instead, in this paper, we are going to present a new attitude, a new way how one can also think of “*life*”. It is not a competitor to the original notion, instead, it is another different way how one can approach this hard problem. If this new way will also succeed, then it will not decrease the importance and significance of the original notion. In other words, we think that both notions represent important models for biological systems. Models that have common roots, but models that express different attitudes on describing that part of the world. After presenting the basic mathematical concepts of the paper, especially the definition schema, in subsection 2.5 we will present a detailed comparison between the original life concept and our new model, while here we just sketch some differences but without deeper explanation.

There is no commonly accepted definition to the notion of “*life*”. However despite of the fact that there is no definition, the ways how people think of “*life*” are not too far from each other. There are differences, however there is a common ground and basically they are similar regarding the base underlying concept. In this paper we are going to use the word “*life*” differently, not very much differently, but the differences will be essential in some sense. Differences, because there will be many such. While reading this paper please always be aware that from now on, the words “*life*” and “*alive*” will present that different meaning that we will explain and present in detail later.

We think that there is no a priori answer to the question whether a thing is alive or not. It cannot be <u>simply</u> read from the organism or from the properties of the organism. “*Being alive*” is not such a property as e.g. color or weight. Therefore it is much easier and perhaps more natural to compare the level of “*aliveness*” among several assorted organisms. We are saying here that comparison may play a key role in a new type model, in other words, one can express “*life*”-type relations among organisms.

We think that from our new point of view, the good question is not that: “*is it alive*?”. Instead: “*how much is it alive*?” Strongly stated, we would say that life has grades, many many grades.

Our new model is comparison based that means that the actual values assigned to organisms do not matter (or just slightly), what matter are the ordering relations among the values.

#### The scope of investigated organisms

We want to allow to investigate various range of (generalized) organisms from life level perspective, much more than the original life concept covers. Of course, one can investigate the usual “mainstream” organisms, but rare natural cases can be also examined, and also artificially modified or (partially) artificially created organisms as well. Likewise, we are interested in snapshots of organisms at their several development life phases.

Partially because of that wider range, comparison can get more importance.

#### On parameters

In certain models there may be many important relationships, better to say parameters that should not be excluded when someone wants to have a life valuating function, in the sense that those (normally hidden) parameters may have significant effect on life level values. Depending on our model, without those parameters one may end up with false or at least not too correct results. Hence we are going to add all of those parameters to our definition schema. Namely time, relations to some groups of organisms, dependence from some part of the environment. Of course life-level models can exist which can deal without considering some of these parameters, however we think that more sophisticated models should take these dependencies into account.

We can rephrase the previous thoughts in a way that one has to answer all of the following questions when one is about to measure life level:

- What exactly is the object am I investigating? What are its borders? What belongs to it and what does not? This question seems to be too trivial to listen to, however better to be careful because it may significantly affect the result.
- What / which environment does the object reside in during the period of investigation?
- In what group of “similar” objects does the object reside during the period of investigation which can alter its life level through various connections?
- When exactly does the investigation take place? Life level is time dependent.
- And finally and most importantly: Under those constraints what is the life-level of the examined object?

A model may work without taking into account the effect of the environment and/or the supporting group of similar objects, however all of those parameters are built into the definition schema for more general possible applications.

Let us also quote John Maynard Smith ([27] Chapter 3) who writes about fitness (that is evidently related to life level): “*Fitness is specific to an environment. Thus in my lifetime, myopic individuals have had a high fitness, because they were less likely to be put in military uniform and shot at. In a hunter-gatherer society, myopics would probably have a low fitness*.” Of course, in his example, he did not differentiate the sub-environment and the supporting-group (as we do), but clearly he meant both as such that can affect the value of fitness.

#### What can such model measure?

First of all, we want to create a tool that is able to measure life level of various (generalized) organisms. Again, we want to measure not also the “mainstream” organisms, but also peripheral ones, e.g. under development, damaged, modified, etc. Also we want to see the change of life level through time. We are also interested in the sensitivity of life level for the change of both the environment and some related groups of organisms. And also we want to see how life level evolves when an organism develops through various life phases.

#### Why just a definition schema?

Why do we present just a definition schema and not a definition? We think that it is not possible to create one <u>final</u> life-level function that is applicable everywhere and for every situations. It is mainly because there are different aspects how organisms can be seen and investigated, and there are also many different models based on various principles that all provide reasonably good values. Certainly we should not distinguish among the different viewpoints, we should not say that this one is better, the other is worse, because every viewpoint can have its own advantage and purpose. The only thing that we can provide is some general viewpoints, general guidelines on how such definitions can be, or better to say, must be formulated when someone is about to have a life-valuating function for some parts of the objects of the world that one is examining.

Of course a real definition can be made in several ways according to the need, scope and purpose of a research project. Therefore it may happen that one definition may say that the same object is alive “*very much*” however the other may say it is hardly alive. In an extreme case it may also be possible that an object is “*alive*” according to one definition however it is not “*alive*” according to another. It is not a failure of the method. Simply we cannot expect more. The level of liveness or just being alive may depend on the viewpoints that one or other research group chooses.

We also want to allow that a definition may be created for a given ecosystem only, even for a smaller one. We do not think that a proper life level type definition necessarily must cover the whole biosphere, or it must work everywhere. Instead, it is possible that many partial definitions may cover the whole biosphere.

#### Models

Apart from the definition schema, we will provide several general models too. Models that accomplish the requirements of the schema and provide reasonable and expectable results. All of those models will stay on a general level, in other words they do not want to be very specific. All of them are based on some general principles from which we can derive life level values for certain group of organisms. Details come in the next subsection.

An important remark: In this paper we will use the word “model” in two senses. When some application applies this theory and creates its own life-level-system and life-level function, then the product is a model (of some biological system). The theory described in this paper is also a model: it is a model of models, i.e. a model about how to create and maintain concrete biological models. It is good to be aware of those two distinct meanings.

Finally we cannot deny that the paper has some philosophical aspects too (which is unavoidable), however our aim is to stay on the mathematical ground as much as possible.

### 1.2 The aim and scope of the paper

In this subsection we would like to outline the main aim of the paper **without any mathematics**. Firstly we explain the (biological) problems we look for solutions, secondly we describe how our approach might be useful for solving those problems. Because of the lack of mathematics, it might be ambiguous. We hope that if the reader gets interested by this brief introduction, then he/she will go on studying the mathematics presented in this paper, and therefore he/she can eliminate the confusion.

Before starting, let us emphasize again that the new concept defined in this paper is fairly independent of the original life concept. Its main aim is not to extend or modify the original life concept, rather to provide a new independent model. Please keep that in mind while studying this section and the rest of the paper as well.

#### 1.2.1 Problems

Let us start with addressing a list of problems that we are looking for (a kind of) solution. We also highlight some of our expectations. Here we will not provide detailed descriptions and explanations; they will come in subsequent sections. E.g. the reader can find additional problems in the Examples section (section 3) whose one of its aims is to enrich the set of such problems and also to provide more detailed presentation of the problems mentioned here.

1. Let us enumerate a few examples where the original life concept may be challenged in the sense whether those organisms are alive or not. Consider a virus, a mitochondrion, a totally frozen freeze tolerant insect, a dormant tree in winter, a man at the state of clinical death, a sperm, a dormant seed, an infertile mule, a worker bee, a man with an implanted cardioverter defibrillator, and one of two organisms living in obligatory symbiosis. In our new model we would like to assign non zero life-level values to those scenarios, despite of the fact that those organisms are not “fully” alive (whatever that means). Remark for dormant tree and seed: They are mostly considered alive by the original life concept, despite of the fact that they both lack total metabolism for a long time, and metabolism is considered as one of the most important life criteria (by the original concept).
2. A muscle cell of an animal may be expected to have lower life-level than a bacterium because e.g. it does not have the capability to inherit its properties.
3. One may ask the following questions. Is a 4 month old fetus not alive? Or is a new born not alive? Or is a 90 year old man not alive? One can ask those questions because none of the previous organisms satisfies at least one (or usually more) very basic characteristics that are usually assigned to being alive (in the original sense). In our new models we want a kind of measurement for what is missing and how much the missing parts affect life-level.
4. We prefer to have a system that can distinguish the life-level of a zygote, a fetus, a newborn, a cub, an adult and an old animal. Of course the granularity can be even finer.
5. Also we would prefer to have a lower life-level value for a man in coma than being fully healthy.
6. A part can have different (usually lower) life-level than the whole, e.g. a muscle cell vs. the whole muscle vs. a man. Consider also a tissue culture or an organ culture. A tissue that is kept alive artificially must have lower life level than the parent organism from which it was separated from. If an organ is grown (cultured) in vitro, then a non-zero life level can be associated with it, especially, if it can be transplanted into an organism. The life level of a cut arm of a starfish which is capable for full regeneration should be less than, but close to the life level of the starfish.
7. We would like to assign higher values to groups of organisms, e.g. a big herd of zebras must have higher life level than a single animal, or a small but complete ecosystem must attain a higher figure than its smaller components.
8. We would like to give the option to take into account the effect of the environment on the life-level, because it can significantly increase or decrease it. In many cases, when the environment is “taken away” then the organism dies almost immediately. E.g. consider these examples: a fish living in a poisonous lake; a person on a breathing machine; a man with a built in artificial cardiac pacemaker; a man living in a town or in a rain-forest alone or on Mars. An even more profound example is if one examines a fish in the water or on the ground (just taken out from the water).
9. We also want to have the option to consider the so called supporting group that can have essential effect on life-level. Similarly in some cases it can be so essential that without it the organism dies in a very short time. E.g. consider these examples: one of the organisms of the two, living in obligatory symbiosis; a young cuckoo raised by its host parents; a completely disabled man with or without people who supply him. A chloroplast with or without its host cell (as its supporting group) can possess very different life level values. The bird kakapo can only breed when rimu trees put out masses of fruit.
10. If one has a life-level function, then it can also be a natural requirement that it has to be time dependent, in other words its values may depend on time. E.g. consider these examples: the life-level through life-span of an organism; a man before a serious injury, in the recovery state, and after complete recovery; a sperm before and after fertilization; some light poison gradually ruins the health of an organism and finally kills it.
11. Continuing the previous point, if one is about to examine the life level of an organism for a very long period of time then one has to face a difficult problem that we may call the identification problem. An organism is always changing. It changes many of its smaller or bigger parts, it changes its traits or even some of its characteristics. There are also smaller parts of the organism which seems to be totally intact, but a closer look can immediately reveal that it is interchanging rapidly, nevertheless it looks the same. More importantly one may see scenarios where two organisms unit into a new one, or the other way around, one organism splits into two new ones. One more such example is a man with a heart transplant; now one of his organs has been changed with a new one that has different DNA. Nevertheless, we consider him to be the same, and can ask about his life level before and after the transplantation e.g. In all of those cases, a developer of a reasonably good model has to make a decision on which “things” belong together in time, in other words, which “things” represent one single development path of the “organism” regarding life level examination.

First we need to have a fairly general way that is capable to describe all of the previous phenomena. Describe in the sense, that the life-level function has to have the capability to express those relations properly. The three definition schemas are devoted to serve that purpose.

On its own, the three definition schemas do not provide any specific ways about how to assign real values to (generalized) organisms. In that sense the definition schema is just a framework that needs configuration in real applications. When an application is just about to build its own life level function, it has two options (that can be mixed of course). It can assign ad hoc values to organisms, or it can find general basic guidelines which somehow determine similar values that the application expects.

#### 1.2.2 Models

In that direction, in the second part of the paper, we are going to suggest several models that are based on some general principles that induce life level values in accordance with our expectations. Those models are supposed to answer (at least partially) the previously outlined questions / problems. It is important to emphasize that different models may provide partially different answers in some cases, but we cannot expect more.

Let us sketch the basic versions of some of those models briefly.

1. One can have a list of properties (roughly attributes of generalized organisms), which one thinks that are essential in measuring life. One can assign weights to all of those properties, and summarize those weights which belong to properties a given organism owns at a given point in time. If the properties are carefully selected, then this model can give a small non-zero value for a totally frozen freeze tolerant insect, a dormant seed/tree, and a man at the sate of clinical death. It can also provide notably smaller values for an infertile mule, a worker bee, a 4 month old fetus, a 90 year old man, and a man in coma. It can provide smaller values for a virus, a mitochondrion, and a completely disabled man. It can assign a bit higher value to a herd of zebras than to a single zebra.
2. A property usually is not active always, it may switch on and off as time goes by. Thus one can enhance the previous model by just requiring from a property to hold (at least once) in a given time interval only. This model can offer a high value for a totally frozen freeze tolerant insect, a dormant seed/tree, and a man at the sate of clinical death. In all the other cases, it yields similar values.
3. A further enhancement can be got when one takes into account the activation time of properties (also with applying some probability if needed). This model can give a higher value than in point 1 but lower than in point 2 for a totally frozen freeze tolerant insect, a dormant seed/tree, and a man at the sate of clinical death. In all the other cases, it yields similar values.
4. Again we have a list of properties with associated weights. For a given (generalized) organism, instead of examining the present, one may estimate the survival time of each property on the development paths of our organism, meaning how long in time the property is owned by the organism or its successors. Then simply take the weighed average of the property weights and the survival times, and it may serve as a life level function. This model can provide high values for a totally frozen freeze tolerant insect, a dormant seed, a man at the sate of clinical death, and a 4 month old fetus; and an even higher value for a dormant tree. However it may provide lower values for an infertile mule, a worker bee, a 90 year old man, a man in coma, and a protoplast. There may not be significant difference for a virus. It can assign a higher value to a herd of zebras than to a single animal.
5. The previous model based on estimations on future values. Hence we may get a more sophisticated model if one enrich the previous model with probability values on the possible future development paths of a given organism. We take that into account that the probability values modify the survival time in the self-evident way. Then we again take the weighed average of the property weights and the new survival times and we end up with a life level function. In contrast to the previous model, this model may provide much lower values for a man at the sate of clinical death, and a man in coma. It may provide lower values for a 4 month old fetus, a dormant seed, and a protoplast. There may not be significant difference in the numbers for an infertile mule, a worker bee, a 90 year old man, a totally frozen freeze tolerant insect, a dormant tree, a virus, and a group of zebras.

One can enrich those models, if one refines the property-owning relation to a function which expresses how much a (generalized) organism owns a given property, i.e. it is not an owned or not-owned relation any more, instead its value says how much the property is owned.

#### 1.2.3 Operations

A different approach is that when we can get more complex or better functioning models from simpler ones. More precisely, if one have a life level function on a set of organisms then one can either extend the function to other generalized organisms, or using the original values, one can derive new, more reasonable life level values for organisms (that might have had values originally). We sketch a few of those kinds of operations.

1. Suppose that we have a life level function defined on “single” organisms only. We can extend that to more complex e.g. group type organisms in a way that we take the maximum life level values of the organisms belonging to the group. Another possible option may be that instead of the maximum, one can take the sum.
2. Take the supporting group of a given organism, and calculate the maximum life level values of all organisms that belong to the supporting group, and assign this value to the underlying organism. A more sophisticated model may take into account weights, that is how much an element of the supporting group increases the life level of the underlying organism comparing to the other elements. Using those weights, one can take the weighted average, instead of max.
3. Modifying the previous operation, instead of the supporting group, one can take the reverse supporting group that consists of those organisms that the given organism supports. This model provides reasonably good figures for an infertile man whose existence is essential for some other (fertile) men and women.
4. One can take a kind of average of the future values of the the life level function by a weight function, and project the calculated average to the present and use it as a value for a new life level function. With that method, one can make the life level curve more smooth with keeping several other properties. This method also serves as a (weighed) projection of the near future to the present, i.e. if one has a life level function that takes into account the present only, then the new derived function compresses information from the near future and the present, and use it as a present value.
E.g. if an organism lives close to a just erupting volcano, then in this model the eruption itself can decrease the present life level.
5. There are some further technical operations that fix some flaws in life level type functions, functions whose values lack some essential properties.

Models 2 and 3 express that (a certain) group can have significant affect on the life level, it can even determine it, or at least partially.

#### 1.2.4 Averages

We will also present some averages on already defined life level functions.

1. In the future an organism can “choose” several different sub-environments where it can reside to, and also several supporting-groups which it can belong to. Based on that and using probabilistic bases, we can estimate the future life level of the organism. Of course this is a kind of average (based on probability), that is why this is presented here.
2. One can also calculate life level averages on time intervals. Or also for the whole life-span of an organism.
3. If we have a group of organisms, then one can assign average-type life level values to the group based on the life level values of organisms which consist the group.
4. The usual moving averages and some of their properties are also examined. We remark that this can be considered as an operation too.
5. Operation 4 is about smoothening the life-level curve and in that sense can be considered as an average.

#### 1.2.5 Synchronized distances

If a project has some biologically meaningful distance on the set of all organisms (and/or sub-environments, supporting-groups), then it is worth investigating connections between life-level functions and that biological distance function. We call a life-level function and a distance synchronized if the closer the objects (organisms, sub-environments, supporting-groups) are, the closer the associated life-level values will be. We will examine such relations.

#### 1.2.6 Vitality measurement

One can approach to the general concept of life-level through the notion of vitality as well, that is a kind of measurable relation between a sub-organism and a containing (generalized) organism. It is supposed to express how much a sub-organism is important for the life of a containing generalized organism. For example this is a generalization of the notion of vital organ, and it can also be used to measure the vitality of a member of a group of gorillas regarding to the whole group.

This approach can be compared to the life-level concept and see if one concept can be derived from the other, and also the associated similar properties can be investigated.

### 1.3 How the definition schema is built

Here we are going to sketch the inner structure of the definition schema, but only roughly, without any mathematics. There are three definition schemas. In all of them the examined object is a quadruple, that is the combination of an organism, a part of the environment, some fellow organisms and a time point. Hence we never examine an organism “alone”. We assign a life-level value to such quadruples. The domain of a life-level function, in other words, which such quadruples are investigated together, is up to the application.

In the **simplest schema**, it is assumed that a so small time interval is investigated that nor the organism, no the environment, no the supporting group changes essentially. This schema provides option for two things:

1. Describe life-level through time.
2. Taking into account the effect of the environment and the supporting group.

We remark that one can get an even simpler model if one omits the effect of the environment and the supporting group i.e. both are empty.

In the schema with **medium complexity**, it is still assumed that nor the organism, no the environment, no the supporting group changes essentially in time, however we allow here that the organism changes its environment and / or supporting group as time goes by.

In the **most complex version** of the definition schema everything (organism, environment, supporting group) is allowed to change in time. Moreover the organism may change its environment and / or supporting group too. Here we need to formulate development paths that are kind of links through time among things that belong together. So one development path expresses how an organism changes through life forms and still stays the “same” thing. The same logic is applied for the environment and the supporting group. Moreover these three things (organism, environment, supporting group) change together as time goes by, and together express one single development path. In this model version the life-level value still belongs to the quadruples (organism, environment, supporting group, time), but this additional formalism provides more precise description of the world. For example one can easily describe unitness or splitting of organisms. Or one can also handle many optional development paths at the same time. Or in this case one can also assign probability for the optional paths, and use it for more sophisticated models.

In order to be the most general as the definition can be, here we do not identify organisms / environments / supporting groups directly, the identification is done by the development paths only. It means that at different point in time we assume that every organism, every environment, every supporting group is different, even a small time later. In other words at every time point we have a big set of organisms, environments, supporting groups. And the development paths choose the ones (at different time points) that belong together. That is the formalism that we apply here.

One more element of the definition schemas is that we need a partial order among the organisms existing (in our model) at a given point in time. We want a relation that expresses that some simpler (generalized) organisms are parts of some more complex ones.

Similar requirement hold for the sub-environment structure, hence a partial order is applied there too.

### 1.4 The structure of the paper

The paper is divided into two parts. In the first part (this introduction and sections 2 and 3) we provide the basics of the definition schema, while in the second part we add many more details and deliver various models fulfilling the requirements of the definition schema.

In section 2 we present the definition schema, better to say three of them because we give two simpler versions too that may suit for more modest applications. Here we provide some basic definitions and statements related to the schema. We then make many remarks in order to clarify the definitions and the new notions inside them. We also make a detailed comparisons between the usual life concept and this new model and highlight the differences (see 2.5).

In section 3 we provide many many examples that are supposed to support the understanding of the model. The examples will be deliberately not worked out properly. Their aim is not to give final values, the aim is just to perceive how reality may be projected to the model and through them get a better understanding how the model can be built.

In the second part, in section 4 we provide many possible ways how one can partially configure this definition schema. We will suggest some ways how life-level can be calculated from some easier assumptions using probabilistic and other methods. All of those are based on properties that a generalized organism can or can not own at a given point in time. Here we just outline one such method: the life-level can be calculated by the survival time of given properties. Those methods are still going to stay on a general level, however they will suggest some more specific method. They will get closer to the original concept of life, but only closer.

In section 5 we examine algebraic operations on life-level-systems. First, we will investigate the extension of life-level-systems, i.e. when one extends the domains of one’s life-level function; we will also examine how one can unit two life-level-systems into a new one. We will also study biologically relevant mappings between life-level-systems, and using that, we will define the equivalence of life-level-systems.

Then in section 6 we define many operations on life-level functions, functions only. The first group contains some elementary (mainly algebraic) operations, the second group of operations is about to fix flaws of life level type function, while the third group is related to averages.

Section 7 presents some biologically meaningful operations, e.g. operation which somehow extends the original function with the aim that the supporting group can determine life-level as well. Another example of such operations deals with life-level functions that are defined on non group-type organisms only, and we try to find natural ways for extending life-level functions to group-type organisms too.

In section 8 we define various averages on already existing life level functions. One such type takes averages on time. The other takes averages on certain groups of organisms. In the most complex average, we will be dealing with how one can get better estimations for life level values in the future. The method is based on calculating averages on probabilistic bases.

We remark that in section 6 we also add average-type operations (e.g. when we smooth the life level curve or when we deal with moving averages). Section 9 presents some methods how one can assign life-level values to species <u>if</u> some life-level function is already given, i.e. we derive the values from already existing life-level functions.

In section 10 we investigate connections between a given biological distance on organisms (and/or sub-environments, supporting-groups) and life-level functions.

Section 11 is devoted to building a basic theory on measuring vitality of sub-organisms regarding a containing (generalized) organism, and also investigating its relations to life-level functions.

In the last part of the paper (section 12) we add some remarks / suggestions on possible generalizations of the model that may be considered by applications.

As you can see from the structure of the paper, **the theory is built deductively**, in other words we first outline the abstract model and then we provide explanation, examples and suggestions about the usage. We have deliberately chosen that way, even if it is more difficult to grasp at first sight.

### 1.5 Some historical remarks on the original concept

Here we are going to enumerate some of the previous life definitions with some short excerpts and very brief and not complete explanations.

There are two types of attempts:

1. (more or less) purely mathematical
2. structural, property/process based.

#### Mathematical type definitions

[22, Rosen] and [23, Rosen] suggest a relational approach to distinguish between alive and not alive. Actually the way how the components of a system interact with each other determines if a system is complex or simple. And high complexity is considered as alive, while low as not alive. See also [26, Siekmann] for deep, profound and very detailed explanations.

Based on the ideas of [19, Neumann], living beings can be considered as a special physical realization of the mathematical idea of a self-reproducing automata.

[3, Chaitin] and [4, Chaitin] proposes and discusses a measure of degree of organization and structure of geometrical patterns which is based on the algorithmic information theory. Hence here the notion of life is more or less identified with complexity as well.

[17, Mack] uses Gauge theory which is an application of category theory, to define processes that are elementary for biological life.

#### Structural, property/process based definitions

In [28] John Maynard Smith provides necessary properties of a living system: A living organism must maintain a constant controlled flow of energy through the system, it must have metabolism, it must own properties of multiplication, variation and hereditary.

In [2, Campbell] there is a list of properties that are characteristic to living systems: Order, Reproduction, Growth and Development, Energy Utilization, Response to Environment, Homeostasis, Evolutionary Adaptation. Actually [2, Campbell] does not consider this as a definition, instead, just emergent properties and processes of life.

[11, Gánti] has two groups of criterions, absolute and potential ones. The absolute criterions are: a living system must inherently be an individual unit; a living system has to perform metabolism; a living system must be inherently stable; a living system must have a subsystem carrying useful information for the whole system; processes in living systems must be regulated and controlled.

Potential criterions are: a living system must be capable of growing and multiplying; a living system must have the capacity of hereditary change, furthermore the capacity of evolution; living systems must be mortal.

The aim of [29, Smith, Szathmáry] is not to present a proper definition, however it wants to describe and analyze two main types, two alternative definitions: phenotypic and hereditary ones. Phenotypic means that a living system is a self-regulating complex machine-like entity with the feature of metabolism. While hereditary means that a living system must have the properties of multiplication, variation and heredity.

[1, Akkerhuis] uses the common denominator of the “operator” for a unified ranking of both particles and organisms, from elementary particles to animals. This ranking is called “the operator hierarchy”. Using this hierarchy, life can be defined as matter with the an operator that possesses a complexity greater than or equal to the complexity of the cellular operator.

[18, Ruiz-Mirazo, Peretó, Moreno] proposes a definition that is built on two properties: “a living being” is an autonomous system and it has open-ended evolutionary capacities. Autonomy regards for self maintenance. While evolutionary capacity means two thing: Reproducing its basic functional constitutive dynamics and also creating limited variety of equivalent systems.

[35, Zhuravlev, Avetisov] composes a definition from three components: 1) a living system is a specific state of matter and energy 2) it has a specific hierarchical system consisting of self-reproducing agents 3) it goes through a specific process that is expressed in transformations of surroundings and in transmutations of the self-reproducing agents themselves.

[10, Forterre] argues that a virus can be considered as a machinery that produces (better to say encodes) some generalized organs and based on that, a life definition can be formalized that viruses fulfill.

According to [7, Crick], the requirements for life are the followings. The system must be able to replicate directly or by “using other machinery”. The replications must be exact, but mutations must occur at a rather low rate. A gene and its products must be kept reasonably close together. The system must have a supply of raw material and free energy.

In [31] Szent-Györgyi identifies some quantum-physical type properties of living-systems that are unique to the investigated systems.

#### Miscellaneous

There are some survey articles as well, e.g. [34, Trifonov] that also tries to identify common properties of the already existing definitions.

[33, Stewart] argues that life should not be defined by just the laws and properties of genetics. It says that mathematical patterns should provide the building blocks and the very fabric of life. It does not give a proper definition, but raises many deep questions and it analysis several life related mathematical patterns.

Finally, we also need to mention [16, Lovelock], which makes stress on considering much bigger objects as living beings in some sense. Much bigger means that bigger than usual “bodies”. Actually, it deals with generalized organisms, meaning, small and big special groups of usual organisms. More precisely it also considers smaller or bigger ecosystems or even the whole biosphere of the Earth as a living being in a certain sense.

### 1.6 Summary of mathematical notions and notations applied in the paper

We will use the usual abbreviations: ∃ for “*exists*”, and ∀ for “*for all* “. They are just simply abbreviations without any further meaning.

ℝ, ℝ^+^, 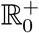, ℕ, ℕ_0_ will denote the set of real numbers, positive real numbers, non-negative real numbers, positive integers, non-negative integers respectively. We will also use the notation 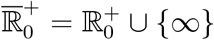 with extending the sum and the order in the usual ways: 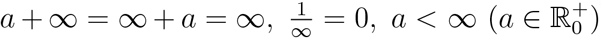.

For a set *H* ⊂ ℝ, inf *H*, sup *H* will denote the infimum and supremum of *H* respectively. We will use the conventions that inf ∅ = +∞, sup ∅ = −∞.

The **sign function** is defined as

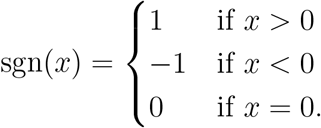

If *x* ∈ ℝ, then ⌈*x*⌉ denotes the smallest integer that is greater than or equal to *x*, i.e. ⌈*x*⌉ = min{*z* ∈ ℤ : *x* ≤ *z*}.

For a set *H, P* (*H*) will denote the **power set** of *H* i.e. *P* (*H*) consists of all subsets of *H* i.e. *P* (*H*) = {*K* is a set : *K* ⊂ *H*}.

A set *H* is said to be **countable** if it is either finite or countably infinite. We use the usual conventions sup ∅ = −∞, inf ∅ = +∞.

If *H* ⊂ ℝ is a countable set, then Σ *H* will denote the sum of its elements i.e. 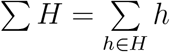.

A system of sets ℋ is **closed for finite union** if *H*_1_ …, *H*_*n*_ ∈ ℋ implies that *H*_1_ ∪ … ∪ *H*_*n*_ ∈ ℋ. It is closed for finite intersection if *H*_1_ …, *H*_*n*_ ∈ ℋ implies that *H*_1_ ∩ … ∩ *H*_*n*_ ∈ ℋ.

Let **𝒯** be a set. The binary relation ≤ on **𝒯** is called a **partial order** if it is reflexive (*a* ≤ *a*), antisymmetric (*a* ≤ *b, b* ≤ *a* ⇒ *a* = *b*) and transitive (*a* ≤ *b, b* ≤ *c* ⇒ *a* ≤ *c*). It is called a total order if ∀*a* ∀*b a* ≤ *b* or *b* ≤ *a* holds. We will use the notation⟨**𝒯**, ≤⟩, where **𝒯** is a set and ≤ is a partial order. In this paper we will often use the symbol ⊂ instead of ≤.

In a partially ordered set ⟨**𝒯**,≤⟩, *m* ∈ **𝒯** is called a **maximal element** if *m* ≤ *a* ∈ **𝒯** implies that *m* = *a*. In a partially ordered set there can be many maximal elements, or also none.

⟨**𝒯** ≤⟩, is called a **lattice** if every two elements has a supremum (a least upper bound) and an infimum (a greatest lower bound).

Let ⟨**𝒯**, ≤⟩ be a totally ordered set. Then a set *A* ⊂ **𝒯** is called **convex** if *a, b* ∈ *A, a* ≤ *c* ≤ *b* implies that *c* ∈ *A*. If **𝒯** ⊂ R then a set *A* ⊂ **𝒯** is convex iff there is an interval *I* ⊂ ℝ such that *A* = *I* ∩ **𝒯**. If **𝒯** ⊂ℝ is closed then *A* ⊂ **𝒯** is convex iff it is an interval (in **𝒯**).

Let *f* : *A* → *B* be a function. Then Dom *f* = *A*, Ran *f* = {*y* ∈ *B* : ∃*x* ∈ *A f* (*x*) = *y}*, the **domain** and **range** of *f*. Let *A*^*′*^ ⊂ *A*. Then *f* |_*A*_*′* denotes the restriction of *f* to the smaller domain *A*^*′*^.

A function 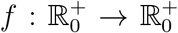 is (strictly) **increasing** if *x < y* implies that *f* (*x*) ≤ *f* (*y*) (*f* (*x*) *< f* (*y*)). It is (strictly) **decreasing** if *x < y* implies that *f* (*y*) ≤ *f* (*x*) (*f* (*y*) *< f* (*x*)).

Let a system of sets be given: {*H*_*i*_ : *i* ∈ *I*} where all *H*_*i*_ is a set and so is *I*. We say that the system is (pairwise) disjoint if ∀*i* ∈ *I* ∀*j* ∈ *I H*_*i*_ ∩ *H*_*j*_ = ∅. In this case the (disjoint) union can be denoted by 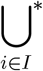 *H*_*i*_ where * denotes that it is a **disjoint system of sets**.

In ℝ^*n*^ *π*_*k*_ denotes the projection to the *k*^*th*^ coordinate, in other words *π*_*k*_(*x*_1_, …, *x*_*n*_) = *x*_*k*_ (1 ≤ *k* ≤ *n*).

Let *I* be a set and ℋ = {*H*_*i*_ : *i* ∈ *I*} be a system of sets indexed be *I*. We call *f* : *I* → ℋ a **choice function** if ∀*i* ∈ *I f* (*i*) ∈ *H*_*i*_.

For a given set *X*, a set function *μ* : 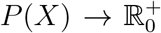 is called **additive** if *A*_1_, …, *A*_*n*_ ⊂ *X, A*_*i*_ ∩ *A*_*j*_ = ∅ (*i* ≠ *j*) implies that 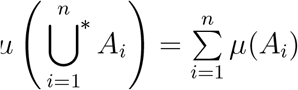. It is **sub-additive** if *A*_1_, …, *A*_*n*_ ⊂ *X* implies that 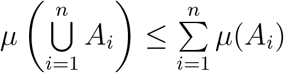.

A **multiset** (or mset) is a pair ⟨*H, m*⟩ where *H* is a set and *m* : *H* → *ℕ* is a function (see [32]). It generalizes the common notion of set in a way that it allows multiple occurrences of elements of a set. When the elements of a multiset are given, then we use the notation {… }_multi_ instead of {… } that is for “normal” sets. E.g. *M* = {1, 1, 1, 2, 5, 5}_multi_ is a multiset with support set *S* = {1, 2, 5}. The cardinality of a multiset *H* is denoted by |*H*| similarly for normal sets. E.g. for the previous examples |*M* | = 6, |*S*| = 3.

A **pseudo-metric** *d* on the set *X* is a function *d* : 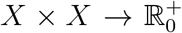 such that *d*(*x, x*) = 0, *d*(*x, y*) = *d*(*y, x*) and *d*(*x, z*) ≤ *d*(*x, y*)+*d*(*y, z*) hold. We say that a sequence (*x*_*n*_) of points of *X* converges to *x* ∈ *X* when *d*(*x*_*n*_, *x*) → 0 holds. In this case we will also use the notation 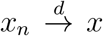. The **topology** generated by the pseudo-metric *d* is denoted by *τ*_*d*_ where a set *N* is open iff ∀*x* ∈ *N* ∃*ε >* 0 such that {*y* ∈ *X* : *d*(*x, y*) *< ε*} ⊂ *N*.

## 2 The definition schema(s)

Before we start, let us enumerate the **primitive notions** according to our theory i.e. concepts that are not defined in terms of previously-defined concepts. They are:

1. organism / generalized organism
2. sub-environment
3. gorganism-particle
4. environment-particle
5. time
6. property (see 4) (“*bio-bit owns a property* “ is a primitive relation)

We will present two types of the definition schema; in the first one (definition 2.1.10) 1 and 2 are primitive notions, and 3 and 4 are not present, while in the second one (definition 2.1.22) 1 and 2 are not primitive, they are derived from 3 and 4 respectively.

Despite of the fact that the notion of supporting-group is defined in terms of more elementary notions, it is still more or less primitive to some extent.

### 2.1 Presenting the definition schemas

We are going to provide three versions of the definition schema. One generic and two special cases. Later, in subsection 2.1.1, we will provide some more elementary forms.

First we need some preparation: we need to define the objects for which a life-level value can be assigned i.e. the possible domains of a life-level function, and also relations among the elements of the domain.

#### Definition 2.1.1.

*The structure* Λ_*sb*_ = ⟨⟨**𝒪**, ⊂⟩, ⟨**𝒦**, ⊂⟩, ***𝒢*, 𝒯**, Θ⟩ *is called a* ***life-level-subbase-system*** *if* **𝒯** ⊂ ℝ *is closed*, ∀*t* ∈ **𝒯** (*t*), ***𝒦***(*t*), **𝒢**(*t*) *are sets such that*

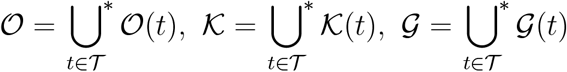

*and*

1. ∀*t* ∈ **𝒯** ⊂ *is a partial order on* ***𝒪***(*t*),
∀*t* ∈ **𝒯** ⊂ *is a partial order on* **𝒦**(*t*),
2. ∀*t* ∈ **𝒯** *there is a least element in* ***𝒪***(*t*) *that is denoted by* ∅,
3. ∀*t* ∈ **𝒯 𝒢** (*t*) *P*(**𝒪** (*t*)), *and* ***𝒢***(*t*) *is closed for finite union and finite intersection*.

*Moreover*

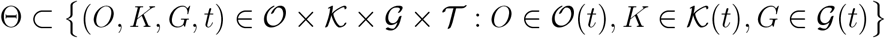

*holds*.

*Naming conventions:*

- *t* ∈ **𝒯** *is a time point*.
- *An element of* ***𝒪***(*t*) *is called* ***generalized-organism*** *or* ***gorganism*** *or simply* ***gorg*** *existing at time t*.

*If O*^*′*^, *O* ∈ **𝒪**(*t*), *O*^*′*^ ⊂ *O, then O*^*′*^ *is called the* ***sub-gorganism of O***.

- *An element of* **𝒦**(*t*) *is called* ***sub-environment*** *existing at time t*.
- *An element of* **𝒢**(*t*) *is called* ***group*** *existing at time t*.
- *The elements of* Θ *are called* ***(admissible) bio-bits*** *of* Λ_*sb*_. *If we drop the last coordinate (time) the remaining* (*O, K, G*) *can also be called* ***bio-bit (at time point t)***. *For* Θ *we also use the notation* Θ(Λ_*sb*_).

#### Definition 2.1.2.

*Let a life-level-subbase-system* ⟨⟨**𝒪**, ⊂⟩, ⟨**𝒦**, ⊂⟩, ***𝒢*, 𝒯**, Θ⟩ *be given. If b* = (*O, K, G, t*) ∈ Θ, *then we will use the notations π*(*b*), *π*_**𝒦**_(*b*), *π*_**𝒢**_(*b*), *π*_**𝒯**_ (*b*) *for O, K, G, t respectively. I.e. π*_*𝒪*_ = *π*_1_, *π*_**𝒦**_ = *π*_2_, *π*_**𝒢**_ = *π*_3_, *π*_**𝒯**_ = *π*_4_ *in this case*.

#### Definition 2.1.3.

*Let a life-level-subbase-system* Λ_*sb*_ = ⟨⟨**𝒪**, ⊂⟩, ⟨**𝒦**, ⊂⟩, ***𝒢*, 𝒯**, Θ⟩ *be given. If G* ∈ **𝒢**(*t*), *O* ∈ **𝒪**(*t*) *and there is a bio-bit b* ∈ Θ *such that b* = (*O, K, G, t*) *(i.e. π*_*𝒪*_(*b*) = *O, π*_**𝒢**_(*b*) = *G), then we will refer to G as a* ***supporting-group*** *of O at time t*.

#### Definition 2.1.4.

*Let a life-level-subbase-system* Λ_*sb*_ = ⟨⟨**𝒪**, ⊂⟩, ⟨**𝒦**, ⊂⟩, ***𝒢*, 𝒯**, Θ⟩ *be given. If b*_1_, *b*_2_ ∈ Θ, *b*_1_ = (*O*_1_, *K*_1_, *G*_1_, *t*_1_), *b*_2_ = (*O*_2_, *K*_2_, *G*_2_, *t*_2_) *then* ***b***_**1**_ *⊆*_𝒪_(***b***_**2**_ *will denote that O*_1_ ⊂ *O*_2_, *K*_1_ = *K*_2_, *G*_1_ = *G*_2_, *t*_1_ = *t*_2_. *While* ***b***_**1**_ ***⊂ b***_**2**_ *will denote that O*_1_ ⊂ *O*_2_, *K*_1_ ⊂ *K*_2_, *G*_1_ ⊂ *G*_2_, *t*_1_ = *t*_2_.

#### Definition 2.1.5.

*Let a life-level-subbase-system* Λ_*sb*_ = ⟨⟨**𝒪**, ⊂⟩, ⟨**𝒦**, ⊂⟩, ***𝒢*, 𝒯**, Θ⟩ *be given. Then γ is called a* ***development-path*** *of* Λ_*sb*_ *if*

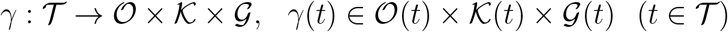

*such that*

1. ∀*t* ∈ **𝒯** *γ*(*t*), *t* ∈ Θ
2. {*t* ∈ **𝒯** : *π*_𝒪_ *γ*(*t*) ≠ ∅ *is an interval with finite left end-point*.

*For a given development-path γ, set*

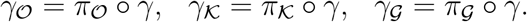

*Then γ*_𝒪_ *is called* ***gorganism-development-path***,

*γ*_**𝒦**_ *is called* ***sub-environment-development-path***,

*while γ*_**𝒢**_ *is called* ***supporting-group-development-path***.□

**Remark 2.1.6**. *Clearly γ*_*𝒪*_, *γ*_*𝒦*_, *γ*_*𝒢*_ *is a choice function from* ***𝒪, 𝒦, 𝒢*** *respectively. With this notation we get that γ* = (*γ*_*𝒪*_, *γ*_***𝒦***_, *γ*_***𝒢***_) *and {t* ∈ **𝒯** : *π*_*𝒪*_ *γ*(*t*) ≠ ∅ = {*t* ∈ **𝒯** : *γ*_*𝒪*_(*t*) ≠ ∅}.

#### Definition 2.1.7.

*For a given development-path γ, set* 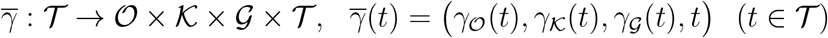

#### Definition 2.1.8.

Λ_*b*_ = ⟨**𝒪**, ⊂⟩, ⟨**𝒦**, ⊂⟩, ***𝒢*, 𝒯**, Θ, Γ *is called a* ***life-level-base-system*** *if* ⟨⟨**𝒪**, ⊂⟩, ⟨**𝒦**, ⊂⟩, ***𝒢*, 𝒯**, Θ⟩ *is a life-level-subbase-system and* Γ *is a subset of the set of all development-paths*.

#### Definition 2.1.9.

*A life-level-base-system* Λ_*b*_ = ⟨⟨**𝒪**, ⊂⟩, ⟨**𝒦**, ⊂⟩, ***𝒢*, 𝒯**, Θ, Γ ⟩*is* ***countable*** *if* Γ *and* ∀*t* ∈ **𝒯 𝒪**(*t*), **𝒦**(*t*), **𝒢**(*t*) *are countable sets*.

Λ_*b*_ *is* ***finite*** *if the previous sets are all finite*.

Now we can define the most generic version of the definition schema.

#### Definition 2.1.10.

*Let a life-level-base-system* Λ_*b*_ = ⟨⟨**𝒪**, ⊂⟩, ⟨**𝒦**, ⊂⟩, ***𝒢*, 𝒯**, Θ, Γ⟩ *be given. In the most general context* ***life-level*** *is a function* **ℒ** *that such that*

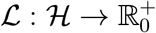

*where* ℋ = Dom ℒ ⊂ Θ, 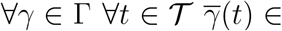 Dom ℒ, *and ℒ satisfies the following properties:*

*(es) ℒ, K, G, t* = 0 (*t* ∈ **𝒯**, *K ∈* ***𝒦***(*t*), *G* ∈ **𝒢** (*t*)

*(do) If γ* ∈ Γ, *t*_0_ ∈ **𝒯**, *γ*_*𝒪*_(*t*_0_) ≠ ∅, 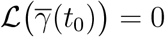, *t*_0_ *< t*_1_, *then* 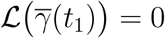.

*(ss) If O*_1_ ⊂ *O*_2_ *then* **ℒ**(*O*_1_, *K, G, t*) ≤ **ℒ**(*O*_2_, *K, G, t*) (*O*_1_, *K, G, t*), (*O*_2_, *K, G, t*) ∈ Dom **ℒ**

*Some naming conventions:*

- *The system* Λ = ⟨**𝒪**, ⊂⟩, ⟨**𝒦**, ⊂⟩, **𝒢, 𝒯**, Θ, Γ, **ℒ** *is called* ***life-level-system***.
- *We say that the life-level function* **ℒ** *is* ***associated to the life-level-base-system***. ⟨⟨**𝒪**, ⊂⟩, ⟨**𝒦**, ⊂⟩, **𝒢, 𝒯**, Θ, Γ⟩ *if properties (es), (do) and (ss) are satisfied. In this case* **ℒ** *is also called a* **Λ*-function***.
- *The function* ℒ *is just called a* **Λ*-base-function*** *or life-level-base-function if it does not necessarily satisfy all properties (es), (do) and (ss)*.

#### Definition 2.1.11.

*In the sequel we will refer to the value of* **ℒ**, *or better to say the value of* **ℒ**(*O, K, G, t*), *the* ***life-level*** *of* (*O, K, G, t*) *or when K, G, t is clear from the context then simply the life-level of O*.

**Remark 2.1.12**. *The general term for the elements of* ***𝒪*;** *is gorganism. We have the term bio-bit for* (*O, K, G, t*) *and that is the object for which a life-level value can be assigned*.

**Remark 2.1.13**. *If* Λ *is a life-level system, then in the sequel we will use both formalisms* **ℒ**(*O, K, G, t*) *and* **ℒ**(*b*) *where b* = (*O, K, G, t*) ∈ Dom **ℒ**.

#### Definition 2.1.14.

*Let a lif e-leve l-syst em* Λ *be given and let γ* ∈ Γ, *t* ∈ **𝒯**. *Then* **ℒ***γ*(*t*), *t will denote* 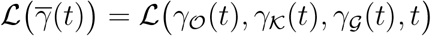.□

#### Definition 2.1.15.

*Let a life-level-system* Λ *be given. If γ* ∈ Γ *then set*

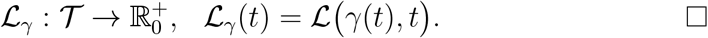

**Remark 2.1.16**. *With this notation, property (do) can be formulated as*

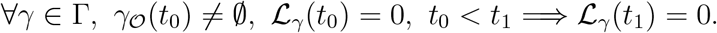

**Remark 2.1.17**. *Property (ss) can be formulated as*

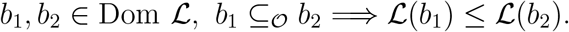

**Remark 2.1.18**. *We just mention some obvious facts that are good to be aware:*

- Γ *can be empty*,
- ∅ *can be an element of any of* ***𝒪***(*t*), ***𝒦***(*t*), ***𝒢***(*t*) (*t* ∈ **𝒯**).

#### Medium complexity version of the definition schema

We immediately give a simpler version that may be suitable for several not-too-complex applications. In this version we may assume that the examined time period is so small that the investigated gorganisms, sub-environments, supporting-groups can be considered intact in that period. However we allow that a gorganism may change its sub-environment and/or its supporting-group. Therefore one may end up with this lighter version of definition when one uses the definition 2.1.10 with the following simplifications:

- ∀*t* ∈ **𝒯** ∀*s* ∈ **𝒯 𝒪**(*t*) = **𝒪**(*s*), ***𝒦***(*t*) = **𝒦**(*s*), ***𝒢***(*t*) = **𝒢**(*s*).
- ∀*γ* ∈ Γ ∀*t* ∈ **𝒯** ∀*s* ∈ **𝒯** *γ*_*𝒪*_(*t*) = *γ*_*𝒪*_(*s*).

Let us summarize this in the following definition.

##### Definition 2.1.19.

***In the general context, life-level** is a function* 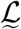 *that is defined by the following system* 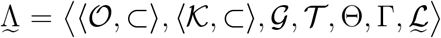 *where* **𝒯** ⊂ ℝ *is closed and*

- ⊂ *is a partial order on* ***𝒪***,
⊂ *is a partial order on* ***𝒦***,
- *there is a least element in* ***𝒪*** *that is denoted by* ∅,
- **𝒢** ⊂ *P* (***𝒪***), *and* ***𝒢*** *is closed for finite union and finite intersection*,
- Θ ⊂ **𝒪** × **𝒦** × **𝒢** × **𝒯**
- Γ *consists of some development-paths such that* ∀*γ* ∈ Γ ∀*t* ∈ **𝒯** ∀*s* ∈ **𝒯** *γ*_*𝒪*_(*t*) = *γ*_*𝒪*_(*s*), *and*

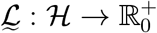

*where* ℋ = Dom 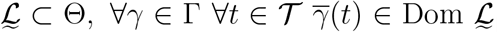, *and* 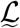 *satisfies the following properties:* *(es)* 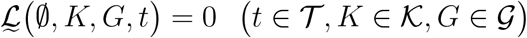 *(do) If* 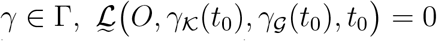, *t*_0_ *< t*_1_ *then* 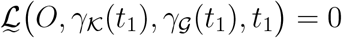 (*where* O = *γ*_𝒪_ (*t*_0_) = *γ*_𝒪_ (*t*_1_)). *(ss) If O*_1_ ⊂ *O*_2_ *then* 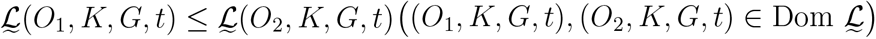.

*We call* 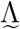 *a* ***life-level-system in the general context***. *If we want to emphasiz e that* **ℒ** *is a life-level function in the general context only then we use the notation:* 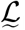.□

#### The basic version of the definition schema

Finally we provide the simplest version where there are no development-paths at all. I.e. neither the gorganism, nor the sub-environment, nor the supporting-group is due to change in time, in other words Γ can be considered empty.

##### Definition 2.1.20.

***In the less general context, life-level*** *is a function* **ℒ** *that is defined by the following system* Λ = ⟨⟨**𝒪**, ⊂⟩, ⟨**𝒦**, ⊂⟩, ***𝒢*, 𝒯**, Θ, **ℒ**⟩

*where* **𝒯** ⊂ ℝ *is closed and*

1. ⊂ *is a partial order on* ***𝒪***,

⊂ *is a partial order on* ***𝒦***,

- *there is a least element in* ***𝒪*** *that is denoted by* ∅,
- **𝒢** ⊂ *P (****𝒪****), and* ***𝒢*** *is closed for finite union and finite intersection*
- Θ ⊂ **𝒪** × **𝒦** × **𝒢** × **𝒯** *and*

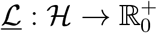

*where* ℋ = Dom **ℒ** ⊂ Θ *and* **ℒ** *satisfies the following properties:* *(do) If* **ℒ**(*O, K, G, t*_0_) = 0, *t*_0_ *< t*_1_ *then* **ℒ**(*O, K, G, t*_1_) = 0 ((*O, K, G, t*_0_), (*O, K, G, t*_1_) ∈ Dom **ℒ)**. *(ss) If O*_1_ ⊂ *O*_2_ *then* **ℒ**(*O*_1_, *K, G, t*) ≤ **ℒ**(*O*_2_, *K, G, t*) ((*O*_1_, *K, G, t*), (*O*_2_, *K, G, t*) ∈ Dom **ℒ)**.

*We call* Λ *a* ***life-level-system in the less general context***. *If we want to emphasize that* **ℒ** *is a life-level function in the less general context only, then we use the notation:* **ℒ**.

**Remark 2.1.21**. *One may use the (not too precise but understandable) notation that* (*O, K, G*) ∈ Dom **ℒ** *if there is a non-degenerate interval I* ⊂ **𝒯** *such that* {(*O, K, G, t*) : *t* ∈ *I*} ⊂ Dom **ℒ**.

##### 2.1.1 Deriving the order relations from pre-structures

In definition 2.1.10 we have a partial order relation for all *t* ∈ **𝒯** on both **𝒪**(*t*) and **𝒦**(*t*). In the simpler versions we have a partial order relation on both **𝒪** and **𝒦**. Here we will provide some more specific versions of the definition schemas, where we derive those partial order relations from some pre-structures. The pre-structures will consist of some basic elements from which the higher elements can be got as a composition i.e. a higher element is a subset of some basic elements.

In order to do so, we assume that for all *t* ∈ **𝒯** there are sets **𝒪**_*e*_(*t*), ***𝒦***_*e*_(*t*) such that ∀*t* ∈ **𝒯 𝒪**(*t*) ⊂ *P* (***𝒪***_*e*_(*t*)), ∀*t* ∈ **𝒯 𝒦**(*t*) ⊂ *P(****𝒦***_*e*_(*t*)). Then the required ordering relation ⊂ in 2.1.10 is the subset relation in **𝒪**(*t*) and **𝒦**(*t*) respectively. I.e. if *O*_1_, *O*_2_ ∈ **𝒪**(*t*), then *O*_1_ ⊂ *O*_2_ holds iff *O*_1_ is a subset of *O*_2_. Similarly for **𝒦**(*t*).

Let us rephrase our definitions based on that i.e. let us create more specific versions of the definition schemas.

###### Definition 2.1.22.

Λ_*sb*_ = ⟨**𝒪**_*e*_, ***𝒦***_*e*_, ***𝒪, 𝒦, 𝒢, 𝒯***, Θ⟩ *is called an* ***e-life-level-subbase-system*** *if* **𝒯** ⊂ ℝ *is closed*, ∀*t* ∈ **𝒯 𝒪**_*e*_(*t*), **𝒦**_*e*_(*t*), **𝒪**(*t*), **𝒦**(*t*), **𝒢**(*t*) *are sets such that*

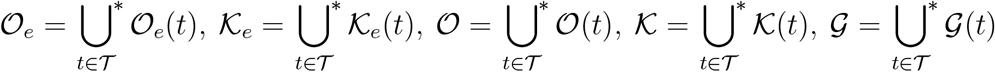

*and*

- ∀*t* ∈ **𝒯 𝒪**(*t*) ⊂ *P* (**𝒪**_*e*_(*t*)), ∀*t* ∈ **𝒯 𝒦**(*t*) ⊂ *P (****𝒦***_*e*_(*t*))
- ∀*t* ∈ **𝒯 𝒢**((*t*) ⊂ *P (****𝒪***(*t*)),
- ∀*t* ∈ **𝒯 𝒪**(*t*), **𝒦**(*t*), **𝒢**(*t*) *are all closed for finite union and finite inter-section*.

*Moreover*

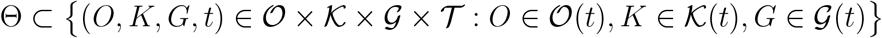

*holds*.

*Naming conventions:*

1. *An element of* ***𝒪***_*e*_(*t*) *is called* ***gorganism-particle*** *existing at time t*.
2. *An element of* ***𝒦***_*e*_(*t*) *is called* ***environment-particle*** *existing at time t*.

###### Definition 2.1.23.

Λ_*b*_ = ⟨**𝒪**_*e*_, **𝒦**_*e*_, **𝒪, 𝒦, 𝒢, 𝒯**, Θ, Γ⟩ *is called an* ***e-life-level-base-system*** *if* ⟨**𝒪**_*e*_, ***𝒦***_*e*_, ***𝒪, 𝒦, 𝒢, 𝒯***, Θ⟩ *is an e-life-level-subbase-system and* Γ *is a subset of the set of all development-paths*.

###### Definition 2.1.24.

*Let an e-life-level-base-system* Λ_*b*_ = ⟨**𝒪**_*e*_, ***𝒦***_*e*_, ***𝒪, 𝒦, 𝒢*, 𝒯**, Θ, Γ⟩ *be given. In the most general context* ***life-level*** *is a function* **ℒ** *that such that*

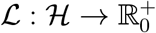

*where* ℋ = Dom **ℒ** ⊂ Θ *and* **ℒ** *satisfies properties (es), (do) and (ss)*.

*The system* Λ = ⟨**𝒪**_*e*_, ***𝒦***_*e*_, ***𝒪, 𝒦, 𝒢*, 𝒯**, Θ, Γ, **ℒ**⟩ *is called* ***e-life-level-system***.

In this manner, for the first type of simplification of the definition (definition 2.1.19) we need that ∀*t* ∈ ***𝒯*** ∀*s* ∈ ***𝒯 𝒪***_*e*_(*t*) = **𝒪**_*e*_(*s*), ***𝒦***_*e*_(*t*) = **𝒦**_*e*_(*s*). The definition can be rephrased according to that.

Similarly the corresponding version of definition 2.1.20 can be formalized. We are not going to add them here in full details.

### 2.2 Properties and basic facts

#### Definition 2.2.1.

*Let a life-level-system* Λ *be given and let* (*O, K, G, t*) ∈ Dom **ℒ**. *We say that the bio-bit* (*O, K, G, t*) *is* ***alive*** *(according to* Λ *or* **ℒ***), if* **ℒ**(*O, K, G, t*) *>* 0. *One can also say that* (*O, K, G*) *is alive at time t*.

#### Proposition 2.2.2.

*Let a life-level-system* Λ *be given and γ* ∈ Γ. *If property (do) in 2.1.10 holds, then* {*t* ∈ **𝒯** : **ℒ**_*γ*_(*t*) *>* 0} ⊂ **𝒯** *is convex in i.e. it is an interval*.

*Proof*. Let *t*^*′*^ ≤ *t*^*′′*^ ≤ *t*^*′′′*^ such that **ℒ**_*γ*_(*t*^*′*^) *>* 0 and **ℒ**_*γ*_(*t*^*′′′*^) *>* 0. Property (es) gives that *γ*_*𝒪*_(*t*^*′*^) =? ∅ and *γ*_*𝒪*_(*t*^*′′′*^) ≠ ∅, hence *γ*_*𝒪*_(*t*^*′′*^) ≠ ∅ because {*t* ∈ **𝒯** : *γ*_*𝒪*_(*t*)≠ ∅} is convex by the assumption on any *γ* ∈ Γ. Property (do) gives that **ℒ**_*γ*_(*t*^*′′*^) cannot be 0 since otherwise **ℒ**_*γ*_(*t*^*′′′*^) would be 0 as well.

#### Definition 2.2.3.

*Let a life-level-subbase-system* Λ_*sb*_ *be given. Let O*_1_, *O*_2_ ∈ **𝒪**. *Set O*_1_ *⥽ O*_2_ *if O*_1_ ⊂ *O*_2_ *and* ∄*O*_3_ ∈ **𝒪** *such that O*_1_ ⊂ *O*_3_ ⊂ *O*_2_.

Now we are going to categorize gorganisms by a kind of complexity, that is roughly if it contains smaller gorganisms or not.

#### Definition 2.2.4.

*Let a life-level-subbase-system* Λ_*sb*_ *be given. For O* ∈ **𝒪** *set*

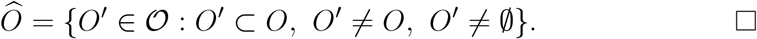

#### Definition 2.2.5.

*Let a life-level-subbase-system* Λ_*sb*_ *be given. A gorganism O* ∈ **𝒪** *is called* ***primitive*** *if* 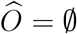.

#### Definition 2.2.6.

*Let a life-level-subbase-system* Λ_*sb*_ *be given. A gorganism O* ∈ **𝒪** *is called* ***group-type*** *if* 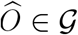.

**Remark 2.2.7**. *A group-type gorganism should not be mixed with a supporting-group. I.e. if O* ∈ **𝒪** *is group-type then it does not necessarily mean that* 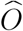 *is a supporting-group of some gorganism*.

**Remark 2.2.8**. *A gorganism is not either primitive or group-type, it can be none of those. Technically it is because that* ***𝒢*** *defines what a group is*.

We define the hierarchy-level for gorganisms recursively.

#### Definition 2.2.9.

*Let a life-level-subbase-system* Λ_*sb*_ *be given. If a gorganism O* ∈ ***𝒪*** *is primitive then set h*(*O*) = 1 *and we say that it has hierarchy-level 1. Suppose that we have already defined gorganisms with h*(*O*) ≤ *n* (*n* ∈ N). *If we have a gorganism O* ∈ **𝒪** *for which h*(*O*) *is not defined yet, then set h*(*O*) = *n* + 1 *if* ∀*O*^*′*^ ∈**𝒪** *O*^*′*^ ⊂ *O implies that h*(*O*^*′*^) *is defined and h*(*O*^*′*^) ≤ *n. We call h*(*O*) *the* ***hierarchy-level*** *of O*.

#### Proposition 2.2.10

*Let O* ∈ **𝒪**. *Then*

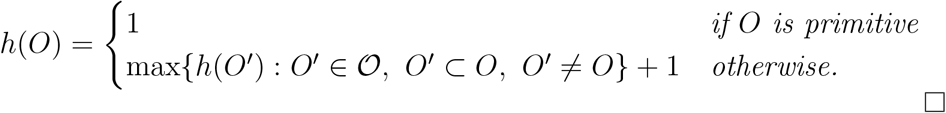

#### Definition 2.2.11.

*Let a life-level-system* Λ *be given. We call the life-level function* **ℒ *continuous*** *if* ∀*γ* ∈ Γ *the function* **ℒ**_*γ*_|_*𝒯*_ *is continuous where T* = {*t* ∈ **𝒯** : ∃*t*^*′*^ ∈ **𝒯** *such that t*^*′*^ ≤ *t*, **ℒ**_*γ*_(*t*^*′*^) *>* 0} ⊂ **𝒯**.

#### Proposition 2.2.12.

*In the less general context the life-level function* **ℒ** *is continuous if I* ⊂ **𝒯** *being a non-degenerate interval and* {(*O, K, G, t*) : *t* ∈ *I*} ⊂ Dom **ℒ** (*O* ∈ **𝒪**, *K* ∈ **𝒦**, *G* ∈ **𝒢**) *implies that the function t* 1→ **ℒ**(*O, K, G, t*) (*t* ∈ *I*) *is continuous*.

#### Proposition 2.2.13.

*Let a life-level-system* Λ *be given. Let* (*O*_1_, *K, G, t*) ∈ Dom **ℒ**, (*O*_2_, *K, G, t*) ∈ Dom **ℒ**. *Suppose that the supremum of O*_1_, *O*_2_ *exists and denote it by O*_1_ ∪ *O*_2_. *Then*

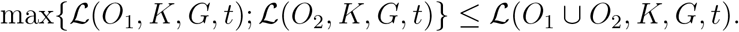

*Proof*. Obvious consequence of property (ss).

The life-level of a group is not lower than the life-level of one of its elements, more precisely:

#### Proposition 2.2.14.

*Let a life-level-system* Λ *be given. If O, G* ∈ **𝒪**(*t*), *O* ⊂ *G and* 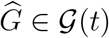 *then* **ℒ**(*O, K, G*^*′*^, *t*) ≤ **ℒ**(*G, K, G*^*′*^, *t*).

*Proof*. Obvious consequence of property (ss).

We are now going to formalize when we consider two life-level functions to be the same in the sense of measuring life-level.

#### Definition 2.2.15.

*Let a life-level-base-system* Λ_*b*_ *be given with two associated life-level functions* **ℒ**_1_ *and* **ℒ**_2_. *We say that the two life-level functions are* ***equivalent*** *if* Dom **ℒ**_1_ = Dom **ℒ**_2_ *and there is a strictly increasing function f* : 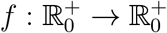 *such that*

1. *f* (*x*) = 0 ⇐⇒ *x* = 0 *and*
2. **ℒ**_2_(*b*) = *f* (**ℒ**_1_(*b*)) (*b* ∈ Dom **ℒ**_1_).

*In this case we also use the notation* **ℒ**_1_ ∼ **ℒ**_2_.

**Remark 2.2.16**. *Condition (1) can be weakened to f* (0) = 0 *since f* (*x*) = 0 ⇒ *x* = 0 *follows from the strictly increasing property of f*.

#### Proposition 2.2.17.

*The equivalence of life-level functions is an equivalence relation*.

*Proof*. The inverse of a strictly increasing function is strictly increasing. Also the composition of two strictly increasing function is strictly increasing.

#### Proposition 2.2.18.

*Let a life-level-base-system* Λ_*b*_ *be given with two associated life-level functions* **ℒ**_1_ *and* **ℒ**_2_. *Then* **ℒ**_1_, **ℒ**_2_ *are equivalent if and only if*

1. *they have the same domain*
2. *they assign 0 to exactly the same elements*
3. *if* **ℒ**_1_(*b*_1_) = **ℒ**_1_(*b*_2_) *then* **ℒ**_2_(*b*_1_) = **ℒ**_2_(*b*_2_)
4. *4) if* **ℒ**_1_(*b*_1_) *<* **ℒ**_1_(*b*_2_) *then* **ℒ**_2_(*b*_1_) *<* **ℒ**_2_(*b*_2_) *where b*_1_ = (*O*_1_, *K*_1_, *G*_1_, *t*_1_), *b*_2_ = (*O*_2_, *K*_2_, *G*_2_, *t*_2_), *b*_1_, *b*_2_ ∈ Dom **ℒ**_1_.

*Proof*. The necessity is obvious.

To prove the sufficiency, let *x* ∈ Ran **ℒ**_1_, say *x* = **ℒ**_1_(*b*) for some *b* ∈ Dom **ℒ**_1_. Then set *f* (*x*) = **ℒ**_2_(*b*). By property (3) *f* is well defined. Property gives that *f* (*x*) = 0 ⇐⇒*x* = 0, while property (4) yields that *f* is strictly increasing on its domain. Now one can extend *f* to the whole of 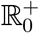 in the usual way.

#### Proposition 2.2.19.

*Let a life-level-base-system* Λ_*b*_ *be given with two associated life-level functions* **ℒ**_1_ *and* _2_**ℒ**. *Then* **ℒ**_1_, **ℒ**_2_ *are equivalent if and only if*

1) *they have the same domain*
2b) **ℒ**_1_(*b*_1_) ≤ **ℒ**_1_(*b*_2_) *iff* **ℒ**_2_(*b*_1_) ≤ **ℒ**_2_(*b*_2_) *where b*_1_ = (*O*_1_, *K*_1_, *G*_1_, *t*_1_), *b*_2_ = (*O*_2_, *K*_2_, *G*_2_, *t*_2_), *b*_1_, *b*_2_ ∈ Dom **ℒ**_1_. *Proof*. First let us note that (3) and (4) implies (2). Hence we have to show the equivalence of (3) (4) and (2b).

Set *b*_1_ = (*O*_1_, *K*_1_, *G*_1_, *t*_1_), *b*_2_ = (*O*_2_, *K*_2_, *G*_2_, *t*_2_), (*b*_1_, *b*_2_ ∈ Dom **ℒ**_1_).

Suppose first that (3) (4) hold and **ℒ**_1_(*b*_1_) ≤ **ℒ**_1_(*b*_2_). If **ℒ**_1_(*b*_1_) = **ℒ**_1_(*b*_2_) then apply (3); if **ℒ**_1_(*b*_1_) *<* **ℒ**_1_(*b*_2_) then apply (4). If **ℒ**_2_(*b*_1_) ≤ **ℒ**_2_(*b*_2_) then suppose that **ℒ**_1_(*b*_1_) *>* **ℒ**_1_(*b*_2_). By (3) we would get that **ℒ**_2_(*b*_1_) *>* **ℒ**_2_(*b*_2_) – a contradiction.

Suppose that (2b) holds and **ℒ**_1_(*b*_1_) = **ℒ**_1_(*b*_2_). Then **ℒ**_1_(*b*_1_) ≤ **ℒ**_1_(*b*_2_) and **ℒ**_1_(*b*_1_) ≥ **ℒ**_1_(*b*_2_) both hold, which yields that **ℒ**_2_(*b*_1_) = **ℒ**_2_(*b*_2_). The same holds for **ℒ**_2_. Hence we have showed (3).

If **ℒ**_1_(*b*_1_) *<* **ℒ**_1_(*b*_2_) then suppose that **ℒ**_2_(*b*_1_) ≥ **ℒ**_2_(*b*_2_). Then (2b) gives that **ℒ**_1_(*b*_1_) ≥ **ℒ**_1_(*b*_2_) – a contradiction.

#### Definition 2.2.20.

*Let a life-level-base-system* Λ_*b*_ *be given with two associated life-level functions* **ℒ**_1_ *and* **ℒ**_2_. *We say that the two life-level functions are* ***alive-equivalent*** *if* Dom **ℒ**_1_ = Dom **ℒ**_2_ *and*

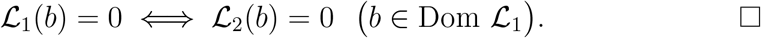

#### Proposition 2.2.21.

*By the notation of the previous definition, if* **ℒ**_1_ *and* **ℒ**_2_ *are equivalent, then they are also alive-equivalent*.

#### Proposition 2.2.22.

*By the notation of the previous definition*, **ℒ**_1_ *and* **ℒ**_2_ *are alive-equivalent if and only if* sgn(**ℒ**_1_) = sgn(**ℒ**_2_).

#### Definition 2.2.23.

*Let a life-level-base-system* Λ_*b*_ *be given with two associated life-level functions* **ℒ**_1_ *and* **ℒ**_2_. *We say that* **ℒ**_2_ *is* ***more permissive*** *than* **ℒ**_1_ *if* Dom **ℒ**_1_ = Dom **ℒ**_2_ *and*

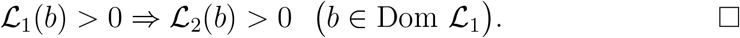

We can connect the values of a life-level function with addition however we think it is just an optional property of a life-level function.

#### Definition 2.2.24.

*Let a life-level-system* Λ *be given such that for all O*_1_, *O*_2_ ∈ **𝒪**(*t*) *the supremum of O*_1_, *O*_2_ *exists and belongs to* ***𝒪***(*t*) *(denoted by O*_1_ ∪ *O*_2_*). The associated life-level function* **ℒ** *is called* ***subadditive*** *if* (*O*_1_, *K, G, t*), (*O*_2_, *K, G, t*) ∈ Dom **ℒ** *implies that*

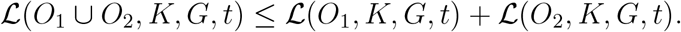

*Furthermore if the infimum of O*_1_, *O*_2_ *exists and belongs to* ***𝒪***(*t*) *(denoted by O*_1_ ∩ *O*_2_*), then* **ℒ** *is called* ***additive*** *if* **ℒ**(*O*_1_ ∪ *O*_2_, *K, G, t*) = **ℒ**(*O*_1_, *K, G, t*) + **ℒ**(*O*_2_, *K, G, t*) − **ℒ**(*O*_1_ ∩ *O*_2_, *K, G, t*).

#### Proposition 2.2.25.

*Let a life-level-system* Λ *be given such that O*_1_∪*O*_2_ *and O*_1_∩*O*_2_ *exist for all O*_1_, *O*_2_ ∈ **𝒪**(*t*). *Then* **ℒ** *being subadditive (additive) is equivalent with that the function O ⟼* **ℒ**(*O, K, G, t*) *is subadditive (additive) for fixed t* ∈ **𝒯**, *K* ∈ **𝒦**(*t*), *G* ∈ **𝒢**(*t*).

#### 2.2.1 Properties of Γ

Here we enumerate some properties of the set of development-paths Γ which we will refer to in the sequel.

First we define when a bigger gorganism always has a development-path synchronized with the development-path of the smaller gorganism i.e. on that development-paths they stay on the same sub-environment and supporting-group.

##### Definition 2.2.26.

*Let a life-level-base-system* Λ_*b*_ *be given. We call* Λ_*b*_ **Γ*-above-synchronized*** *if b, b*^*′*^ ∈ Θ, *b* ⊆_*𝒪*_ *b*^*′*^, *γ* ∈ Γ, 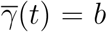 *implies that* ∃*γ*^*′*^ ∈ Γ *such that* 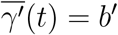 *and* 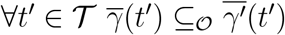.

*We call* Λ_*b*_ **Γ*-above-synchronized-forward*** *if the last condition is changed to* 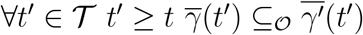.

*We call* Λ_*b*_ **Γ*-above-synchronized-backward*** *if the last condition is changed to* 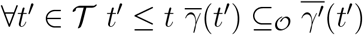.

When we swap the bigger and smaller gorganism, then we get the notion of Γ-synchronized-below.

##### Definition 2.2.27.

*Let a life-level-base-system* Λ_*b*_ *be given. We call* Λ_*b*_ **Γ*-below-synchronized*** *if b, b*^*′*^ ∈ Θ, *b* ⊆_*𝒪*_ *b*^*′*^, *γ*^*′*^ ∈ Γ, 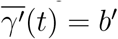 *implies that* ∃*γ* ∈ Γ *such that* 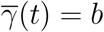 *and* 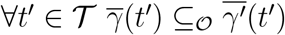.

*We call* Λ_*b*_ **Γ*-below-synchronized-forward*, Γ*-below-synchronized-backward*** *if t*^*′*^ ≥ *t, t*^*′*^ ≤ *t added to the last condition respectively*.

We define the notion that if two development-paths intersects, then we can combine the before and after parts to get a new development-path.

##### Definition 2.2.28.

*Let a life-level-base-system* Λ *be given. Then* Γ *is called* ***closed*** *if γ, γ*^*′*^ ∈ Γ, *γ*(*t*) = *γ*^*′*^(*t*) *implies that γ*|_(*−∞,t*]_ ∪ *γ*^*′*^|_[*t*,+*∞*)_ ∈ Γ.

We define the notion that if a gorganism dies, then it vanishes immediately.

##### Definition 2.2.29.

*Let a life-level-system* Λ *be given. We call* Λ_*b*_ **Γ*-vanishing*** *if γ* ∈ Γ **ℒ**(*γ*_*𝒪*_(*t*_0_), *γ*_*𝒦*_(*t*_0_), *γ*_*𝒢*_(*t*_0_), *t*_0_) = 0, *t*_0_ *< t*_1_ *implies that γ*_*𝒪*_(*t*_1_) = ∅.

In some simpler models, we do want development-paths, but each bio-bit can have only one such.

##### Definition 2.2.30.

*Let a life-level-base-system* Λ_*b*_ *be given. We call* Λ_*b*_ **Γ*-unique*** *if for every b* ∈ Θ *there is a unique γ* ∈ Γ *such that* 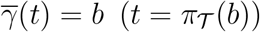.

We set a notation for all development-paths that start from a given bio-bit.

##### Definition 2.2.31.

*Let a life-level-base-system* Λ_*b*_ *be given and let b* = (*O, K, G, t*) *be an admissible bio-bit. Then set*

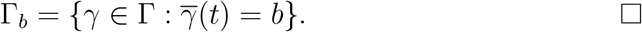

### 2.3 Some explanation

In this subsection and the following few, we are going to support the understanding of the many kinds of definitions, notions and the terminology. Here we do not provide examples (or just rarely). They come in section 3.

Again, we emphasize that the theory is based on some primitive notions which remain undefined here too (see the beginning of section 2). The main ones are “*organism*” and “*generalized organism*”. They can be anything whose life-level is worth examining.

**Remark 2.3.1**. *A given life-level function* **ℒ** *(more precisely a life-level-system* Λ*) belongs to a specific research project. In other words every research project can have its own life-level function. The project has to define its own*

- *sets:* ***𝒪, 𝒦, 𝒢***,
- *partial orders on* ***𝒪, 𝒦*** *(or* ***𝒪***_*e*_, ***𝒦***_*e*_, *if the partial orders are derived from those)*,
- **𝒯** *if it is special*,
- *the set of admissible bio-bits* Θ,
- *the set of possible development-paths* Γ *(in the more complex cases)*,
- *and finally* **ℒ**.

*In the sequel we will handle and understand* **ℒ** *(and* Λ*) in this sense, without emphasizing that those sets and function(s) belong to a given research project and they do not exist a priori*.

**Remark 2.3.2**. *0 has a special highlighted role in the definition, namely* 0 *is used for expressing “not-alive”. It is the only such value. The special role of 0 exists because we are not going to distinguish between non-living levels*.

**Remark 2.3.3. 𝒪**_*e*_ *is the set of the gorganism-particles that may be too simple to be investigated but all gorganisms that are due to be investigated are somehow built from those. We of course allow that gorganism-particles can be among the elements of* ***𝒪*** *but the formalism gives that it is possible in the following form: if O* ∈ **𝒪**_*e*_ *then* {*O*} ∈ **𝒪** *is possible. In such a case we may identify* {*O*} *with O*.

**Remark 2.3.4. 𝒦**_*e*_ *is the set of the environment-particles that may be too simple to be used in any investigation but all sub-environments that are due to become part of an investigation are somehow built from those. Similarly we allow that environment-particles can be among the elements of* ***𝒪*** *but in the following form: if K* ∈ ***𝒦***_*e*_ *then {K}* ∈ ***𝒦*** *is possible. In such a case we may identify {K} with K. An environment-particle is strictly “non-living”*.

**Remark 2.3.5. *𝒪*** *is the set of all gorganisms (regarding a given research project) that are due to be investigated regarding in the sense of life-level*.

**Remark 2.3.6**. *It is important to emphasize that* ***𝒪*** *is not necessarily the full set of all gorganisms of the Earth; an application can focus on a (very) small ecosystem only, separated from all other living beings, and can create its life-level model for that small system only. The application does not need to make efforts to synchronize its model to other ecosystems outside its own one. This theory provides an option to have partial life-level models, i.e. separate models for distinct ecosystems of the Earth*.

**Remark 2.3.7**. *We remark again that “gorganism” is slightly the synonym of organism however it may be much much more. It can denote smaller, less complex, or even bigger, much more complex beings than the word “organism” usually denotes*.

**Remark 2.3.8**. *The partial order* ⊂ *on* ***𝒪*** *is supposed to express which gorganism is “part” of some other gorganism. Usually “part” can be meant that it is part physically, but it is not necessary, an application can define it according to its need and purpose. I.e. “part” expresses some kind of biological smallness (better to say partial ordering) between gorganisms that has to be in accordance with property (ss) in definition 2.1.10*.

*It is important to emphasize that an application does not have to add all possible components of a gorganism to its life-level-system. E.g. it may add none at all, or just a few selected ones*.

*Similarly, the partial order* ⊂ *on* ***𝒪*** *is supposed to express which sub-environment is “part” of some other sub-environment. All previous comments can be applied here too*.

*It may be slightly confusing that the partial order is denoted by* ⊂ *instead of* ≤, *however we think that it more intuitively expresses what that is for*.

*For technical reasons we need a minimum element in* ***𝒪***. *Again, It may be slightly confusing that it is denoted by* ∅, *however we have deliberately chosen that*.

*We remark that if O*_1_ ⊂ *O*_2_ *holds, then necessarily O*_1_, *O*_2_ *belong to an* ***𝒪***(*t*) *for the same t, because that is how we defined the relation* ⊂ *and the sets* ***𝒪***(*t*) *are disjoint for different t*.

**Remark 2.3.9**. *One may argue against property (ss) in definition 2.1.10, saying that when an organism dies, then some of its organs may remain alive for some time after death. In the model built here, another approach is followed, namely, the organism is partially alive when it has some components that are still partially alive. Moreover it is at least as much alive as its components are alive*.

*Or one may say that property (ss) implies: a gorganism is not alive, then none of its components can be alive. But the opposite implication is not part of definition 2.1.10 deliberately (or it cannot be derived). We wanted to keep the option that in a model, none of the components are alive, while the gorganism is alive*.

**Remark 2.3.10**. *Regarding property (es) in definition 2.1.10, the only reason why we formulate this condition for the empty set in* **𝒪** (*t*), *is that we need to have development-paths defined on the whole* **𝒯**. *I.e. it is a kind of technical condition, however it is obviously a natural requirement*.

**Remark 2.3.11**. *Property (do) in definition 2.1.10 obviously expresses that if an gorganism died at some time point, then it remains dead, never gets alive again*.

**Remark 2.3.12. 𝒦** *is the set of some meaningful parts of the environment. They are called sub-environments that may have effect on the life-level of a gorganism from* **𝒪**. *An element of* **𝒦** *is strictly* “*non-living"*.

**Remark 2.3.13. 𝒢** *is the set of some groups of gorganisms from* **𝒪** *i.e. an element of* **𝒢** *is a meaningful subset of* **𝒪** *that may have influence on the life-level of a gorganism from* **𝒪**, *or it may be important from some other aspect too*.

**Remark 2.3.14. 𝒢** *contains the supporting-groups as well. Groups that might have effect on the life-level of gorganisms*.

*It should not be mixed with the notion of a social group*, “*supporting-group” is a far more general concept*.

*It is important to mention that despite of the usual meaning of the adjective* “*supporting", it is also possible that the group supports the gorganism negatively i.e. it decreases its life-level*.

“*Supporting” is not a synonym for* “*helping through contact". It can happen that an element of the supporting group does not have any close contact with the gorganism, or even no contact at all, no closer, no further. See example 3.1.5 item 12*.

**Remark 2.3.15**. *The notion of* “*supporting-group” can be considered as the* “*living” (nearby) sub-environment of an organism. In that sense, we split the environment of an organism into two categories: the* “*living” and and* “*non-living” parts of the broader environment*.

**Remark 2.3.16**. *In the definition 2.1.10 we have not assumed that O* ∈ *G when* (*O, K, G, t*) ∈ Dom ℒ. *It is because we wanted to keep the possibility that the supporting-group does not contain the gorganism, but still has non neglectable effect on the life-level of the gorganism. However we think that in many applications O* ∈ *G will be assumed*.

*Moreover in some cases it can be especially useful if the supporting-group of a gorganism contains the gorganism itself. See e.g. subsection 7.2*.

**Remark 2.3.17**. *The supporting-group of a group-type gorganism usually also contains group-type gorganisms, however it is not necessary. See example 3.1.5 item 10*.

**Remark 2.3.18**. *The life-level of a group G can be investigated too i.e. a group can be in* **𝒪**, *more precisely it is allowed that* 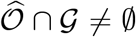 *where* 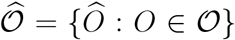. *In this case, such G might have two roles in the model: its life-level can be asked (together with the other parameters) or it can serve as a supporting-group for some gorganism*.

**Remark 2.3.19**. *We allow that a supporting-group G can contain an only element, i.e. G* = {*O*} *for some O* ∈ **𝒪**.

**Remark 2.3.20**. *Among the elements of* **𝒢** *the subset* ⊂ *relation gives a partial order on* **𝒢**.

**Remark 2.3.21. 𝒯** *is the time or the timeline where we investigate life-level. Usually it can be identified with some (finite or infinite) interval of* ℝ *however it is not necessary*.

*We also allow that* **𝒯** *can be discrete. For example if* **𝒯** = ℕ *then we* “*measure” life-level in e.g. every second or every month or every millennium etc*.

*If* **𝒯** *is discrete then* ℒ *is always continuous since on discrete sets any function is continuous*.

**Remark 2.3.22**. *One may argue here (2.3.21) that it does not make sense measuring life-level on a millennium time scale since organisms do not live that long. It is a misunderstanding since in this paper organism is understood much more generally. An organism can be e.g. a small but whole biosphere of a forest, or all members of a given species, or even the whole biosphere of the Earth, etc. That is also why we prefer to use the term* “*generalized organism” instead*.

**Remark 2.3.23**. *When* **𝒪, 𝒦, 𝒢, 𝒯** *are defined, then the next step is that the project has to specify which objects* (*O, K, G, t*) (*O* ∈ **𝒪** (*t*), *K* ∈ **𝒦** (*t*), *G* ∈ **𝒢** (*t*)) *are worth investigating, i.e. it has to define the set of admissible bio-bits* Θ. *Of course the domain of a life-level function can be smaller than that, e.g. because many life-level functions are maintained with different domains*.

**Remark 2.3.24**. *In every moment we can ask the life-level of a gorganism that exists in that moment and is in a currently existing sub-environment and belongs to a currently existing supporting-group*.

**Remark 2.3.25**. *A gorganism of* **𝒪** *is an existing generalized organism at a given point of time. E.g. a rabbit that really exists somewhere. It can also be an existing groups of rabbits. Or all rabbits in a forest or on the whole planet at a given point of time. But a gorganism must not be just* “*rabbit” as there is no such <u>existing</u> thing as* “*rabbit”. In other words*, ***in the original concept we do not assign values to species***. *It is very important. (cf. subsection 8.3 where we assign values to groups and also section 9 where we derive life-level values for species)*

*Instead, one may ask the following reasonable questions for example:*

1. *What is the minimum of life-levels of all rabbits live in this small countryside now?*
2. *What was the maximum of life-levels of all rabbits live on the whole planet at 20.06.2019?*
3. *What was the average of life-levels of all rabbits live on the whole planet in the 19th century? (see also subsection 8.2)*
4. *Let us compare the following two sets: the (multi-)set of life-levels of all rabbits and the (multi-)set of life-levels of all foxes live in this small countryside now. (see also subsection 8.3)*

**Remark 2.3.26**. *Normally life-level cannot be investigated without the sub-environment and/or without the supporting-group as they may have remarkable effect on the result. Without considering those, our life-level result may be even false or at least it may be very far from a value that somehow more reasonably reflects* “*reality"*.

**Remark 2.3.27**. *In the model we allow that* ∅ ∈ **𝒦** *and* ∅ ∈ **𝒢**. *It is because we wanted to keep the possibility to have a simpler model where no sub-environment and/or no supporting-group is taken into account. As we mentioned before, it is not recommended, however the possibility is built in the model. Actually in very rare situations, the supporting-group may not exist naturally i.e. it can be empty (see example 3.1.5 item 13), but the sub-environment always exists, but can be excluded from the investigation if one wishes so*.

**Remark 2.3.28**. *How we determine the life-level of a given bio-bit is very much dependent on our knowledge. The more information we can use, the better values we can provide*.

*In that sense, but only in that sense, one may also say that life-level exists a priori and our aim is to get the best approximation that we can. And the more we know, the better approximation we can produce*.

**Remark 2.3.29**. *The natural expectation is not built in the definition 2.1.10 that similar gorganisms under similar circumstances should have similar life-level values. E.g. if we have two members of the same species living at the same area, under very similar circumstances, being at the same age, having the same sex, etc*., *then one would expect the same life-level values. For formalizing this expectation see 5.2.14*.

**Remark 2.3.30**. *According to definition 2.2.15, we may say that among the life-values, the order relations count only. The actual value of* ℒ *does not matter in the sense that if we move every value upwards (except 0) such that the* = *and < relations are kept, then the new life-level function will express exactly the same thing*.

*Therefore we may say roughly that two life-level functions* ℒ_1_, ℒ_2_ *are different if at least one of the following holds:*

- ∃ (*O, K, G, t*) *such that* ℒ_1_(*O, K, G, t*) = 0 *and* ℒ_2_(*O, K, G, t*) ≠ 0 *or vice versa*,
- ∃(*O*_1_, *K*_1_, *G*_1_, *t*_1_) ∃(*O*_2_, *K*_2_, *G*_2_, *t*_2_) *such that* ℒ_1_(*O*_1_, *K*_1_, *G*_1_, *t*_1_) = ℒ_1_(*O*_2_, *K*_2_, *G*_2_, *t*_2_) *and* ℒ_2_(*O*_1_, *K*_1_, *G*_1_, *t*_1_) ≠ ℒ_2_(*O*_2_, *K*_2_, *G*_2_, *t*_2_) *or vice versa*,
- ∃(*O*_1_, *K*_1_, *G*_1_, *t*_1_) ∃(*O*_2_, *K*_2_, *G*_2_, *t*_2_) *such that* ℒ_1_(*O*_1_, *K*_1_, *G*_1_, *t*_1_) *<* ℒ_1_(*O*_2_, *K*_2_, *G*_2_, *t*_2_) *and* ℒ_2_(*O*_1_, *K*_1_, *G*_1_, *t*_1_) *>* ℒ_2_(*O*_2_, *K*_2_, *G*_2_, *t*_2_) *or vice versa*.

**Remark 2.3.31**. *It is important to mention that property (ss) in definition 2.1.10 may not be true for elements of* **𝒦** *or* **𝒢** *(see examples 3.2.31 and 3.2.32)*.

**Remark 2.3.32**. *In definition 2.1.10 and the two simpler version too, we require three properties (es), (do) and (ss) from* ℒ *to be satisfied. The first one, (es) is a technical one, the other two, (do) and (ss) express meaningful requirements. In some cases, in some applications one can omit some of those properties and still can have a fairly applicable model. See e.g. subsection 7.2*.

**Remark 2.3.33**. *We are now going to give some insight why we build the following system into our model:* ∀*t* ∈ **𝒯 𝒪** (*t*), **𝒦** (*t*), **𝒢** (*t*) *are sets such that* 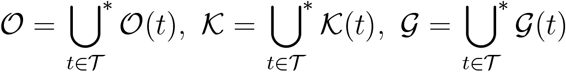 *(similarly for* **𝒪**_*e*_(*t*), **𝒦**_*e*_(*t*), *if needed). Why do we have time here? What does this formulation express? The short answer is that in every moment we have a reality. Every moment is different, more precisely, reality is different in every moment. We have* “*slices” of reality by time. The difficulty arises when we want to* “*connect” an* “*object” through time, i.e. when we want a kind of identification that says that this is more or less the same* “*object” all the way along that time interval. Or we do want to consider the change of the* “*object", but we want to follow its development-path in time. And in this case too, we need a kind of identification through time*.

*In concrete cases, the human perception may provide us an identification that may be more or less the same for everyone/every research project, however this is not always the case, and better to define and handle this phenomena precisely*.

*Therefore e.g*. **𝒪** (*t*) *contains all gorganisms that we see at time t, and* **𝒪** *is the set of all such slices. Similar is true for sub-environments and* **𝒦** (*t*), **𝒦**, *and for supporting-groups and* **𝒢** (*t*), **𝒢**.

We are now going to analyze the concept of development-path. On further analysis see subsection 2.4.

**Remark 2.3.34**. *Regarding a gorganism, a development-path is about to describe one of the followings:*

1. *How a generalized organism evolves through different life-forms (see for example 3.4.3 item 4)*.
2. *It may also describe how its* “*body” develops/changes significantly i.e. how the gorganism itself changes* “*inside” without taking a new life-form*.
3. *Or both at the same time*.

**Remark 2.3.35**. *Regarding sub-environment and/or supporting-group, a development-path may have following purposes:*

1. *It may describe how a gorganism changes its sub-environment and/or its supporting-group. It is about the* “*decisions” of the gorganism (see example 3.2.33)*.
2. *It may describe how the associated sub-environment and/or supporting-group develops in time. It is the* “*own business” of the sub-environment / supporting-group, i.e. independent from the gorganism (see example 3.2.34)*.
3. *Or both at the same time*.

**Remark 2.3.36**. *Consider a Γ* ∈ Γ. *Then Γ*_*O*_(*t*) *is the development-path of a gorganism, while Γ*_*G*_(*t*) *is the development-path of its supporting-group. One can have another Γ*^*′*^ ∈ Γ *such that* 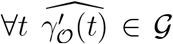 *i.e*. 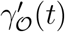 *is group-type. It is still the development-path of a gorganism (that may serve as a supporting-group for another gorganism as well). Of course in a special case, Γ*_*G*_(*t*) *and* 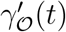 *can be either identical on the whole timeline or on a some short interval or just at a single point in time, as Γ and Γ*^*′*^ *describe very different phenomena*.

**Remark 2.3.37**. *The concept of development-path has one more purpose and it is to describe the possible <u>future</u> outcomes how a bio-bit can evolve. One can also use a probability distribution on the set of all development-paths. More precisely one may assign probability values to subsets of* Γ *and use it in various calculation (see e.g. subsection 4.4.2)*.

*As always in science, we try to predict the future and apply probability distributions using our knowledge at the present that is based on the past (and the present)*.

**Remark 2.3.38**. *Of course one cannot expect that the development-path of an initially group-type gorganism produces group-type gorganisms all along. And vice versa, the development-path of a non group-type gorganism can produce group-type gorganisms at some point in time*.

**Remark 2.3.39**. *Regarding the life-level-system in the less general context (definition 2.1.20) and also in the general context (definition 2.1.19), one may ask that if nothing is due to change, neither the gorganism, nor the sub-environment, nor the supporting-group, then why would the life-level change at all? In these simpler models the mapping between the model and reality is different (than in the most complex version). Actually the real gorganism (sub-environment, supporting-group) is changing slowly, but we do not want to represent it in the model. In other words we map the real world gorganism to the same gorganism in the model in a longer time interval, despite of the fact that it may be significantly different at the beginning and at the end of the interval. However, the change is so slow that we decide to keep the identification*.

*A simple example can be if we examine a man during 50 years of his life and we identify him to the same gorganism during all that time period*.

*Similar arguments may regard for the sub-environment and the supporting-group in the applications of definition 2.1.20 and 2.1.19*.

### 2.4 Life-level through lifespan

Here we are going to explain why the most complex definition schema has the right to exist, what it is for, and how to apply it. The main difference between the simplest and the most complex schema is the existence of development-paths. Hence it is about to explain why the notion of development-path is essential in complex applications.

It can be a general requirement to examine the life-level of a given gorganism through of its lifetime (or for some longer period of its lifetime). At first glance this requirement itself seems to be straightforward however it is not.

We have to be careful since the gorganism cannot be intact, it is constantly changing, hence it may be becoming to a different gorganism. So is the sub-environment where we investigate the gorganism. And so is the supporting-group the gorganism belongs to. All of those are changing in time.

Thus in a demanding and fairly sophisticated model, it is not possible to assign a life-value to a fixed gorganism in a fixed sub-environment in a fix supporting-group in a longer period of time. This simplification may be possible if we narrow the time for a sufficiently small period where the gorganism, the sub-environment and the supporting-group are more or less the same and intact. In longer periods we need a more sophisticated manner. Such changes that we are going to describe may not be trivial in the sense that at a given point of time our gorganism can split into two (or more) similar gorganisms e.g. cell division. And vice versa two gorganisms can form a new gorganism e.g. at some strong symbiotic relation (e.g. obligatory symbiosis) where after some time one dies without the other. Similar difficulties may arise for supporting-groups or even for sub-environments. Because supporting-groups / sub-environments can also change dramatically in time or can split into several pieces or some separate pieces may merge into one.

In such cases we have to choose how we proceed on the timeline, which gorganisms, sub-environments, supporting-groups belong together in the sense that they express one single way of development.

In order to handle that, we need functions for **𝒪, 𝒦, 𝒢** that describe the changes of gorganisms, sub-environments and supporting-groups respectively

i.e. we need functions of the following type.

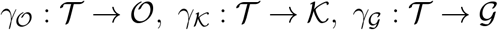

or simply we can write

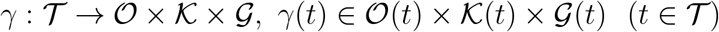

with the notation *Γ*_𝒪_ = *π*_𝒪_ º*Γ, Γ*_**𝒦**_ = *π*_**𝒦**_ º*Γ, Γ*_**𝒢**_ = *π*_**𝒢**_ º*Γ* as in definition 2.1.10. One single *Γ* is supposed to describe the lifespan of a gorganism, the lifespan of its associated sub-environment and the lifespan of its associated supporting-group, all at the same time.

Then for a given development-path *Γ*, we are able to examine the following function: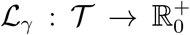, ℒ_*Γ*_(*t*) = ℒ (*Γ*_**𝒪**_ (*t*), *Γ*_**𝒦**_ (*t*), *Γ*_**𝒢**_ (*t*), *t*. It is going to describe the change of life-level in time on the development-path of our bio-bit.

Therefore we enrich the model by adding such development-paths to it and the set of such functions is denoted by Γ.

### 2.5 Comparison of the original life concept and the current model

We are going to summarize here in what contexts this definition schema or better to say a ready-made definition provides different viewpoints than the usual life concept. Again, we emphasize that this is not an assessment in any way, since we are talking about two models with more or less different aims.

The only purpose of this comparison is that one could see more clearly how these two models work.

First, let us gather briefly the key points of the **original life concept**.

1. Something is either alive or dead (no third option).
2. Mainly “*abstract* “ organisms are examined such as species. More precisely, “*perfect* “ representatives of a species are examined and they are identified with the species.
3. Only the “mainstream” organisms are examined from a species, rare naturally existing special cases are not. Individual organisms are not examined.
4. Only naturally existing organisms are examined without any artificial modifications.
5. Through examination the organism is completely detached from its surrounding and its fellow organisms.
6. The examination takes into account the information in the present only.
7. In some investigations (not all!) it is assumed that there has to be one final definition that is applicable everywhere and in every situation (of course it is meant that one definition with possible equivalent forms).

In comparison, the **current model** presented in this paper provides the following.

1. Life-level is not digital (where digital means: either alive or dead). It is worth using a scale that is infinite both directions: upwards and between two values. Let us emphasize that it is two additions, not one. On one hand, it is a refinement, but on the other hand, the model also allows arbitrarily large aliveness values.
2. “*Abstract*” organisms are not considered. Instead, only individual and existing ones examined. Also, rare, not usual natural cases can be examined.
3. The targets for investigation regarding life-level are not restricted to usual “*bodies*”. Instead, one can assign life-levels to a great variety of things, even much more simpler and/or much more complex ones than usually considered. Many different parts of a usual “*body*” can be examined in several layers as well.
4. Artificially modified or (partially) artificially created organisms can be examined.
5. This model takes into account the affect of the surrounding environment: Life-level does depend on the sub-environment.
6. Life-level may depend on the group to which the organism is somehow connected and/or belong.
7. The life-level of a given organism may change as time goes by. Either significantly or just slightly, the life-level time graph is an important addition.
8. The development-paths are about to describe how an organism evolves through different life-forms and connects that kind of change to life-level. And also a development-path describes how the sub-environment and/or the supporting-group may change and/or develop in time and how that affects life-level.
9. There is no final life-level model that would be applicable everywhere and in every situation, i.e. many such functions can exist. It is valid in two senses. Firstly, a life-level model may be created just for a given ecosystem, in other words, partial life-level models may cover the whole biosphere, and only one may not. Secondly, for the same ecosystem, one can have different life-level models based on e.g. different principles.

**Remark 2.5.1**. *At first glance one might think that the current model is a generalization of the usual life concept, i.e. it may contain the original one as a special case, since set K* = ∅, *G* = ∅, Ran ℒ = {0, 1} *and make* ℒ *time independent. However, it is not exactly true because the investigated objects are different, species vs. individual (generalized) organisms*.

Here we make a slight additional remark that there is one attribute that may provide enrichment to the original notion that may be worth considering to build into the original notion as well. It is to switching to a broader scale i.e. considering the option to use many different levels between “*not alive*” and “*fully alive*”, in other words use a scale that quantifies “*how much alive*” an organism is.

### 2.6 Some remarks on usage

Here we enumerate some remarks, some light suggestions on usage of this model. Please consider them as possible options on usage and nothing more.

**Remark 2.6.1**. *Consecutive integers could be used as the order of magnitudes on the life-level scale. For example 4 may express a significant change in life-level comparing to just 3 i.e. there is some <u>strong qualitative jump</u> between 3 and 4. In this case the numbers between 3 and 4 serves as a gradual transition between those two life-level layers*.

*In this sense the model may work similarly to the Richter scale for earth-quakes*.

**Remark 2.6.2**. *One may prefer logarithmic scale instead, similarly to measuring our sensation (which is proportional to the logarithm of the actual intensity measured with an accurate instrument (Weber–Fechner law)). Therefore e.g. the powers of* 10 *may serve as the order of magnitudes on the life-level scale*.

**Remark 2.6.3**. *One may prefer to use either finite decimals or rational numbers as the range of one’s life-level function. Rational numbers can be handy in the case when a kind of division-like algebraic operation also plays a role in one’s model*.

## 3 Examples

In this section we will provide many many examples in that belief that they will support the understanding of the main notions of this paper and also the way how the real world may be projected into this model.

Here we will regularly mention cases (examples) where we will deliberately not say anything about the possible values of life-level of the bio-bits in question. It is because we think it is not self-evident how to define it, and better to leave the question open and leave the decision to the actual research project because different viewpoints may result different values for life-level. However we think that just mentioning some possible order relations among the bio-bits regarding life-level will express sufficient information to grasp how a corresponding model may work in those cases. Of course, the mentioned order relations will be nothing more than reasonable preferences that are natural in some sense, but other viewpoints may result different order.

Likewise, we will just formulate questions without even sketching the answer. Again, we hope that the question itself helps understanding of the usage of the definition schemas.

In sections 4 and 7 we will present several models that accomplish the requirements of the definition schemas. Here, **in this section we are not going to apply those models** i.e. we will not discuss the applications of them to the current examples in details. At the moment our aim is not that. It is just to realize the problems one may face and outline some possible basic ordering relations. In this sense, this section can be considered as a natural continuation of subsection 1.2.1, because we describe the examples presented there in more detail and we add many more new examples as well.

One more reason that we do not give precise life-level values in the examples is that in some cases not only one of our models (from sections 4 and 7) can be applied. We suggest that after studying sections 4 and 7, the reader should reread this section and try to apply some of those models for the examples here, however we will analyze a few of those there.

We divide this section into smaller subsections from building blocks till lifespan examples.

### 3.1 On building blocks

In this subsection we provide examples for the building blocks of the definition schema. In each cases we are also going to provide many possible options how sets can be defined and chosen. This subsection is just about the building blocks, hence no life-level analysis happens here.

#### Gorganisms

##### Example 3.1.1.

*When an e-life-level-system is built, then the set of gorganism-particles* **𝒪**_*e*_ *contains the building blocks for the set of gorganisms* **𝒪**. *Certain subsets of* **𝒪**_*e*_ *may be the elements of* **𝒪**. *A few examples:*

1. **𝒪**_*e*_ *can contain all cells of all organisms living on a specific area*.
2. *Continuing item 1 we can go even down and* **𝒪**_*e*_ *can be all cell organelles of certain cells*.
3. *Or even down:* **𝒪**_*e*_ *can consist of all macro-molecules that build certain cells*.
4. *We can go upwards and* **𝒪**_*e*_ *can be all organisms living on a specific area*.
5. *If we are about to investigate e.g. birds then* **𝒪**_*e*_ *should also contain beaks, claws, feathers, bones, etc. that alone are definitely not alive but are essential components of any birds. We never examine any of those with respect to life-level (or even life), but some of them are* “*mandatory” building blocks of such living creatures. Mandatory in the sense that without such component the gorganism can hardly survive or expressing more strictly, we would never investigate the life-level of a bird without such*.

##### Example 3.1.2.

*The set* **𝒪** *contains all gorganisms that we are interested in regarding life-level (subsets of* **𝒪**_*e*_, *if an e-life-level-system is built). A few examples:*

1. **𝒪** *can consists of cells / tissues / organs / organ systems (just one layer or more or all). We can e.g. ask how much this cell is alive in this environment among those other cells, in that tissue etc. We can ask the same for a given organ. Etc*.
2. **𝒪** *can consists of all organisms living on a specific area. Or just some single organisms belonging to a given species*.
3. **𝒪** *can consists of all social groups of some animals living on a specific area in a given period of time*.
4. *Let the elements of* **𝒪** *be groups of organisms and one such group contains all organisms belonging to a species and we take into account all species of a given genus living on a specific area in a given period of time*.
5. *We can go on similarly for e.g. order / class / domain etc. (any hierarchical taxonomic ranks)*.
6. *Continuing the previous four points, we can also mix the different layers, e.g*. **𝒪** *can consists of some individual organisms, some social groups and also all members of another species, all at the same time*.
7. **𝒪** *can contain the whole biosphere of Earth in a given period of time as one gorganism (cf. [16])*.
8. *Continuing example 3.1.1 item 4, in this case* **𝒪** *would consist of groups of organisms*.
9. *Siphonophore is a kind of colonial organism, that appears to be an individual organism, but it is not. Zooids are the multicellular units that build the colonies, and they are genetically identical, because all developed from the same bud by fission at a later phase of development. Zooids are also functionally specialized for e.g. reproduction, digestion, floating, etc*. *When we examine some siphonophores that situated far from each other, then we can put the colonies into* **𝒪**, *or the different functional sub-colonies, or just the zooid components, or all of them*.
10. *Suppose that at some future point in time we will find organisms outside Earth (whatever the word* “*organism” means here). Then* **𝒪** *can contain the organisms of several biospheres of several planets, or even bigger: all organisms of a galaxy. E.g. we can also assign life-level values to the whole biospheres of different planets, we mean that one value for each biosphere (planet), and we can compare them*.

**Remark 3.1.3**. *In an e-life-level-system when we use* **𝒪**_*e*_ *to derive the partial order on* **𝒪**, *it is good to be aware that not all compositions from* **𝒪**_*e*_ *makes sense. Moreover most of the subsets of* **𝒪**_*e*_ *is not in* **𝒪**.

*E.g. if* **𝒪**_*e*_ *consists of cells, then a few cell from a liver and a few cell from a brain creating one gorganism does not seem to be worth investigating regarding life-level*.

#### Groups

The set **𝒢** contains all subsets of **𝒪** that are of interest regarding certain life-level investigations. It also contains the supporting-groups.

##### Example 3.1.4.

*The elements of a supporting-group may be put into one of the following categories, however the borders are usually not strict. Moreover one gorganism can belong to several such categories*.

- *Close relatives, e.g. parents, cousins, offspings, sibling, etc*.
- *Organisms from the same species*.
- *Organisms from the same genus/order/class/etc*.
- *Organisms functioning as* “*nutrient” for the organism*.
- *Organisms that* “*does” things which increase or decrease the life-level of the underlying organism either directly or indirectly (e.g. by one of its byproducts)*.

##### Example 3.1.5.

*A few examples for modeling situations via supporting-groups:*

1. *If we examine the life-level of single cells in a given animal, then can* **𝒢** *consist of all tissues / organs / organ systems of that animal*.
2. *A mitochondrion and the cell that it resides to can be mutually in the other’s supporting group*.
3. *If our targets are birds living in a given forest, then* **𝒢** *can consist of groups of nearby birds of the same species or some other species, groups of nearby trees, insects that the bird eats, etc*.
4. **𝒢** *can consists the groups of all members of all species living on a mountain i.e. one group consists of all members of a given species*.
5. *If we investigate the life-level of a man/woman then* **𝒢** *can consist of spouses, partners, families, groups of friends, people living in a street / town / county / country etc. Or all such*.
6. *A supporting-group of a plant (with some attracting flower) can consist of nearby living bees who pollinate it, a man who waters it regularly (in a dry area), a few chickens who eat its leaves, and of course some nearby plants of the same species*.
7. *The male of a European mantis can be in the supporting-group of the female due to many categories mentioned in 3.1.4, even as nutrient*.
8. *A cleaner wrasse and a shark whose teeth are cleaned by the small fish can be mutually in the other’s supporting group*.
9. *Consider an endoparasite living inside a host’s body. Then the host can be in its supporting-group and it can be essential from life-level perspective, e.g. when the host dies, the parasite dies as well*. *Of course, the parasite can be in the supporting-group of the host too. This is an example when a supporting-group supports negatively*.
10. *Consider all members of one given species of bacteria living in a given human’s gastrointestinal tract. In our model it can be a group-type gorganism. Its supporting-group can contain all other groups of species of bacteria living there, and also the human itself*.
11. *Let our group-type gorganism be a group of American bison. Its supporting-group can contain similar such groups. We remark that normally one would not add those other groups (or the members of them) to the supporting-group of a single bison because they would have negligible effect on the life-level of that single bison, however it may have important effect on the life-level of the group*.
12. *Depending on the model, an offspring can be in the supporting group of the parent, even if it lives far away from its parent and they never contact each other. For example if the model wants to take into account the survival time of some properties, then it could be the case (see subsection 4.4)*.
13. *A man alone on Mars without any other living being may be an example for a gorganism with an empty supporting-group (if we do not consider e.g. gut bacteria)*.

#### Sub-environments

##### Example 3.1.6.

*When an e-life-level-system is built, then the set of environment-particles* **𝒦**_*e*_ *contains the building blocks for the set of sub-environments* **𝒦**. *Certain subsets of* **𝒦**_*e*_ *are the elements of* **𝒦**. *A few examples:*

1. **𝒦**_*e*_ *can be all grains of sand in a desert*.
2. **𝒦**_*e*_ *can be all* 1*m*^2^ *area on the ground of a given forest*.
3. **𝒦**_*e*_ *can consist of all relatively small fields and hills on a specific area*.
4. **𝒦**_*e*_ *can consist all relatively small space parts in a given big lake i.e. basic space territories in the water that are somehow worth to be distinguished*.

##### Example 3.1.7.

*The set* **𝒦** *contains all important parts of the environment that can have effect to the life-level of gorganisms (subsets of* **𝒦**_*e*_ *when an e-life-level-system is built). A few examples:*

1. *A territory of a lion*.
2. *A suburban of a town where a man lives*.
3. *A breathing machine that keeps a person alive*.
4. *A high diopter glass of a man*.
5. *Clothes of a woman (in a cold area)*.
6. *A bone implant in a man*.
7. *The seminal fluid for a motile sperm cell*.

#### Time

##### Example 3.1.8.

*Examples on the set of time points* **𝒯** :

*1*. **𝒯** = ℝ.

*2*. **𝒯** = [0, +∞).

*3*. **𝒯** = [0, 1].

*4*. **𝒯** = ℕ *(we already mentioned that with explanation in 2.3.21 and 2.3.22)*.

*5*. **𝒯** *can be a single point only:* **𝒯** = {0}.

### 3.2 Various examples

Now we are going to present some more detailed examples with at least some thoughts on possible life-level values.

It is important to emphasize here, that in some of the examples we will mention species, not individual organisms and we will talk about their life-level. We will do that in the sense of section 9 and/or subsection 8.3. That is, we either think of some “mainstream” organisms of the species (see 9.0.6), or when we compare, then we think of the set of all life-level values of all organisms belonging to the species.

#### Example 3.2.1.

*It is widely accepted that a virus is not alive. Yes, it may be right on a digital life-scale, where 0 and 1 are the only possible values for life-level. However if we allow a much bigger scale, nothing will prevent us to assign a non-zero value to a virus. Or may be different but non-zero values to different viruses*.

*Just a few example and a possible order among them regarding life-level: virophages, satellite viruses*, “*regular” viruses, mimiviruses/mamaviruses (see also [15])*.

#### Example 3.2.2.

*Viroids also can be considered as alive in the new sense. However, clearly, one may want to assign a smaller life-level value to a viroid than a virus. Or more precisely a reasonable expectation can be that the maximum of life-levels of all viroids should not be larger than the minimum of life-levels assigned to all viruses*.

*The life-level of a prion is an interesting question. Whether it is worth choosing to be different than 0 or not*.

#### Example 3.2.3.

*A life-level of a single bacterium should be definitely greater than 0 and higher than the maximum of life-levels of all viruses. However different species of bacteria may have different life-levels*.

*Again: this comment is not about assigning values to species, it is more precisely said that the set of values assigned to all members of a species of bacteria is far from a similar set for other bacteria in some sense (see also subsection 8.3)*.

#### Example 3.2.4.

*One can compare life-levels of bacteria that lack a cell wall around their cell membranes (either naturally or artificially) to regular bacteria. E.g. consider protoplasts, spheroplast, L-form bacteria and bacteria from the genus Mycoplasma (see also [9])*. *Regarding life-level, a possible order among them can be: stable L-forms (that are unable to revert to the original bacteria), unstable L-forms (that can revert), Mycoplasma bacteria, regular bacteria. (Note that L-form bacteria can form in nature too with low probability, hence they can also be considered as natural phenomena.)*

#### Example 3.2.5.

*Suppose that a bacterium is infected by a virus, but it does not cause lysis, it just cause focal degeneration. Hence the bacterium does not die, but it cannot replicate itself. How much is it alive? At this stage, of course, its life-level must be definitely lower than a virus-less similar bacterium, but no doubt, this must be greater than 0*.

#### Example 3.2.6.

*We can end up with a similar result in ordering relation (not in values) if we examine the life-level of a bacterium whose nucleus was completely removed by some cell surgery, but the rest of the cell remained intact. See also 3.4.1*.

#### Example 3.2.7.

*One can also ask the life-level of a single cell in a muscle of an animal. We can compare its life-level to other cells in other organs or to some bacterium as well. One can expect that a bacterium has a higher life-level since it has capability to heredity while the muscle cell does not*.

#### Example 3.2.8.

*It may be an interesting question whether a stem cell possesses a higher life level than a somatic cell*.

#### Example 3.2.9.

*Is a nephron just a tiny unit in the kidney for a specific function that is a conglomerate of many cells and as such it just possesses the same life-level value as any of its cells? Or does it attain a higher life-level value in some sense?*

#### Example 3.2.10.

*We can assign a value to a given man’s liver, to the whole organ. To avoid confusion let us emphasize again we must not assign value to* “*liver” because no such thing as* “*liver"; there are individual livers existing at different time points*.

*To be more precise when we try to assign a value to a given man’s liver then somehow we may have to take into account the followings:*

- *The man’s health or the health of some related organs of the man (which belong to the supporting-group of the given liver). Or the health of livers of men living in the same household as the examined man, etc. Depends on our points of view or the scope of the research project*.
- *The environment where the man lives, smaller or larger*.
- *The time when we investigate (e.g. a year ago we might have ended up with a significantly different result)*.

#### Example 3.2.11.

*A single ant with or without its colony are two different bio-bits for investigation. Hence we may end up with different life-level results*.

*It is important to underline that examining the life level of the whole colony or the life-level of the single ant with its colony are two different things. In the first case the gorganism is the whole colony. In the second case the gorganism is the ant while its supporting-group is its colony*.

#### Example 3.2.12.

*Continuing the previous example 3.2.11, suppose that we separate the ant from its colony. If the colony exists and is prosperous in the distance then does it affect the life-level of our ant? And what if the colony deceases in the distance. How can we take that into account? We have several options of course; it is worth considering all of them and choose which one serves our purposes best*.

#### Example 3.2.13.

*Consider a male gorilla with his natal group, where his position is low in the dominant hierarchy, and he has no chance to upgrade. If he left the group and found a new one, he would become the dominant male there. This example demonstrates the significance of the supporting-group regarding life-level*.

#### Example 3.2.14.

*The life-level of an adult infertile mule should be less than its parents (parents’ life-level at the time of fertilization) however it should be still a very high value*.

#### Example 3.2.15.

*The life-levels of a man in a prosperous town or in a rain-forest alone or on Mars alone are three very different values showing that importance of the sub-environment*.

#### Example 3.2.16.

*A high fever may decrease the current life-level of an antelope compared to before and after. For applicable models see 4.2.6 and 4.4.34*.

#### Example 3.2.17.

*A man in coma must have a non-zero life-level even in the case when it is obvious that he will never awake. However he must have a less life-level than most human being not in coma. It can be an interesting question how its life-level may relate to the life-level of a 4-month old fetus*.

#### Example 3.2.18.

*The life-level of a man at the sate of clinical death can be definitely positive, even far from 0, but much lower than it was when the man was healthy and lower than any healthy man or woman. Its actual life-level value may depend on how much other data one can put into the model e.g. the current and/or previous health status of the man or his age, etc. E.g. if he is 90 years old then that simply may result significantly different values than if he is just 20. Also the sub-environment is crucial too, e.g. if he is in a hospital, that may increase a lot. For applicable models see 4.4.30*.

#### Example 3.2.19.

*Continuing 3.2.18, one can examine the life-level of a healthy muscle cell of a man who is currently at the sate of clinical death. Its value may depend on the chosen model. If one does not care about the* “*far” future, then its value can be as high as it was before. If one does want to take into account the possible future outcomes, then its value may significantly drop down*.

#### Example 3.2.20.

*The life-level of a completely disabled man without other people who supply him (without such supporting-group) is close to 0. With a suitable supporting-group it can be almost as high as for a completely healthy man*.

#### Example 3.2.21.

*We can consider the life-level of a human being at the age of 1-5-15-30-60-80 etc. At different ages one might want to assign different life-levels to the actual gorganism that is: a given human being at the age of x (with a certain sub-environment and supporting-group). Obviously it is not the age that only counts i.e. not the age itself, not the number itself that determines the assigned value. Of course it has significant effect on the life-level value. One might expect that the life-level curve is increasing till 30-40 and then decreasing. However it may depend on the actual human and his current sub-environment and supporting-group, and it can be different, depending on the individual case*.

#### Example 3.2.22.

*For investigating a couple of a man and a woman at the age of 20 we have (at least) two options:*

1. *They together can form a gorganism. Its life-level must be definitely higher than any of its components however it is not necessary that much higher. One of the reasons of being higher is that neither the man nor the woman can breed alone*.
2. *Another option for modeling this situation is examining e.g. the man alone and put the woman into his supporting-group*.

**Remark 3.2.23**. *A remark that is related to the previous example 3.2.22: In the commonly accepted sense, a virus is not considered alive, partly, because it cannot replicate alone, i.e. it has to use* “*outside” machinery for replication. However, on a general level, there are many species or even organisms that own that property*.

*E.g. a similar problem arises when one consider a self-incompatible plant that is pollinated by some specific insects, but that insect suddenly disappears in that region (e.g. Coelogyne fimbriata is pollinated by female Vespula wasps)*.

*One more example is a plant whose seeds are dispersed by some birds, and suppose that beneath the parent plant, the top layer of the ground cannot support the development of any seed (because it got poisoned), but a few hundred meters far, there would be suitable soil. Again, without the birds, the plant cannot replicate*.

*As another example, take a brood parasite who cannot replicate without some host species who raises its cub*.

*We can go on and observe that this only property is also valid for a man being alone*.

*Therefore adding another gorganism to the original gorganism that makes breeding/replication possible should increase life-level. Again, here we are not saying anything about a possible necessary alteration of the original life concept. We have just used the previous arguments for supporting our model*.

#### Example 3.2.24.

*The life-level of a small colony i.e. a few hundred men and women should be definitely higher than the life-level of a couple if all other parameters are more or less similar e.g. sub-environment*.

#### Example 3.2.25.

*Is the life-level of a man with an identical twin higher with his brother being alive and lower when not? It may depend on our points of view. See e.g. 4.4.16 and 7.2.12*.

#### Example 3.2.26.

*The life-level assigned to the whole biosphere of a big forest should be higher than the values of any of its components. In this case the gorganism is the whole biosphere itself. Its associated sub-environment can be the land where it is situated however we can add greater area or some ground below or some atmosphere above as well*.

#### Example 3.2.27.

*The whole biosphere of a small area is very similar to 3.2.26 in many senses however there may be differences. For example one may want to take into account the effects of the nearby ecosystems i.e. we can add the nearby similar biospheres to the supporting-group of the examined one*.

#### Example 3.2.28.

*The life-level of a 4-month old fetus should be definitely less than a new born baby and definitely greater than any cell or any organ of an adult*.

*A sperm or an egg cell must have non-zero life-level too however definitely lower than the zygote has after fertilization*.

*The life-level curve from a zygote to a new born baby has to be more or less increasing. Again individual cases may differ where* “*case” may mean not only the gorganism but also the associated sub-environment and/or the supporting-group*.

#### Example 3.2.29.

*When a fetus is inside its mother’s womb then a fetus and its mother can be considered as separate gorganisms or they can be considered as one single gorganism too depending on the point of view. Or we can investigate all three gorganisms at the same time as well*.

#### Example 3.2.30.

*When a frog has just swallowed a fly then we have a few options how to evaluate the life-level of the frog and the fly. We consider the moment when it has just happened thus the digestion has not started yet, the fly is undamaged at the moment. The options are:*

1. *One can examine the life-level of either the frog or the fly without taking into account what had happened. In this case the two life-levels can be the same than it was before*.
2. *One can put the fly into the frog’s supporting-group which can slightly increase the life-level of the frog comparing it to prior to this action*.
3. *One can put the frog into the fly’s supporting-group which can significantly decrease the life-level of the fly comparing it to prior to this action. It is important to emphasize that the life-level of the fly can be considered much lower despite of the fact that then the fly is still untouched by any dangerous effect. It is because that one might want to take into account the future effects of this action now*.
4. *One can examine the life-level of the set that contains both the frog and the fly as one single gorganism. In this case the life-level curve may decrease dramatically at the moment of the action. It can be an interesting question that when the fly deceases then how the life-level of this gorganism changes e.g. is it decreasing slightly or significantly? (see also 4.4.17)*

#### Example 3.2.31.

*Consider a lizard in its normal habitat (as sub-environment) together with the group of lizards living there (as supporting-group). However a bit further there is a just erupting volcano. If we change the sub-environment to the latter then we will end up dramatically lower life-level. This demonstrates that property (ss) is generally not true for* **𝒦** *(see 2.3.31)*.

#### Example 3.2.32.

*Consider a dominant male gorilla in his habitat (as sub-environment) together with the his family i.e. with a few females and a few cubs (as supporting-group). However another male in the bigger social group is just about to fight with our male for dominance. If we change the supporting-group to this bigger group then we may end up significantly lower life-level. This justifies that property (ss) is generally not true for* **𝒢** *(see 2.3.31)*.

#### Example 3.2.33.

*When an antelope migrates to a new field where the grass is higher, then it is an (partial) example for a development-path when the gorganism changes its sub-environment*.

#### Example 3.2.34.

*An antelope stays on a field for a longer time which dries out completely because of lack of rain. This is an (partial) example for a development-path when the sub-environment changes dramatically independently from the behavior of the gorganism*.

#### Example 3.2.35.

*There may not be questions regarding a self hibernated animal from life-level perspective, one may say that it is as much alive as when it is not hibernated. However one may have a model that is based on the activation time of certain properties, i.e. how long it takes to have the capability for some properties. In such a model, the life-level of a self hibernated animal must be a bit lower. See subsection 4.3*.

#### Example 3.2.36.

*One may get an analogous example as the previous one (3.2.35) when one examines the life-level of a pupa of an insect. It is analogous from the perspective that this type of animal lacks many properties (usual life characteristics) in this stage. Hence its life-level value should be (much) less than at the adult stage*.

#### Example 3.2.37.

*One can compare the life-level of an oocyte being dormant in the ovary to a fully developed and ovulated egg cell. Also, in that comparison one can take into account the low probability (approx*. 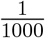 *in humans) of an oocyte becomes a fully developed and ovulated egg cell after it started to develop*.

*It is also interesting that it seems that one of the key factors of being dormant and remain viable for so long is that the oocyte has different mitochondrial metabolism than a fully developed egg cell has (see [21])*. *Hence its mitochondria also work differently. One can also examine how it may affect the life-level of mitochondria*.

#### Example 3.2.38.

*One can investigate the life stages of a butterfly, namely, the egg, larva, pupa and adult stages from life-level perspective. One may expect the following order in life-levels: egg, pupa, larva, adult. The low life-level of the pupa stage may decrease the life-level of the larva stage comparing to the similar developmental stage of other organisms*.

#### Example 3.2.39.

*An organism can be in the supporting-group of some other organism because of its byproducts, and not because of its direct actions. I.e. it modifies the sub-environment of the other organism only (either positively or negatively), but its actions does not touch the other organism directly*.

*E.g. the black rush (Juncus gerardii) alters soil conditions in ways that benefit other plant species in New England salt marshes. Juncus shades the soil surface, which reduces evaporation and thus lowers the concentration of salt in the soil. The presence of Juncus also increases soil oxygen levels. When Juncus was removed from areas of the marsh, those areas supported 50% fewer plant species. (See [2]54.)*

#### Example 3.2.40.

*The breeding cycle of the bird kakapo (Strigops habroptilus) is strictly linked to the fruiting cycle of the tree rimu (Dacrydium cupressinum). Breeding occurs only in years when rimu trees mast (i.e. fruit heavily), and rimu mast occurs only every three to five years. Hence rimu trees are significant elements in the supporting-group of kakapos. (See [6].)*

### 3.3 Some counter-intuitive examples

#### Example 3.3.1.

*On gorganism:*

1. *Consider a man with an organ transplant e.g. heart transplant. His body contains the heart of a different man with different DNA of course. Nevertheless the gorganism should be composed of not only all other parts of the body but also the transplanted heart*.
2. *One can examine the life-level of a cut arm of a starfish from the Linckia genus, which is capable for full regeneration (disk-independent bidirectional regeneration). Usually an arm is just a small part of the body with much less life-level value than the whole body, however in this case the difference should be much lower*.
3. *Conjoined twins have to be considered as one single gorganism, not two*.

#### Example 3.3.2.

*On the sub-environment:*

1. *A breathing machine that keeps a person alive is an essential part of his sub-environment*.
2. *An artificial cardiac pacemaker built in a man is part of the man’s environment and clearly significantly increases his life-level*.
3. *A man with a dental tooth filling. Then the filling material belongs to the environment*.
4. *A fish has swum into a plastic bag from which it cannot escape. In this case the plastic bag itself belongs to the sub-environment of the fish. Clearly without taking that into account we may end up with a false value for the life-level*.

#### Example 3.3.3.

*On the supporting-group:*

1. *Take an example for obligatory symbiosis e.g. siboglinid tube worms and its symbiotic bacteria. The supporting-group of each organism can be the other organism*.
2. *Take a brood parasite, e.g. a young cuckoo who is being raised by its host parents. Then one of its supporting-group can consists of its host parents. This group extremely increases the life-level of the cub, it is necessary for its survival*.
3. *Consider a kidney transplant i.e. a kidney before transplantation. Here the usual supporting-group (a man’s whole body) is completely missing. Therefore the life-level of that kidney is massively lower at this moment than it will be after the transplantation*.
4. *A lion lives on an area where a group of gazelles lives too and gazelle is the main food supply of the lion. In this case the group of gazelles can be the supporting-group of the lion (or at least part of)*.
5. *The life-level of a cat with no nails can be enormously different if it lives in the wild or with men*.

#### Example 3.3.4.

*On the importance of time and the life-level–time graph:*

1. *Something serious happens with a healthy young man who gets into the state of clinical death for a minute or two but soon afterwards he recovers completely. In this case his life-level decreased and then increased enormously in e.g. an hour. If we did not consider time then we would loose important information on life-level*. *We added more details to this example in 3.2.18*.
2. *See example 3.3.3 item 3 and compare life-level of the kidney before and after the transplantation*.

**Remark 3.3.5**. *Example 3.3.3 item 2 also provides an example for the case when O* ∉ *G, but the effect of G is essential. Moreover the elements of G are from different species*.

*We remark that one may also build his model in a way that he puts O into G however we think it is much more natural to keep them separately in this special example*.

#### Example 3.3.6.

*Take a given freeze tolerant insect, that can live a full life after being frozen and thawed. And freeze it totally. Then at this stage, its life-level may be set to almost as high as when it* “*lives", since only the* “*time is shifted". No doubt, its life-level can be definitely positive. For applicable models see subsection 4.4 and for explanation 4.4.18*.

#### Example 3.3.7.

*Consider a tissue culture that is kept alive and grown artificially. It is not a natural phenomenon again, however that is something to which a life-level value can be assigned and that can be useful. Of course, a tissue culture must possess a less life-level value than its parent organism. Certainly, here the sub-environment and the supporting-group have important role*.

*Similarly one can investigate a cultured organ. See also example 3.4.4*.

#### Example 3.3.8.

*Consider a plant that reproduces by vegetative propagation which has a fully developed daughter plant that is still connected to its parent and can get nutrients from the parent. When one separates the daughter from its parent, then the life-level of the daughter has to decrease slightly*.

#### Example 3.3.9.

*The body of a male animal produces seminal fluid for motile sperm cells that contains several components which promote the survival of the spermatozoa, and provide a medium through which they can move. Hence it can be considered as an essential sub-environment of the spermatozoon. In that sense* ***the body generates sub-environment*** *for some of its descendants, the sperms*.

#### Example 3.3.10.

*The few hundred species of bacteria living in the gastrointestinal tract are very important and valuable for many animals. Therefore it is essential to keep and maintain the sub-environment, mainly the inner part of the large intestine, to be a place where those bacteria can live and flourish (or at least that sub-environment should not be harmful to those bacteria). One example of that is when the antibody Immunoglobulin A (which can be found in the gut mucosa too) can help diversify the gut community (see [14])*. *Therefore, in that sense*, ***one gorganism maintains a sub-environment for some other gorganisms***, *partly according to the requirements of those other gorganisms*.

#### Example 3.3.11.

*A supporting-group of a cell of an animal body can be some other neighboring cells. This view can be important from several perspective e.g. the cells can communicate with each other and some of them affect the behavior of some others significantly, e.g. just think of apoptosis of the cell by the signal of some other neighboring cells*.

#### Example 3.3.12.

*Forsteropsalis pureora is a species of long-legged harvest spider where males may be one of three morphs that differ in chelicerae size, chelicerae shape, and body size. The alpha male has short yet strong chelicerae, the beta male has longer and thinner chelicerae, while the gamma male is seven times smaller than the other two. Obviously the life-level of males must also rely on their morphology hence can be remarkably different. Also it can depend on sub-environmental factors, and / or supporting-group factors too, e.g. proportions of the same and different type males in the close neighborhood. (see [20])*

*We remark that this is also an example where life-level cannot be simply assigned to a species*.

### 3.4 Examples for life-level through lifespan

Here we provide some examples that hopefully serve the understanding of the concept of development-path.

#### Example 3.4.1.

*During reproductive cloning, one examine the change of life-level of an egg cell whose nucleus has just been replaced by the nucleus of a somatic cell. It may massively increase the life-level of the cell. We note that the (half of the) DNA of the somatic cell may be slightly different from the DNA removed from the egg, because of possible mutations, therefore this operation (the result of the cell surgery) is not only an addition, it is partially an alteration as well*.

*The result can be even more dramatic, if we consider cross-species nucleus transfer, where the DNA of the mitochondria remains untouched. If the DNAs are strictly incompatible, then the cell may decease in a short time. In this case, we might have decreased the life-level of the original egg cell, instead of increasing it*.

*In these cases we always consider one development-path that starts with the original cell and goes on with the modified one after the surgery*.

#### Example 3.4.2.

*When a lizard loses its tail then one can maintain two development-paths: one continues on the* “*new” lizard (without tail), the other continues on the lost tail. Of course, normally the first one is investigated only, however the second one may have some sense too (e.g. one can ask how long the cells in the tail remain alive). We emphasize here that the lizard has changed a lot by that action, it is far more not the same object, but we need an identification through time how we proceed*.

*Taking the first (usual) development-path, one can investigate the life-level that drops down at the action, and gradually increases back to its previous level as the tail regrows*.

#### Example 3.4.3.

*Continuing example 3.2.29 one can have several options for development-paths to be examined*.

1. *If we started with the zygote then we may follow its life-level as a fetus and then a newborn baby (and so on)*.
2. *If we investigated the mother before fertilization then through pregnancy we can add the fetus to the supporting-group and we can also keep the newborn baby in that supporting-group after giving birth. In this case the mother always remain a separate gorganism from its embryo/fetus/newborn*.
3. *If we examined the fetus and the mother as one single gorganism then we can keep it that way as time goes by i.e. after birth we can still keep the mother and the newborn baby together and investigate the life-level of that gorganism*.
4. *In contrast to the previous point we may have a development-path that keeps them together for some time however it separates them at a later point of time e.g. at giving birth (but other options are possible too). Thus we have to make a choice if we follow the mother’s lifespan or the newborn’s lifespan. The two development-paths are the same till the separation but afterwards it diverts*. *It is important to mention that we can keep all two development-paths in our model or just one of them, as we prefer*. *Note that in these cases the continuity of the life-level function (in time) may not be hold*.

#### Example 3.4.4.

*Suppose that an organ is cultured from some stem cells in vitro. Then through development, the life-level of the different phases can be investigated, i.e. the change of life-level in time*.

#### Example 3.4.5.

*Consider a plant that reproduces by vegetative propagation which has already produced a root sprout which is not strong enough yet to be separated from its parent (it would not survive separation). At this stage one can consider both the root sprout and the parent as one gorganism or one can put the parent to the supporting-group of the root sprout that is currently necessary for its survival. In the first case there is one development-path which follows the changes of the parent and the sprout together, while in the second case there are separate development-paths of the sprout and the parent*.

*If from life-level perspective one models the phases of the plant developing from the root sprout, then one can end up with higher and higher life-level values. In the last stage, the new plant has full potential but is still connected with its parent*.

#### Example 3.4.6.

*How can one model if a man has a child and we would like to see it as a factor increasing his life-level? One option is to add the child to his supporting-group. The other option is that the child has its own development-path which has a common point in the past with the father’s development-path. More precisely there is a development-path that starts with the father, but at some point it switches to the sperm that will fertilize the egg cell from which the child evolved. On models providing such life-level calculations see subsection 7.4*.

#### Example 3.4.7.

*Before fertilization the sperm and the egg cell have separate development-paths, but at fertilization and after, the two paths become equal. They are still two development-paths, not one, but they are equal after that time point*.

#### Example 3.4.8.

*One more similar example: an apple has just fallen from an apple tree. A second ago we considered them as one gorganism. Now we want to see them separately. Regarding our development-path we have to make a decision if we follow the life line of the tree or the apple i.e. we may model this situation with two development-paths*.

#### Example 3.4.9.

*Continuing example 3.2.30 we have a few options for development-paths. It is not only the different ways we can choose, but also what events we include them*.

1. *We can consider the frog before and after the action and long after the action without taking into account what had happened. We will end up with the same life-level values independently of time*.
2. *Or one can put the action into the model which may result that at some time after the action, the life-level of the frog will be slightly higher*.
3. *The development-path of the fly must contain the action, and therefore as time goes by, the life-level will dramatically decrease and tends to 0 finally*.
4. *One can examine the life-level through a development-path of the gorganism that contains both the frog and the fly. It can be higher than the life-level of any animals before the action and at the moment of the action. But it may be decreasing as time goes by and the fly dies and vanishes. For applicable models see subsection 4.4 and see also 4.4.17*.

#### Example 3.4.10.

*One purpose of the notion of development-path is to express identity through (a long period of) time, when most (or all) components are interchanged, but we still consider the initial and the final gorganisms to be the same (not equal but the same)*.

*For example we can think of multi-cell organisms where some cells die, while new cells are created, but our perception gives that the gorganism is the same (that is of course not true if we examine just the matter that builds the organism)*.

*Or another example can be if we consider some social group of gorillas consisting of 5 members initially. In a longer period of time two things happen:*

- *All of those 5 members leave the group gradually, but at different point in time*.
- *In the mean time, new very similar members join the group (same age, same sex, etc.); when one leaves, a new member joins*.

*Therefore, finally, we end up with a group with totally different members, however our perception gives again that the group remained more or less intact (at least in its main functions)*.

## 4 Some possible ways to derive life-levels: life-level models

In this section we are going to provide some models that accomplish the requirements of the definition schemas. All of them are based on some general principles from which life-level values can be derived. All models provide a still very general view or better to say method on how one can derive values for one’s life-level function. They express possible ways that might be worth considering to use or combining with other principles.

We will present various groups of models. One of the key differences among them will be that we consider the present or some “close” time interval around the present or the “far” future of our bio-bit. Those models which consider the closer or further future actually express some future potential of the bio-bit as a life-level value. In other words measuring some biologically interesting future potential can serve as a life-level function (in the present). The simplest model is based on list of properties where property can be any attribute of generalized organisms. We assign weights to those properties, and summarize those weights that belong to properties that a given gorganism owns. We can get more sophisticated models if we take into account the activation frequency / time of the properties.

The next model is based on the survival time of properties. In the short run a property can either survive or can be lost and then perhaps reappear, all happening with the original generalized organism. In the long run a property may survive through inheritance however we will see that it is worth considering in the derivation that it is not the only option that the gorganism inherits it.

The most complex model is an enhancement of the previous one with adding and using probability distribution on the set of development-paths.

We will always try to establish the life-level function properties described in definition 2.1.10; we will refer to them in the same way as they are mentioned there. i.e (es), (do), (ss).

### 4.1 Properties

#### Definition 4.1.1.

*In the sequel* **𝒫** *will denote a countable set of properties. We also associate weights to the elements of* **𝒫** *i.e. there is a function w* : **𝒫** → (0, 1] *such that* 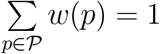.

*Such* ⟨ **𝒫**, *w*⟩ *pair is called a* ***property-system***, *while* **𝒫** *alone is called a* ***property set***. □

#### Definition 4.1.2.

*Let a life-level-subbase-system* Λ_*sb*_ = ⟨⟨ **𝒪** ⊂⟩,⟨ **𝒦** ⊂⟩,**𝒢**,, **𝒯**, ∈ Θ⟩ *and a property set be given. Let b* = (*O, K, G, t*) ∈ Θ *be an admissible bio-bit. If this bio-bit owns property p* ∈ **𝒫** *at time t, then we will denote this fact by*

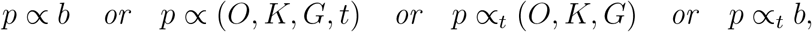

*and* ∝ *is called* ***property relation*** *associated to* Λ_*sb*_.

*If K and G are clear from the context then we can simply write p* ∝_*t*_ *O*.

*If a bio-bit b* ∈ Θ *does not own property p, then we use the notation* 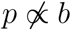.

*We also require that* 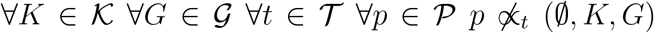 *holds*. □

#### Definition 4.1.3.

*If a life-level-subbase-system* Λ_*sb*_ *and a property set* **𝒫** *(or a property-system* ⟨**𝒫**, *w*⟩*) are given together with an associated property relation, then we will say that* ⟨Λ_*sb*_, **𝒫**, ∝⟩ *(or* ⟨Λ_*sb*_, **𝒫**, *w*, ∝⟩*) is a* ***propertized life-level-subbase-system*** *(similarly for life-level-base-system or life-level-system)*.

We do not give a definition what a property is, it is a primitive concept identified by the current research project. A property can be any attribute of generalized organisms or group of organisms that is worth investigating. E.g. a property can be a characteristic (or even a trait), but we deliberately use the word “property” here, because we want a more general concept than characteristic or trait.

Likewise, when a property is owned by a gorganism has to be determined by the application, i.e. this relation does not exist a priori.

#### Example 4.1.4.

*We present here some examples of properties without any explanation: (high) complexity, metabolism (the three types can be considered separately: creating energy, creating building blocks, eliminate waste), capability to reproduction, capability to growth and development, energy utilization, capability to respond to the environment, homeostasis, capability to evolutionary adaptation, capability to (exact) multiplication, capability to variation (in multiplication), capability to heredity*.

*Of course, generally, one may want to consider (much) simpler properties as well, e.g*. “*does fur cover the skin"*, “*it lays at least 3 eggs per nest", etc*.

**Remark 4.1.5**. *It is better to be careful about how we define our properties i.e. what our property exactly describes. This warning is about time. There are important biological properties which rarely hold always for an organism in a long time interval, but when they do, they are essential. E.g. consider property* “*responding to the environment". An organism does not do that all the time. It does sometimes and then it is important. Hence it is better to define our property as* “*capability to respond to the environment"*.

*A better example may be fertility. A young female fertile gorilla is not fertile at 100% of the time, where here fertility means strictly when she can get pregnant. Strictly speaking, it just takes a few days in a month, nevertheless we may consider her fertile all the time (up to our model)*.

*Let us investigate 4 examples from the respect of fertility in the strict sense. A totally frozen freeze tolerant insect is not fertile. A female cub gorilla is not fertile. An ovulating young gorilla is fertile. A young female gorilla outside of her ovulation period is not fertile. We emphasize that here we consider the very moment only*.

*We can refine our properties / property relations even more in a way that we require some time interval length during which that capability has to exist. Hence we do not require the capability to exist always, just exist regularly or at least once in a certain time interval. (cf. subsections 4.2.2 and 4.3)*

*Of course we can go even further on the way how detailed we define our properties. The more precise definitions we can provide, the better model we can have*.

*Nevertheless* ***the property relation is what really matters***, *therefore it is also a possibility that one keeps the name of its property as* “*responding to the environment", but one builds one’s property relation carefully according to the purposes of the current application*. □

**Remark 4.1.6**. *At a given point in time a bio-bit either owns a property or not. If a property is not applicable for a bio-bit, then we consider that the bio-bit does not own the property. E.g. the property* “*males fight for dominance” is not applicable for one single gorilla, hence such bio-bit does not own it. Or vice versa: The property of* “*fur covers its skin” is not applicable (directly) to a big group of gorillas, hence the group (as a gorganism) does not own it (cf. 4.1.13)*.

**Remark 4.1.7**. *At time t, generally a bio-bit* (*O, K, G*) *owns a property, not a gorganism O only. It is because either the sub-environment or the supporting-group or both can have influence if the given property is present or not, as e.g. the next example demonstrates it*.

#### Example 4.1.8.

*An animal may lose its fertility temporarily if it lives in an area with a lake containing some poisonous substance, however when it leaves that area, it gains back fertility almost immediately*.

*It is also an example to show that time is essential when we examine if a bio-bit owns a property or not. Because e.g. it can own a given property in several disjoint time intervals, and does not own it in between*.

**Remark 4.1.9**. *Weights are going to express the application’s preferences on the properties, which are more important, which are less. More precisely the function w is a property valuation. It will be essential in the later definitions and calculations*.

**Remark 4.1.10**. *If the property set* **𝒫** *is finite then it is possible to choose equal weights, however if it is infinite, then it is not possible*.

**Remark 4.1.11**. *Notice that in the notation p* ∝_*t*_ (*O, K, G*), *O is not necessarily a primitive gorganism, e.g. it can be a group-type gorganism as well*.

#### Definition 4.1.12.

*Let a propertized life-level-subbase-system* ⟨Λ_*sb*_, **𝒫**, ∝⟩ *be given. The relation* ∝ *is called* ***increasing*** *if O, O*′ ∈ **𝒪**, *O* ⊂ *O*^′^ *and p* ∝_*t*_ (*O, K, G*) *implies that p* ∝_*t*_ (*O*^′^, *K, G*).

If we have a property that is originally applicable for gorganisms with lower hierarchy-level then we can push the property upwards with a simple method.

#### Definition 4.1.13.

*Let a propertized life-level-subbase-system* ⟨Λ_*sb*_, **𝒫**, ∝⟩ *be given. Set*

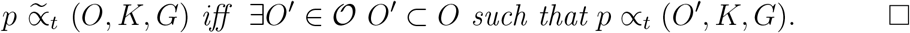

#### Proposition 4.1.14.

*Let a propertized life-level-subbase-system* ⟨Λ_*sb*_, **𝒫**, ∝⟩ *be given. Then the following statements hold*.

1. *If p* ∝_*t*_ (*O, K, G*) *then* 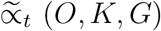,
2. *The operation* ^∼^ *is idempotent i.e*.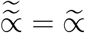,
3. 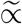 *is increasing*,
4. ∝ *is increasing iff* 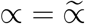.

*Proof*. (1) Obviously *O* ⊂ *O* gives the statement.

(2) By (1) we get that 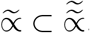 Thus we need the opposite. Let 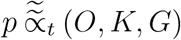. Then ∃*O*^*′*^ ∈ **𝒪** *O*^*′*^ ⊂ *O* such that 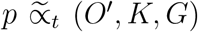. Which yields that ∃*O*^*″*^ ∈ **𝒪** *O*^*″*^ ⊂ *O*^*′*^ such that *p* ∝_*t*_ (*O*^*″*^, *K, G*). But then *O*^*″*^ ⊂ *O* gives that 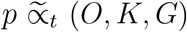.
(3) Let *O, O*^*′*^ ∈ **𝒪**, *O* ⊂ *O*^*′*^ and 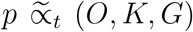. Then ∃*O*^*″*^ ∈ **𝒪** *O*^*″*^ ⊂ *O* such that *p* ∝_*t*_ (*O*^*″*^, *K, G*). But then *O*^*′*^ ⊂ *O*^*′*^ which implies that *p* ∝_*t*_ (*O*^*′*^, *K, G*).
(4) By (3) if 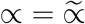 then ∝ is increasing.

To see the converse, observe first that by (1) 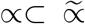 holds. In order to check the opposite, let 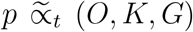. Then ∃*O* ∈ **𝒪** *O* ⊂ *O* such that *p* ∝_*t*_ (*O*^*′*^, *K, G*). By the increasing property we get that *p* ∝_*t*_ (*O, K, G*).

#### Example 4.1.15.

*In this sense we may say that a group of wolfs can hear ultrasound if any of the wolfs can hear ultrasound*.

If one has a continuous time model (**𝒯** ⊂ ℝ is an interval and *p* ∝_*t*_ *b* can be asked in any time point), then it can be a natural requirement that if for a bio-bit property *p* holds arbitrarily close to time point *t*, then it must hold in *t* too.

#### Definition 4.1.16.

*Let a propertized life-level-base-system* ⟨Λ_*b*_, **𝒫**, ∝⟩ *be given. We call* ∝ ***continuous*** *if*

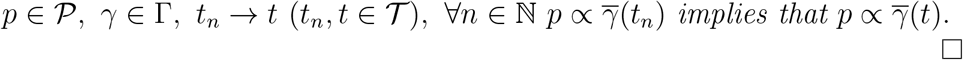

**Remark 4.1.17**. *Let us remark that a property set* **𝒫** *can have an only element i.e. we can define our model with an only property*.

**Remark 4.1.18**. *A very special but significant property: The property of the capability of inheriting other properties. This may be the most generic and most important among all properties. It is also clear that this property survives the longest (the precise definition of* “*property survival” comes in subsection 4.4)*.

*We think that a reasonable life-level function can be defined just based on that single property and some other additional assumptions (see also 4.4.19)*.

#### 4.1.1 Some generalization

One may want to have a more sophisticated system where a property is not only simply owned or not owned by a generalized organism (bio-bit), but also there is a factor which expresses that how much the organism (bio-bit) owns that property. We are going to add the basics of that here.

##### Definition 4.1.19.

*Let a life-level-subbase-system* Λ_*sb*_ = ⟨⟨**𝒪**, ⊂⟩, ⟨**𝒦**, ⊂⟩, **𝒢, 𝒯**, Θ⟩ *and a property set* **𝒫** *be given. Then* ∝ *is called a* ***property-owning-function*** *on* Λ_*sb*_ *if*

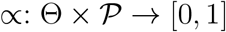

*such that* ∀*K* ∈ **𝒦** ∀*G* ∈ **𝒢** ∀*t* ∈ **𝒯** ∀*p* ∈ **𝒫** ∝ (∅, *K, G, t*), *p* = 0 *holds*. □

**Remark 4.1.20**. *It is a generalization of the previous notion (the simple property relation), since we may say that a bio-bit b owns property p, if* ∝(*b, p*) = 1, *and it does not own it, if* ∝(*b, p*) = 0.

**Remark 4.1.21**. *It is not necessary that the range of* ∝ *is* [0, 1]. *Someone can also implement it as* ∝: Θ × **𝒫** → ∞ [0, +), *however in the latter case there is no maximum value*.

**Remark 4.1.22**. *We use the same symbol* ∝ *for the simple relation and the function as well because from the context it has to be clear which one is applied*.

Now we generalize the increasing property for property-owning-functions.

##### Definition 4.1.23.

*Let a life-level-subbase-system* Λ_*sb*_ *and a property set* **𝒫** *be given with a property-owning-function* ∝. *Then* ∝ *is called* ***increasing*** *if p* ∈ **𝒫**, *b*_1_, *b*_2_ ∈ Θ, *b*_1_ ⊆_𝒪_ *b*_2_ *implies that* ∝(*b*_1_, *p*) ≤∝(*b*_2_, *p*). □

We can also have a kind of push up method similar to the one described in definition 4.1.13, however a much more complex one. For that, we also need a measure on **𝒪** (*t*) that can also be considered as if we had weights on the subsets of **𝒪** (*t*). We do the push up by recursion by the hierarchy-level of the gorganisms and technically we count the weighted average of the property-values on the co-atoms of the given group (those are the “highest” elements that “build” the group).

For recalling the notations *h*(*O*) and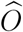, see 2.2.9 and 2.2.4 respectively. Now we define the co-atoms in 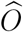 with respect to the partial order ⊂.

##### Definition 4.1.24.

*Let a life-level-subbase-system* Λ_*sb*_ *be given and let O* ∈ 𝒪. *Set*

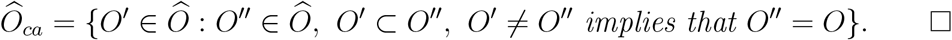

**Definition 4.1.25**. *Let* ⟨Λ_*sb*_, 𝒫, ∝⟩ *be given such that* Λ_*sb*_ *is a life-level-subbase-system*, 𝒫 *is a property set, an d* ∝ *is a property-owning-function that is defined on the set* {((*U, K, G, t*), *p* ∈ Θ(Λ_*sb*_) × 𝒫 : *h*(*O*) = 1}. *For every t* ∈ 𝒯 *let a measure μ*_*t*_ *be given on* 𝒪 (*t*) *such that μ*_*t*_ (𝒪 (*t*)) *<* ∞*and* 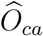 *is measurable when O* ∈ 𝒪 (*t*). *For* (*O, K, G, t*) ∈ Θ *and p* ∈ 𝒫, *we define* 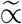 *recursively by hierarchy level h*(*O*).

*Set* 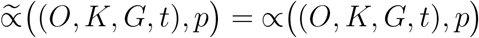 *when h*(*O*) = 1. *If* (*O, K, G, t*) ∈ Θ, *p* ∈ 𝒫 *and h*(*O*) = *n* + 1 *and* ∝ *is already defined for* ((*U,K,G,t*),*P*) *With h*(*U*) ≤ *n, then set*

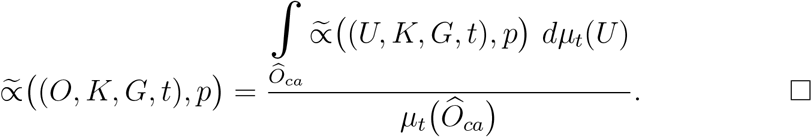

**Remark 4.1.26**. *If* 𝒪(*t*) *is finite in one’s model, then the previous formula reduces to*

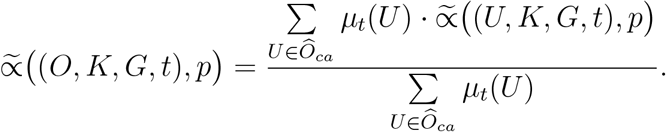

##### Example 4.1.27.

*Applying this method, the level of how much a group of wolfs can hear ultrasound is the weighted average of the level of that ability in the group. E.g. when creating a weight function μ on the elements of the group, one can give higher weights to young adult males than cubs. (cf. example 4.1.15)*

**Remark 4.1.28**. *Unfortunately* ∝ *is not necessarily increasing*.

### 4.2 Deriving life-level using owned properties

Suppose that we have gathered all properties into a set 𝒫 that we think that are essential for life. We also assign weights to our properties. In this case we can simply summarize those property weights that a given generalized organism (more precisely bio-bit) owns at a given point in time, and in this way, one can get a simple life-level function.

#### Definition 4.2.1.

*Let a propertized life-level-subbase-system* ⟨Λ_*sb*_, **𝒫**, *w*, ∝⟩*be given. Let t* ∈**𝒯** *and b* = (*O, K, G, t*) ∈ Θ(Λ_*sb*_) *be an admissible bio-bit. Set*

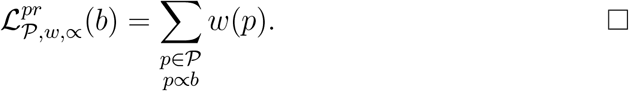

#### Proposition 4.2.2.

*Let* 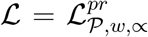 *be defined as in 4.2.1*. *Then the followings hold*.

1. Ran ℒ ⊂ [0, 1].
2. *If b* ∈ Θ, *then* ℒ(*b*) = 0 *iff b owns no property*.
3. *If b* ∈ Θ, *then* ℒ(*b*) = 1 *iff b owns all properties now (at π* _**𝒯**_ (*b*)).
4. ℒ *satisfies property (es)*.
5. *If* ∝ *is increasing, then* ℒ *has property (ss)*. □

**Remark 4.2.3**. *Evidently it may happen that* ℒ *does not satisfy property (do). If one wants that property, then one option is that one should add some very basic property to one’s property set, as e.g. the next example does*.

*Other option comes in subsection 6.3*.

#### Example 4.2.4.

*Take a freeze tolerant insect and freeze it totally. If we have e.g. high complexity among our properties, then in its current state, the life-level of our insect is not zero, it can be some small value according to this model. And after thawing, obviously it can gain back its previous life-level value*.

*We can end up with similar results for a dormant tree or seed by similar reasons. It is because we look at the present only. For more sophisticated models see subsection 4.4, for examples see 4.4.18, 4.4.32, 4.4.33*.

#### Example 4.2.5.

*A cub or a newborn is no doubt alive. However currently, in its current state, it lacks many important properties that are usually associated with life (e.g. heredity, high capability to respond to the environment, etc). If we apply this model to a cub, then it may provide life-level values according to our expectation that its life-level should be much higher than for a fetus, but much less than for a healthy adult*.

#### Example 4.2.6.

*If an antelope has high fever (causing by e.g. foot-and-mouth disease), then at the current moment the antelope will lack (at least partially) some basic usual life characteristics, e.g. capability to reproduction, high capability to respond to the environment, etc. If our model based on those properties, then it will assign lower life level to our antelope than when it was fully healthy. (cf. example 4.4.34)*

We can create a more sophisticated (generalized) version of definition 4.2.1 using property-owning-function instead.

#### Definition 4.2.7.

*Let a propertized life-level-subbase-system* ⟨Λ_*sb*_, **𝒫**, *w*, ∝⟩ *be given where* ∝ *is a property-owning-function. Let t* ∈ **𝒯** *and b* = (*O, K, G, t*) ∈ Θ(Λ_*sb*_) *be an admissible bio-bit. Set*

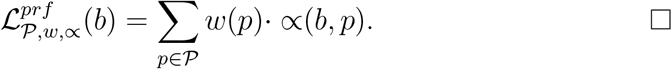

#### Proposition 4.2.8.

*Let* 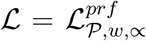 *be defined as in 4.2.7*. *Then the followings hold*.

1. Ran ℒ ⊂ [0, 1].
2. *If b* ∈ Θ, *then* ℒ(*b*) = 0 *iff b owns no property*.
3. *If b* ∈ Θ, *then* ℒ(*b*) = 1 *iff b (fully) owns all properties now (at π*_*𝒯*_ (*b*))
4. ℒ *satisfies property (es)*.
5. *If* ∝ *is increasing, then* ℒ *has property (ss). □*

#### Example 4.2.9.

*Let us suppose that one wants to take the value of fitness into account for one’s life-level-function. We have two comments for that*.

1. *If one wants* [0, 1] *to be the range of the property-owning-function, then e.g. one can apply the function* 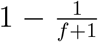 *where f is the usual fitness value*.
2. *If one considers fitness as a property of a population, not of an individual, then the property-owning-function (relation) can be implemented in a way that originally the function (relation) is defined for groups only. Then we inherit its value through the supporting-group, i.e*. ∝((*G, K*, ∅, *t*), *p*) *is defined first, and then we define* ∝((*O, K, G, t*), *p*) = ∝((*G, K*, ∅, *t*), *p*).

#### 4.2.1 A simple application of the property-owning model for some bio-bits

We are going to present a very simple application of this property-owning model for some simple bio-bits. More precisely we will examine some main-stream gorganisms only, we think of them being fully healthy, not damaged, not modified (see also 9.0.6). Moreover here we will take into account neither the effect of the sub-environment, nor the effect of the supporting-group (we will add a few remarks on that only).

First we build a simple property-system ⟨ **𝒫**, *w*⟩. Let **𝒫** = {*p*_*c*_, *p*_*m*_, *p*_*e*_, *p*_*hom*_, *p*_*gd*_, *p*_*res*_, *p*_*her*_, *p*_*evol*_}

where

- *p*_*c*_ = “high complexity”
- *p*_*m*_ = “metabolism”
- *p*_*e*_ = “energy utilization”
- *p*_*hom*_ = “homeostasis”
- *p*_*gd*_ = “capability to growth and development”
- *p*_*res*_ = “capability to respond to the environment”
- *p*_*her*_ = “capability to heredity”
- *p*_*evol*_ = “capability to evolutionary adaptation”.

We apply equal weights, i.e. 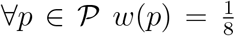. Now let us calculate the life-level of some gorganisms according to this model.

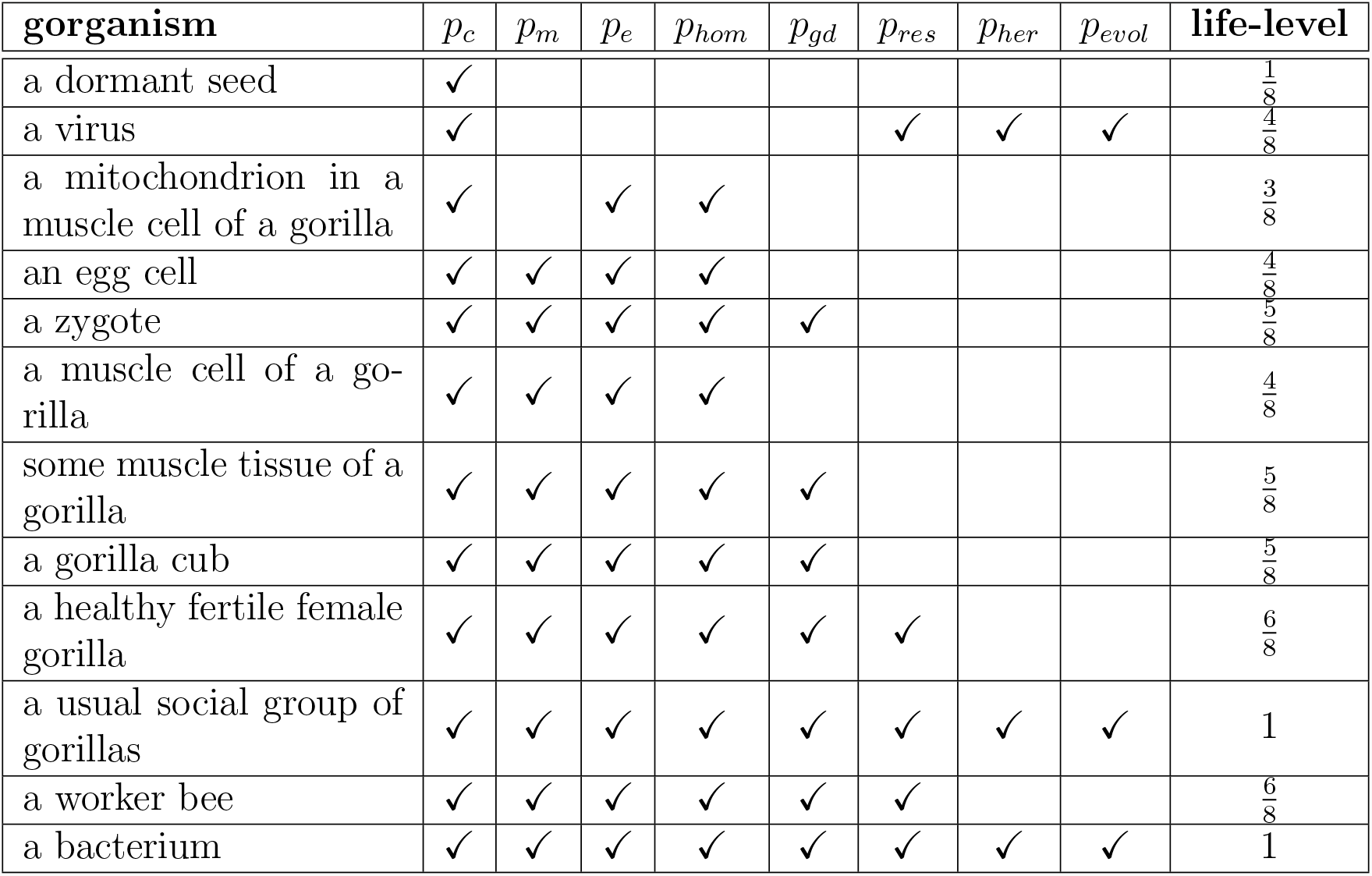

Some remarks:

- If *O* = “a mitochondrion in a muscle cell of a gorilla”, then 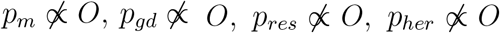 and 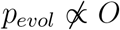, for the remaining *p* ∈ **𝒫** *p* ∝ *O* holds, hence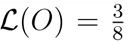. A mitochondrion cannot create all necessary building blocks for itself, that is why it does not own property *p*_*m*_. One can refine this model by separating metabolism into three sub-categories (creating energy, creating building blocks, eliminate waste) and then a mitochondrion could get a higher value, because it lacks only one from the three sub-properties of metabolism. Some of its genes stored in the nucleus of the cell, therefore it is not capable for heredity and evolutionary adaptation alone.
- If *O* = “a muscle cell of a gorilla”, then 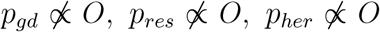 and 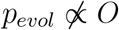, for all other *p* ∈ **𝒫** *p* ∝ *O* holds, hence 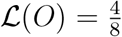. Here we think of a fully developed cell, that is why we set 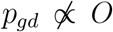. By “environment” we mean the outside environment of the animal, that is why 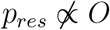.
- If *O* = “some muscle tissue of a gorilla”, then 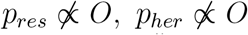 and 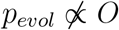, for all other *p* ∈ **𝒫** *p* ∝ *O* holds, hence 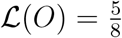. Again by “environment” we mean the outside environment of the animal, that is why 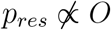
- If *O* = “a healthy fertile female gorilla”, then 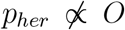 and 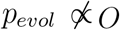 for all other *p* ∈ **𝒫** *p* ∝ *O* holds, hence 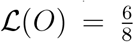 Without a supporting-group, a single animal cannot breed alone, hence 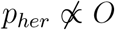 and 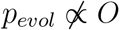.
- If *O* = “a healthy fertile female gorilla” and *G* denotes its social group, then *p*_*her*_ ∝ (*O, G*) and *p*_*evol*_ ∝ (*O, G*) and also all other properties are owned by (*O, G*), hence **ℒ** (*O, G*) = 1.
- The two properties *p*_*her*_ and *p*_*evol*_ always come together, i.e. both hold or none of them in a particular case. Hence one can consider uniting them into a single property.
- We applied equal weights here, but it is worth considering to distinguish properties more carefully.
- It can be seen that this very simple model does not provide all expected features that one would require. One way to achieve that is that to refine our properties and add more additional ones too. The other way is to find different principles as well (see subsequent subsections).

#### 4.2.2 Using activation frequency of properties

We mentioned in remark 4.1.5 that a gorganism rarely owns a property always, the property is switches on and off as time goes by. So definition 4.2.1 may be too strict. It is worth considering to weaken that in a way that we just require from a property to hold in a given time interval only. I.e. we should define a weaker, more permissive property owning relation as a first step.

For doing that, we have to specify the required minimum time interval length for each (property, bio-bit) pair, in which the property is required to be owned by the bio-bit, in order to be considered that the property is “really” owned. However we do not want that fine granularity since the interval length is usually species dependent only, not organism dependent. Hence we divide this assignment into two steps, for assigning a bio-bit to a specific group and then assign each group to a time interval length per property.

In that sense the title of the subsection may be misleading slightly because it is not a proper frequency that we want, since we just require that the property holds at least once in the associated (minimally expected) time interval.

##### Definition 4.2.10.

*Let a life-level-subbase-system* Λ_*sb*_ *is given. A* ***categorization*** *of the bio-bits is a function c* : Θ →*C where C is the set of categories*.

##### Definition 4.2.11.

*Let a propertized life-level-base-system* ⟨Λ_*b*_, **𝒫**, ∝⟩ *be given. Let c* : Θ → *C be a categorization of the bio-bits and let i* : **𝒫**×*C* → ℝ+ *be given. If p* ∈ **𝒫**, *b* ∈ Θ, *t* = *π*_***𝒯***_ (*b*), *γ* ∈ Γ_*b*_, *then set*

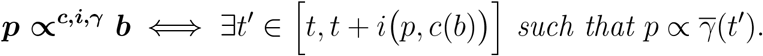

*If* Λ_*b*_ *is* Γ*-unique, then set* ***p ∝c***,***i b*** = *p* ∝^*c,i,γ*^ *b for the unique γ* ∈ Γ *such that* 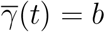

*We call [t, t*+*i (p, c*(*b*))] *the* ***required time frame*** *regarding p and b. □*

**Remark 4.2.12**. *The function c* : Θ→*C can be considered as categorizing the organisms into groups (e.g. species). While function i* : **𝒫** × *C* →ℝ+ *can be considered as the length of the expected minimum time intervals per (property, category) that is required for the property to be certainly owned by the bio-bit belonging to the category*.

##### Proposition 4.2.13.

*Let a propertized life-level-base-system* ⟨Λ_*b*_, **𝒫**, ∝⟩ *be given, where* Λ_*b*_ *is* Γ*-unique. Let c* : Θ → *C and i* : **𝒫** × *C* → ℝ+ *be given. Then* ∝⊂∝*c,i holds*.

##### Definition 4.2.14.

*Let a propertized life-level-base-system* ⟨Λ_*b*_, ***𝒫***, *w*, ∝⟩ *be given, where* Λ_*b*_ *is* Γ*-unique. Let c* : Θ → *C be a categorization of the bio-bits and let i* : **𝒫** × *C* → ℝ+ *be given. If b* ∈ Θ *then set*

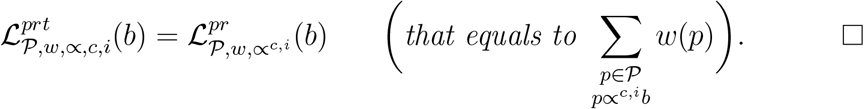

##### Proposition 4.2.15.

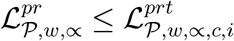

*Proof*. Obvious consequence of ∝⊂∝*c,i*. □

##### Example 4.2.16.

*For a dormant tree* 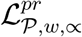 *may provide a value close to 0, or even 0, while* 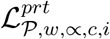 *may provide (almost) as high value as for a non-dormant one. For further possible refinements see subsection 4.3*.

##### Proposition 4.2.17.

*Let a propertized life-level-base-system* ⟨Λ_*b*_, **𝒫**, *w*, ∝⟩ *be given, where* Λ_*b*_ *is* Γ*-unique. Let c* : Θ → *C and i* : **𝒫** × *C* → ℝ+ *be given*. 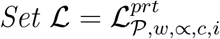 *Then the followings hold*.

1. Ran **ℒ** ⊂ [0, 1].
2. *If b* ∈ Θ, *then* **ℒ** (*b*) = 0 *iff* ∀*p* ∈ **𝒫** *b does not own property p within the time frame i (p, c*(*b*))
3. *If b* ∈ Θ, *then ℒ* (*b*) = 1 *iff b owns all properties within the required time frames*.
4. **ℒ** *satisfies property (es)*.
5. *If* Λ_*b*_ *is* Γ*-above-synchronized-forward, is* ∝ *increasing, then* **ℒ** *has property (ss)*.

**Remark 4.2.18**. *Evidently it is not necessary that ℒsatisfies property (do). However with some effort, one can choose one’s properties and the time frames per property (and species) such that it holds*.

Now we handle the case when Λ_*b*_ is not Γ-unique, first without probability.

##### Definition 4.2.19.

*Let a propertized life-level-base-system* ⟨Λ_*b*_, **𝒫**, *w*, ∝⟩ *be given. Let c* : Θ → *C and i* : **𝒫** × *C* → ℝ+ *be given. If b* ∈ Θ, *γ* ∈ Γ *then set*

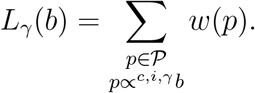

*If b* ∈ Θ *then set*

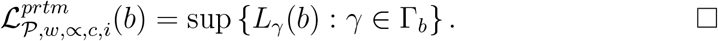

##### Proposition 4.2.20.

*Let a propertized life-level-base-system* ⟨Λ_*b*_, ***𝒫***, *w*, ∝⟩*be given. Let c* : Θ →*C and i* : **𝒫**×*C* → ℝ+ *be given. Let b*∈ Θ, *γ* ∈Γ. *Then the followings hold*.

1. *L*_*γ*_(*b*) = 0 *iff on development-path γ, b does not own any property within the required time frames*.
2. *L*_*γ*_(*b*) = 1 *iff on development-path γ, b owns all properties within the required time frames*.

##### Proposition 4.2.21.

*Let a propertized life-level-base-system* ⟨Λ_*b*_, **𝒫**, *w*, ∝⟩ *be given. Let c* : Θ →*C and i* : **𝒫**× → *C* ℝ^+^ *be given*. 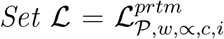

*Then the followings hold*.

1. Ran **ℒ** ⊂ [0, 1].
2. *If b* ∈ Θ, *then ℒ* (*b*) = 0 *iff for each development-paths b does not own any property within the required time frames*.
3. *If b*∈ Θ, Γ *is finite, then ℒ* (*b*) = 1 *iff there is a development-path such that b owns all properties within the required time frames*.
4. **ℒ** *satisfies property (es)*.
5. *If* Λ_*b*_ *is* Γ*-above-synchronized-forward*, ∝ *is increasing, then ℒ has property (ss)*.

Now we add probability as well.

##### Definition 4.2.22.

*Let a propertized life-level-base-system* ⟨Λ_*b*_, **𝒫**, *w*, ∝⟩ *be given, where* Λ_*b*_ *is countable and there is a probability function* P *for each* Γ_*b*_ (*b* ∈ Θ). *Let c* : Θ → *C and i* : **𝒫** × *C* → ℝ+ *be given. If b* ∈ Θ *then set*

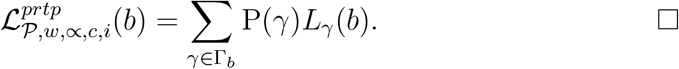

##### Proposition 4.2.23.

*Let a propertized life-level-base-system* ⟨Λ_*b*_, **𝒫**, *w*, ∝⟩ *be given, where* Λ_*b*_ *is countable and there is a probability function* P *for each* Γ_*b*_ (*b* ∈ Θ). *Let c* : Θ→ *C and i* ***𝒫***: ×*C*→ℝ ^+^ *be given*. 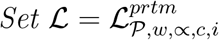.

*Then the followings hold*.

1. Ran **ℒ** ⊂ [0, 1].
2. *If b* ∈Θ, *then* ℝ (*b*) = 0 *iff for each development-paths b does not own any property within the required time frames*.
3. *If b* ∈ Θ, Γ *is finite, then* ℝ (*b*) = 1 *iff for all development-paths b owns all properties within the required time frames*.
4. **ℒ***satisfies property (es)*.

*Proof*.

1. Clearly ∀*b* ∈ Θ ∀*γ* ∈ Γ_*b*_ 0 ≤ *L*_*γ*_(*b*) ≤ 1 which yields the claim.
2. ℒ(*b*) = 0 iff ∀*γ* ∈ Γ_*b*_ *L*_*γ*_(*b*) = 0. Then apply 4.2.20.
3. ℒ(*b*) = 1 iff ∀*γ* ∈ Γ_*b*_ *L*_*γ*_(*b*) = 1. Then apply 4.2.20.

**Remark 4.2.24**. *One can make the models presented in this subsection simpler in a way that one chooses equal time frames per properties, species or both*. □

### 4.3 Deriving life-level using activation time of properties

We have already mentioned in the introduction and among the examples (or also in 4.1.5) that for a gorganism a property usually switches on and off as time goes by, hence it is not there always. Besides, usually it is not needed to be there always, it is enough if it switches on in the near future or it is there “frequently”. In the current model we are going to describe these phenomenons.

First we define the minimum time to be waited till the gorganism owns property *p*.

#### Definition 4.3.1.

*Let a propertized life-level-base-system* ⟨Λ_*b*_, **𝒫**, ∝⟩ *be given. Let b* = (*O, K, G, t*_0_) ∈ Θ, *p* ∈ **𝒫**. *Set*

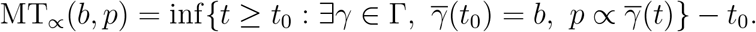

*Here we use the convention that* inf ∅= ∞. *We call* MT_*∝*_(*b, p*) *the* ***activation time*** *of property p regarding bio-bit b*.

#### Proposition 4.3.2.

*Let a propertized life-level-base-system* ⟨Λ_*b*_, **𝒫**, ∝⟩ *be given. Let b* ∈ Θ, *p* ∈ **𝒫**. *Then the followings hold*.

1. . 0 ≤ MT_*∝*_(*b, p*) ≤ ∞.
2. *p* ∝ *b implies that* MT_*∝*_(*b, p*) = 0.
3. MT_*∝*_(*b, p*) = ∞ *iff gorganism b will never own property p*.
4. *If* Λ_*b*_ *is finite*, ∝ *is continuous, then* MT_*∝*_(*b, p*) = 0 *implies that p* ∝ *b holds*.
5. MT_*∝*_((∅, *K, G, t*), *p*) = ∞.

*Proof*. We only show 4. First note that if Λ_*b*_ is finite, then so is Γ. If MT_*∝*_(*b, p*) = 0, then by finiteness of Γ, there is *γ* ∈ Γ and sequence (*t*_*n*_), such that 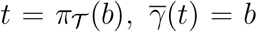 *t*_*n*_ → *t* and ∀*n* ∈ ℕ *p* ∝ *γ*(*t*_*n*_). By ∝ being continuous, we get that 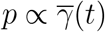.

#### Definition 4.3.3.

*Let a propertized life-level-base-system* ⟨Λ_*b*_, **𝒫**, *w*, ∝⟩ *be given. Let b* ∈ Θ. *Set*

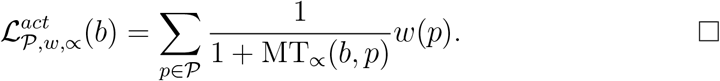

#### Proposition 4.3.4.

*Let a propertized life-level-base-system* ⟨Λ_*b*_, **𝒫**, *w*, ∝⟩ *be given. Let* 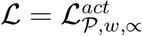 *be defined as in 4.3.3*. *Then the followings hold*.

1. Ran **ℒ**⊂ [0, 1].
2. *If* ∃*p* ∈ **𝒫** *such that p* ∝ *b, then* ℝ (*b*) *>* 0.
3. *If b* ∈ Θ, *then* ℝ (*b*) = 0 *iff b will never own any property*.
4. *If* Λ_*b*_ *is finite*, ∝ *is continuous and b* ∈ Θ, *then* **ℒ**(*b*) = 1 *iff b owns all properties now (at π* _***𝒯***_ (*b*)).
5. **ℒ***satisfies property (es)*.

*Proof*. Each statement is a consequence of the corresponding statement in proposition 4.3.2. □

#### Proposition 4.3.5.

*Let* 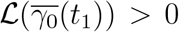 *be defined as in 4.3.3*. *If* Λ_*b*_ *is* Γ*-above-synchronized-forward and* ∝ *is increasing, then* **ℒ** *has property (ss)*.

*Proof*. Evidently if Λ_*b*_ is Γ-above-synchronized-forward, ∝ is increasing, *p* ∈ **𝒫**, *b*_1_, *b*_2_ ∈ Θ, *b*_1_ ⊆ _*𝒪*_ *b*_2_, then MT_*∝*_(*b*_2_, *p*) ≤ MT_*∝*_(*b*_1_, *p*) which implies that

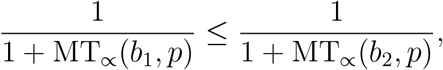

which yields that ℒ(*b*_1_) ≤ ℒ(*b*_2_).

#### Proposition 4.3.6.

*Let* 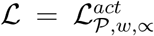 *be defined as in 4.3.3*. *If* Γ *is closed, then* **ℒ** *has property (do)*.

*Proof*. Let *γ*_0_ ∈ Γ, 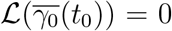 and suppose indirectly that ∃*t*_1_ *> t*_0_ such that 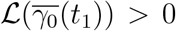. It means that ∃*p* ∈ **𝒫** such that MT 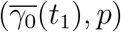, *p*) *<* ∞ which gives that ∃*γ*_1_ ∈ Γ such that 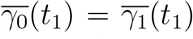 and ∃*t*_2_ ≥ *t*_1_ such that 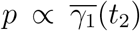. Now take 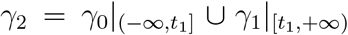 By Γ being closed we get that *γ*_2_ ∈ Γ. But then 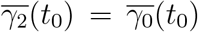 and 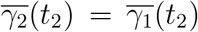 therefore 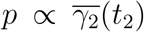 which yields that 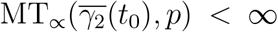 hence 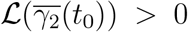 which is a contradiction. □

#### Example 4.3.7.

*One can apply this model to e.g. a dormant tree or a self hibernated or estivated animal. In each cases one may end up with a lower life-level comparing to the life-level of the “normal” state because of the longer activation times*.

*Of course the actual activation time does depend on where the organism is at its dormancy period. Furthermore if the organism just came out of dormancy, then at that very moment, one can get different activation time for certain properties for hibernation or estivation, because usually the recovery is much faster from estivation*.

*Let us calculate the difference between the life-level l*_*n*_ *of non-dormant state and the life-level l*_*d*_ *of dormant state. Suppose that in non-dormant state the activation time of property p is t*_*p*_, *and the length of dormancy is d. Then we get that*

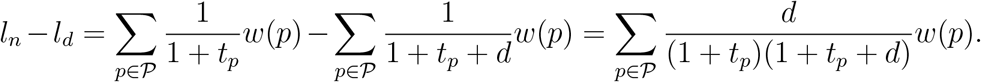

*If we simply replace* 1 + *t*_*p*_ + *d with* 1 + *d, then we get that*

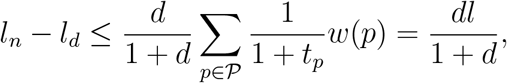

*which yields that*

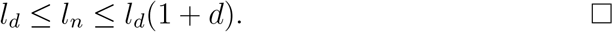

#### Example 4.3.8.

*According to this model one can investigate the life-level of a pupa of an insect as well. Unlike the previous example, here, the gorganism owns some properties (i.e. the activation time is 0), while there are many properties that need time to be activated. Or more precisely, there are properties with the same activation time as at adult state, and also there are others with additional time that is the length of the metamorphosis*.

#### 4.3.1 Using probability

Now we are going to give two more sophisticated versions of the previous notion 4.3.3 by using some probability.

In the sequel P(…) will denote the probability of an event.

First we define how long one has to wait till property *p* appears with probability 1.

##### Definition 4.3.9.

*Let a propertized life-level-base-system* ⟨Λ_*b*_, **𝒫**, ∝⟩ *be given. Let b* = (*O, K, G, t*_0_) ∈ Θ, *p* ∈ **𝒫**. *Set*

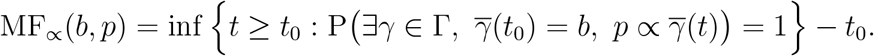

*Here we use the convention that* inf ∅ = ∞. □

##### Definition 4.3.10.

*Let a propertized life-level-base-system* ⟨Λ_*b*_, **𝒫**, *w*, ∝⟩

*be given. Let b* ∈ Θ. *Set*

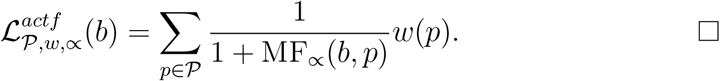

##### Proposition 4.3.11.

*Let* 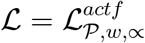 *be defined as in 4.3.10*. *Then the followings hold*.

1. Ran **ℒ** ⊂ [0, 1].
2. *If b* ∈ Θ, *then ℒ* (*b*) = 0 *iff b will never own any property with probability 1 on any of its development-paths*.
3. *If b* ∈ Θ, *then* **ℒ** (*b*) = 1 *iff for every small ε >* 0, *b owns all properties with probability 1 in the time interval* [*t, t* + *ε*] (*t* = *π*_***𝒯***_ (*b*)).
4. **ℒ** *satisfies property (es)*.

##### Definition 4.3.12.

*Let a propertized life-level-base-system* ⟨ Λ_*b*_, **𝒫** ∝⟩, *be given. Let* P *be a probability function that gives the probability that a property is owned by a bio-bit on some time interval. Then we call* ∝ **P*-increasing***

*if b* ⊆ _*𝒪*_ *b*^*′*^, *t*_0_ = *π* _***𝒯***_ (*b*) *implies that*

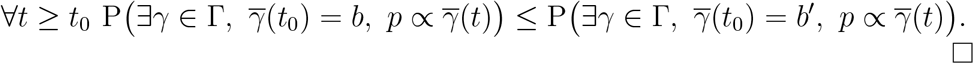

##### Proposition 4.3.13.

*Let* 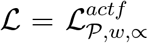 P*-increasing, then* **ℒ** *has property (ss). be defined as in 4.3.10*. *If* ∝ *is P-increasing, then ℒ has property(ss)*.

*Proof*. If *b* ⊆ _*𝒪*_ *b*^*′*^, *p* ∈ **𝒫**, then by ∝ being P-increasing we get that MF_*∝*_(*b*^*′*^, *p*) ≤ MF_*∝*_(*b, p*) which yields that ℒ(*b*) ≤ ℒ(*b*^*′*^).

**Remark 4.3.14**. *Of course, one can lighten definition 4.3.10 by modifying* MF_*∝*_ *(4.3.9) by adding an acceptable small threshold ε, that is instead of requiring probability 1, one would require that the probability is* ≥ 1 − *ε*.

The last version of such notions based on expected value of waiting time. First we define the expected value of time when property *p* is owned by a gorganism.

##### Definition 4.3.15.

*Let a propertized life-level-base-system* ⟨Λ_*b*_, **𝒫**, ∝⟩ *be given such that* **𝒯** = ℝ. *Let b* = (*O, K, G, t*_0_) ∈ Θ, *p* ∈ **𝒫**. *Set*

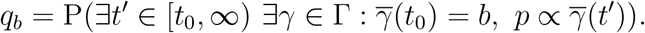

*If q*_*b*_ = 0, *then set* E_*∝*_(*b, p*) = ∞, *otherwise set*

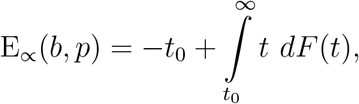

*where*

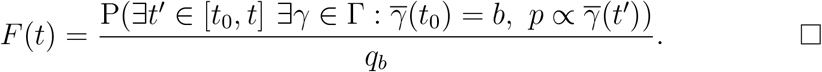

**Remark 4.3.16**. *F* (*t*) *is the conditional probability that property p appears on any development-path in the time interval* [*t*_0_, *t*]. *(Conditional since “we are in the event” that property p appears in* [*t*_0_, ∞) *with positive probability.) Hence F is the conditional cumulative distribution function of the random variable that gives the first time occurrence of property p in* [*t*_0_, ∞). *Furthermore* E_*∝*_(*b, p*) *is the conditional expected value of that random variable minus t*_0_.

##### Proposition 4.3.17.

*Let a propertized life-level-base-system* ⟨Λ_*b*_, **𝒫**, ∝⟩ *be given such that* **𝒯** = R. *Let b* ∈ Θ, *p* ∈ **𝒫**. *Then the followings hold*.

1. 0 ≤ E_*∝*_(*b, p*) ≤ ∞.
2. *If b* ∈ Θ *and it has 0 probability that b will ever own property p on any of its development-paths, then* E_*∝*_(*b, p*) = ∞.
3. *If b* ∈ Θ, *then* E_*∝*_(*b, p*) = 0 *iff* P(*p* ∝ *b*) = *q*_*b*_.
4. E_*∝*_((∅, *K, G, t*), *p*) = ∞.

*Proof*. It is clear that E_*∝*_(*b, p*) = 0 holds if and only if for every small *ε >* 0, *b* owns property *p* with probability *q*_*b*_ on a development-path in the time interval [*t, t* + *ε*] where *t* = *π*_***𝒯***_ (*b*). But P is a measure, therefore if ∀*n* ∈ ℕ 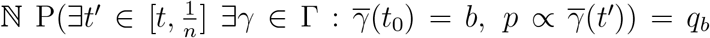 then it implies that P(*p* ∝ *b*) = *q*_*b*_. And trivially the converse is true. □

##### Definition 4.3.18.

*Let a propertized life-level-base-system* ⟨Λ_*b*_, **𝒫**, *w*, ∝⟩ *be given. Let b* ∈ Θ. *Set*

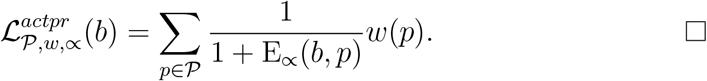

##### Proposition 4.3.19.

*Let* 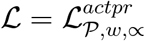 *be defined as in 4.3.18*. *Then the followings hold*.

*1*. Ran **ℒ** ⊂ [0, 1].

1. *If b* ∈ Θ *and it has 0 probability that b will ever own any property on any of its development-paths, then* **ℒ** (*b*) = 0.
2. *If b* ∈ Θ, *then* **ℒ** (*b*) = 1 *iff* P(*p* ∝ *b*) = *q*_*b*_.
3. **ℒ** *satisfies property (es)*.

*Proof*. Each statement is a consequence of the corresponding statement in proposition 4.3.17. □

**Remark 4.3.20**. *It is easy to construct an example that shows that the converse of proposition 4.3.19 2 (and 4.3.17 2) is not necessarily true*.

We validate property (do) in a special case only.

##### Proposition 4.3.21.

*Let a propertized life-level-base-system* ⟨Λ_*b*_, **𝒫**, *w*, ∝⟩ *be given. Set* 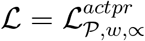. *If* Λ_*b*_ *is* Γ*-unique, then* **ℒ** *has property (do)*. □

*Proof*. Let *γ* ∈ Γ, *t*_0_ ∈ **𝒯**, *γ* _***𝒪***_ (*t*_0_) ?= ∅, 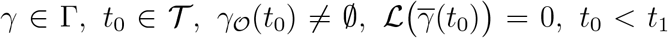 *t*_0_ *< t*_1_. We have to show that 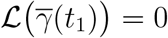

Let 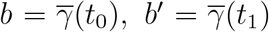 Obviously ∀*p*∈**𝒫** E_*∝*_(*b, p*) = ∞ holds. Then there are three cases.

1. *q*_*b*_ = 0. Clearly we get that *q*_*b*_*′* ≤ *q*_*b*_, hence *q*_*b*_*′* = 0, which yields that E_*∝*_(*b*^*′*^, *p*) = ∞ too, and then **ℒ** (*b*^*′*^) = 0.
2. *q*_*b*_ *>* 0, *q*_*b*_*′* = 0. Evidently we are done.
3. *q*_*b*_ *>* 0, *q*_*b*_*′ >* 0. We show this case when *F* is differentiable. Clearly

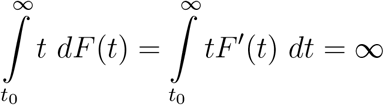

which gives that 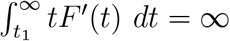 holds too. Let

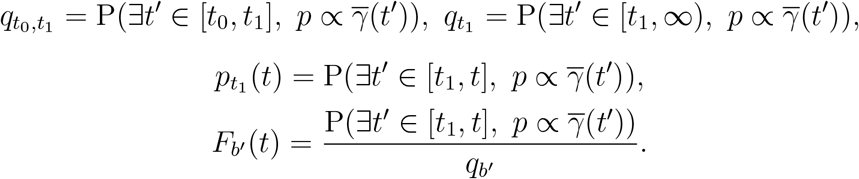

Note that 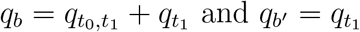 and 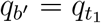. But if *t* ≥ *t*_1_, then

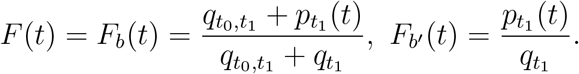

Then

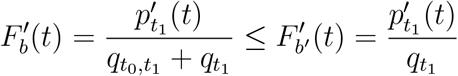

which gives that

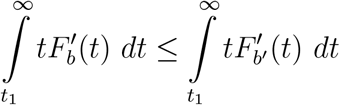

hence 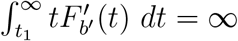 holds too.

### 4.4 Deriving life-level using survival time of properties

First we now define how long after time point *t*_0_ property *p* survives on the given development-path *γ*. Recall the notation that 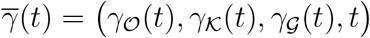 With this notation, 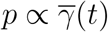 is equivalent with *p* ∝_*t*_ *γ*(*t*), and this expresses if the bio-bit *γ*(*t*) owns property *p* at time *t*.

#### Definition 4.4.1.

*Let a propertized life-level-base-system* ⟨Λ_*b*_, **𝒫**, ∝⟩ *be given. Let t*_0_ ∈ **𝒯**, *p* ∈ **𝒫**, *γ* ∈ Γ. *Set*

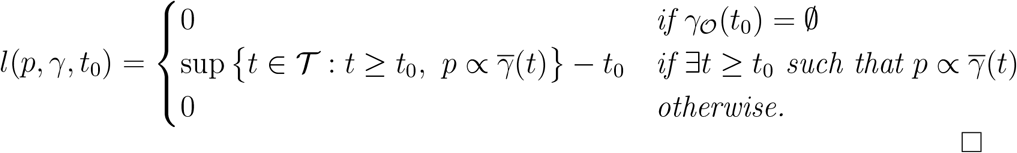

**Remark 4.4.2**. *Of course, by the previous definition it can be possible that l*(*p, γ, t*_0_) = +∞, *however we will always assume that l*(*p, γ, t*_0_) *is finite. Allowing infinity would cause behavior that we want to avoid*.

#### Example 4.4.3.

*It can happen that* 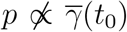 *however* ∃*t > t*_0_, 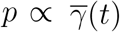 *A simple example is when a cub lacks a property, when it grows up, it owns it*.

*A more interesting example regards for the corresponding phenotype of a recessive allele. In this case the organism itself does not own that property, but an offspring can own it, and if that offspring is in γ, then* 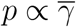 *t*) *holds for a t* ∈ **𝒯** *big enough*.

#### Proposition 4.4.4.

*Let a propertized life-level-base-system* ⟨Λ_*b*_, **𝒫**, ∝⟩ *be given. Let t*_0_, *t*_1_ ∈ **𝒯**, *t*_0_ *< t*_1_, *p* ∈ **𝒫**, *γ* ∈ Γ. *If γ*_***𝒪***_ (*t*_0_) ?= ∅, *then l*(*p, γ, t*_0_) ≥ *l*(*p, γ, t*_1_) *i.e. the function l is decreasing (as a function of t)*.

*Proof. l*(*p, γ, t*_1_) = 0, then nothing to prove. Suppose that ∃*t*_2_ ∈ **𝒯**, *t*_2_ ≥ *t*_1_ such that 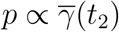 Then *l*(*p, γ, t*_0_) ≥ *t*_2_ − *t*_0_, while *l*(*p, γ, t*_1_) ≥ *t*_2_ − *t*_1_.

#### Corollary 4.4.5.

*Let a propertized life-level-base-system* ⟨Λ_*b*_, **𝒫**, ∝⟩ *be given. Let t*_0_, *t*_1_ ∈ **𝒯**, *t*_0_ *< t*_1_, *p* ∈ **𝒫**, *γ* ∈ Γ. *If γ*_***𝒪***_ (*t*_0_)≠ ∅ *and l*(*p, γ, t*_1_) *>* 0 *then l*(*p, γ, t*_0_) *>* 0.

We now define how long property *p* survives in the future bio-bits of (*O, K, G, t*) by Γ. □

#### Definition 4.4.6.

*Let a propertized life-level-base-system* ⟨Λ_*b*_, **𝒫**, ∝⟩ *be given. Let p* ∈ **𝒫** *and t* ∈ **𝒯**. *Let b* = (*O, K, G, t*) *be an admissible bio-bit*.

*Set*

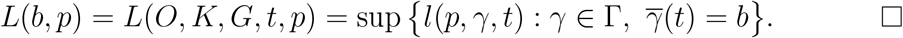

#### Proposition 4.4.7.

*Let a propertized life-level-base-system*, ⟨ Λ_*b*_, **𝒫**, ∝⟩ *be given. Let p* ∈**𝒫** *and t* ∈ **𝒯**. *If γ* ∈ Γ, *then* 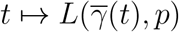 *is a decreasing function of t*.

*Proof*. Consequence of 4.4.4.

Now we are at the position to derive a life-level function based on the survival time of properties. It is the weighted average of the survival times of properties from **𝒫** using the associated weight function *w*.

#### Definition 4.4.8.

*Let a propertized life-level-base-system* ⟨Λ_*b*_, **𝒫**, *w*, ∝⟩ *be given. Let t* ∈ **𝒯** *and b* = (*O, K, G, t*) *be an admissible bio-bit. Set*

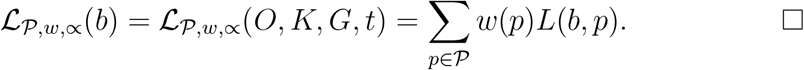

#### Proposition 4.4.9.

*Let* ℒ = ℒ _***𝒫***,*w,∝*_ *be defined as in 4.4.8*. *Then*

1. Ran 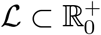
2. **ℒ** (*b*) = 0 ⇔⇒ ∀*t > t*_0_, ∀*γ* ∈ Γ 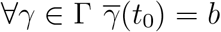 *implies that* 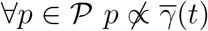 *(π*_***𝒯***_ (*b*) = *t*_0_)
3. **ℒ** *has property (es)*.

*Proof*.

1. Obviously *l*(*p, γ, t*_0_) ≥ 0 which implies that *L*(*O, K, G, t*_0_, *p*) ≥ 0 which implies that **ℒ** (*O, K, G, t*_0_) ≥ 0.
2. 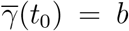 implies that 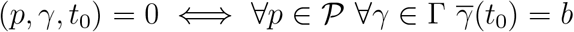 implies that 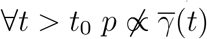
3. Property (es) follows from the fact the when *γ*_***𝒪***_ (*t*_0_) = ∅then *l*(*p, γ, t*_0_) is defined to be 0.

#### Definition 4.4.10.

*Let a life-level-system* Λ *and a property set* **𝒫** *be given. Then* **ℒ** *is called* ***normal regarding 𝒫*** *if b* ∈ Dom ℒ, ℒ(*b*) *>* 0 *implies that* ∃*p* ∈ **𝒫** *such that p* ∝ *b*.

**Remark 4.4.11**. *This property (normality) not necessarily holds for a life-level function ℒ* _***𝒫***,*w,∝*_ *defined as in 4.4.8*. *It may happen that at time t*_0_ (*O, K, G*) *does not own any property, however at a later time it owns some*.

#### Theorem 4.4.12.

*Let* ℒ = ℒ_***𝒫***,*w,∝*_ *be defined as in 4.4.8*. *If* Γ *is closed, then* **ℒ** *has property (do)*.

*Proof*. Let *γ*_0_ ∈ Γ, *γ*_0_(*t*_0_) = (*O, K, G*). Let 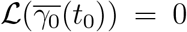 *t*_0_ *< t*_1_ and suppose indirectly that 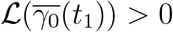 Then ∃*p* ∈ **𝒫** such that 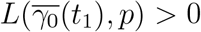. Which implies that ∃*γ*_1_ ∈ Γ, *γ*_1_(*t*_1_) = *γ*_0_(*t*_1_) such that *l*(*p, γ*_1_, *t*_1_) *>* 0. This yields that ∃*t*_2_ ∈ **𝒯** such that 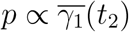.

Let 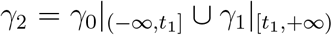 Closeness of Γ gives that *γ*_2_ ∈ Γ. Clearly *p* ∝ 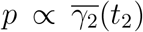 holds. It gives immediately that *l*(*p, γ*_2_, *t*_1_) *>* 0. By *t*_0_ *< t*_1_ we get that *l*(*p, γ*_2_, *t*_0_) *>* 0. Which implies that 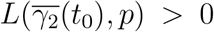 Using *γ*_2_(*t*_0_) = *γ*_0_(*t*_0_), it is equivalent with 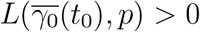. Then obviously we get that 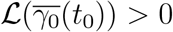 which is a contradiction. □

#### Theorem 4.4.13.

*Let a propertized life-level-base-system* ⟨Λ_*b*_, **𝒫**, *w*, ∝⟩ *be given. Let* ℒ = ℒ _***𝒫***,*w,∝*_ *be defined as in 4.4.8*. *If* Λ_*b*_ *is* Γ*-above-synchronized-forward and* ∝ *is increasing then* **ℒ** *has property (ss)*.

*Proof*. If *b, b*^*′*^ ∈ Θ(Λ_*b*_), *b* ⊆_*𝒪*_ *b*^*′*^, *t*_0_ = *π*_***𝒯***_ (*b*) then it is enough to show that ∀*p* ∈ **𝒫** *L*(*b, p*) ≤ *L*(*b*^*′*^, *p*). For that, it is enough to prove that ∀*p* ∈ **𝒫** ∀*γ* ∈ Γ 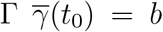 implies that ∃*γ*^*′*^ ∈ Γ 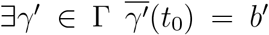, *l*(*p, γ, t*_0_) ≤ *l*(*p, γ*^*′*^, *t*_0_). If *l*(*p, γ, t*_0_) = 0 then it is obvious. Otherwise let *ε >* 0 and choose *t*_1_ ∈ **𝒯** such that *t*_0_ + *l*(*p, γ, t*_0_) − *ε < t*_1_ and *p* ∝ 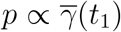 Choose *γ*^*′*^ according to the Γ-above-synchronized-forward property. By ∝ being increasing we get that *p* ∝ 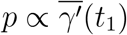 holds which gives that *l*(*p, γ, t*_0_) − *ε < t*_1_ − *t*_0_ ≤ *l*(*p, γ*^*′*^, *t*_0_). While *ε* was arbitrary, *l*(*p, γ, t*_0_) ≤ *l*(*p, γ*^*′*^, *t*_0_) holds. □

#### Proposition 4.4.14.

*Let a propertized life-level-base-system* ⟨Λ_*b*_, **𝒫**, ∝⟩ *be given. If γ* ∈ Γ, *then* 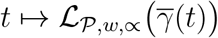 *is a decreasing function of t*.

*Proof*. Consequence of 4.4.7.

#### Proposition 4.4.15.

*Let a life-level-base-system* Λ_*b*_ *and a property-system* ⟨ **𝒫**, *w*⟩ *be given together with two associated relations* ∝_1_, ∝_2_ *such that* ∝_1_⊂∝_2_. *Then* 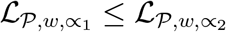

*Proof*. For the associated functions *l*_1_, *l*_2_, *L*_1_, *L*_2_ obviously hold that *l*_1_ ≤ *l*_2_ and *L*_1_ ≤ *L*_2_.

We present some examples where we demonstrate that this method provides results that are in accordance with our expectations.

#### Example 4.4.16.

*Let us apply this method for two gorganisms, two identical twins and for just one of the twins, and compare the life-level of those. We clearly get that the life-level of the latter is smaller than life-level of the first one. For another model, see 7.2.12*.

#### Example 4.4.17.

*Let us recall example 3.2.30 option 4*. *When the frog and the fly exist separately, then both have the possibility to “inherit” their properties or at least keep most of their properties for a longer time. However when the frog swallows the fly, the fly looses this kind of ability hence the life-level of the gorganism that contains both the frog and the fly decreases dramatically at that moment*.

#### Example 4.4.18.

*Let us apply this method for example 3.3.6*. *First we remark that in the freezing state, more precisely at a time point when it is frozen, our insect has no properties at all, or almost none. Nevertheless it is clear that freezing actually causes a small risk for such insects hence the survival time of properties may decrease a little only. Or they can also increase by the length of the freezing period (if it is a longer time period), hence in this case, freezing can increase life-level according to the current model. It is understandable on the stage of this model, since if we compare its life-level to a similar but not frozen insect, then the frozen one may “live” longer in time, i.e. it will “vanish” later in time*.

*We can end up with similar results for a dormant tree or seed by similar reasons. Actually from this perspective, there is small difference between a frozen freeze tolerant insect and a dormant tree*.

*Comparing a dormant tree and a dormant seed, clearly the tree must have a much “richer” development-path structure, richer in the sense that it may provide longer survival times of its properties. Hence most probably we get higher values for the tree than the seed according to this model. See also 4.4.32 and 4.4.33*.

**Remark 4.4.19**. *Continuing remark 4.1.18, let* ***𝒫*** *consist of a single property: the property of the capability of inheriting other properties. Without any additional assumption, we end up with a life-level function ℒ*_***𝒫***,*w,∝*_ *that assigns 0 to organisms that lost their fertility and there is no chance that they can gain it back. In that sense, this model seems to be very strict. For possible amendments see subsection 7.2*.

We are going to provide a kind of enrichment of this model in subsection 7.2.2.

#### 4.4.1 Some generalization

We can create a more sophisticated (generalized) version of definition 4.4.1 using property-owning-function instead.

##### Definition 4.4.20.

*Let a propertized life-level-base-system* ⟨Λ_*b*_, **𝒫**, ∝⟩ *be given. Let t*_0_ ∈ **𝒯**, *p* ∈ **𝒫**, *γ* ∈ Γ. *Set* 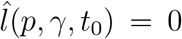 *if γ*_***𝒪***_ (*t*_0_) = ∅, *otherwise set*

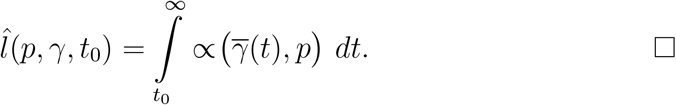

**Remark 4.4.21**. *Again, we want to avoid that* 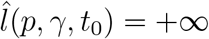 *hence we assume that* 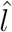 *is finite (see remark 4.4.2)*.

Using function 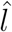, one can go on and define the same more complex functions based on 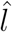 We are not going to fulfill that now.

#### 4.2.2 Using probability distribution on Γ

Now we are going to enrich the previous model with adding a probability distribution on special subsets of Γ.

First let us explain it roughly how we apply probability theory here. The probability is not necessary given on elements of Γ, it is because some development-paths belong together, expressing one possible way of development of the original bio-bit, i.e. they all may happen in the future, or none of them. Hence such set has a probability, not the single development-paths that constitute it. Of course if the set has an only element, then we may consider it as if the element had a probability.

If a bio-bit is given at the present, then we can split all of its possible future development-paths into separate subsets, according to that a subset represents one possible outcome, i.e. one and only one such subset of development-paths will happen in the future. We can assign a probability to each subset and use it to create a more sophisticated model.

We now define a specialized version of *L*(*b, p*).

##### Definition 4.4.22.

*Let a propertized life-level-base-system* ⟨Λ_*b*_, **𝒫**, ∝⟩ *be given. Let p* ∈ **𝒫**. *Let b* = (*O, K, G, t*) *be an admissible bio-bit and* Γ^*′*^ ⊂ Γ. *Set*

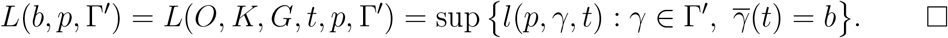

**Remark 4.4.23**. *With this new notation, clearly L*(*b, p*) = *L*(*b, p*, Γ).

We now define the expected value of how long property *p* survives on the development-paths of the bio-bit (*O, K, G*).

##### Definition 4.4.24.

*Let a propertized life-level-base-system* ⟨Λ_*b*_, **𝒫**, ∝⟩ *be given. Let p* ∈ **𝒫**. *Let b* = (*O, K, G, t*) *be an admissible bio-bit. Let a* 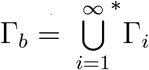 *be a countable disjoint partitioning of* Γ_*b*_ *and let a probability function* P *be given on this partitioning, i.e. it assigns value to each* Γ_*i*_ (*i* ∈ ℕ). *Set*

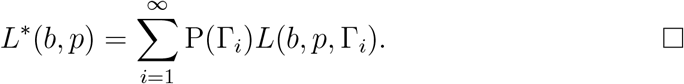

**Remark 4.4.25**. *Evidently the previous formula is applicable for finite partitioning as well (since* P(∅) = 0).

The new life-level function is the weighted average of the expected values of survival times of properties from **𝒫** using the associated weight function *w*.

##### Definition 4.4.26.

*Let a propertized life-level-base-system* ⟨Λ_*b*_, **𝒫***w*, ∝⟩ *be given. Let b* = (*O, K, G, t*) *be an admissible bio-bit. Let a* 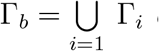 Γ_*i*_ *be acountable disjoint partitioning of* Γ_*b*_ *and let a probability function* P *be given on it. Set*

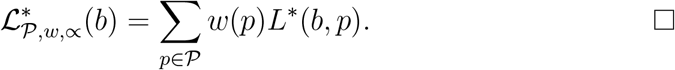

Now we define a restriction of the original **ℒ** _***𝒫***,*w,∝*_ that will be useful later.

##### Definition 4.4.27.

*Let a propertized life-level-base-system* ⟨Λ_*b*_, **𝒫**, *w*, ∝⟩ *be given. Let b* = (*O, K, G, t*) *be an admissible bio-bit. Let* Γ^*′*^ ⊂ Γ. *Set*

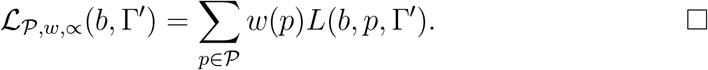

**Remark 4.4.28**. *We remark that both life-levels* 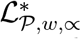 (*b*) *and* **ℒ** _***𝒫***,*w,∝*_(*b*, Γ^*′*^) *can be thought heuristically as time. It is a kind of survival time*.

##### Theorem 4.4.29.

*Let a propertized life-level-base-system* ⟨Λ_*b*_, ***𝒫*** *w*, ∝⟩ *be given. Let b* = (*O, K, G, t*) *be an admissible bio-bit. Let a* 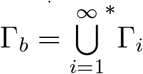 Γ_*i*_ *be a countable disjoint partitioning of* Γ_*b*_ *and let a probability function* P *be given on it. Then*

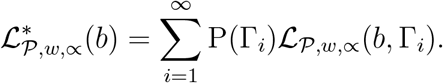

*Proof*. By definition

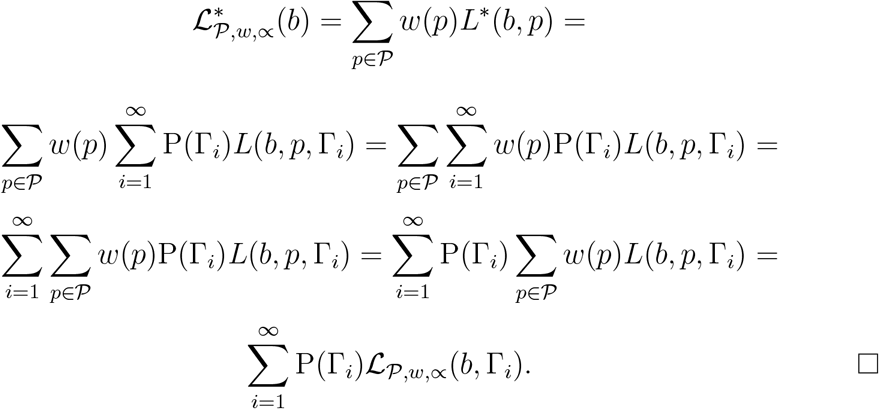

##### Example 4.4.30.

*According to that model, let us calculate the life-level of a man who is at the sate of clinical death. Consider two sets of development-paths* Γ_1_ *and* Γ_2_. *Both sets provide several options on the possible outcomes, but all development-paths of the first set lead to death in a few minutes, all development-paths of the second one provide complete recovery. Let t denote the current moment, let b* = (*O, K, G, t*) *denote the bio-bit in question*.

*Let* P(Γ_1_) = 0.99, P(Γ_2_) = 0.01.

*Let ℒ* _***𝒫***,*w,∝*_(*b*, Γ_1_) = 10 *and ℒ* _***𝒫***,*w,∝*_(*b*, Γ_2_) = 10^7^ *(e.g. one can get figures like these if one measures time in minutes or similar scale)*.

*By 4.4.29* 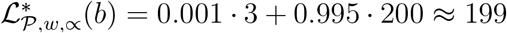 *i.e. it is 2 magnitudes less in the logarithmic scale*.

##### Example 4.4.31.

*Let us enhance the previous example with a third set of development-paths* Γ_3_ *whose development-paths lead to substantial and permanent damage to health. Let us have the following probabilities (we redefine it for* Γ_1_): P(Γ_1_) = 0.89, P(Γ_2_) = 0.01, P(Γ_3_) = 0.1. *Let* **ℒ** _***𝒫***,*w,∝*_(*b*, Γ_3_) = 10^6^.

*Then by 4.4.29 we end up with* 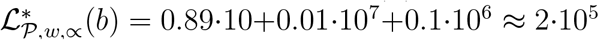

##### Example 4.4.32.

*According to that model, let us calculate the life-level of a dormant tree. Consider two sets of development-paths* Γ_1_ *and* Γ_2_. *The first one leads to death in a few months by some reason related to winter. The second one leads to the usual recovery in spring. Let* P(Γ_1_) = 0.001, P(Γ_2_) = 0.999. *Let* **ℒ** _***𝒫***,*w,∝*_(*b*, Γ_1_) = 3 *and* **ℒ** _***𝒫***,*w,∝*_(*b*, Γ_2_) = 200 *(one measures time in months). By 4.4.29* 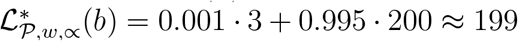 *i.e. almost the same value that we could have got for a tree in the summer e.g*.

##### Example 4.4.33.

*Similarly let us estimate the life-level of a dormant seed. Consider three sets of development-paths* Γ_1_, Γ_2_ *and* Γ_3_. *The first one leads to death in a few decades, no germination here. The second one leads to germination, but the plant does not reach fertility, it dies before. At the third one, it goes through the full usual development of the plant. Let* P(Γ_1_) = 0.9, P(Γ_2_) = 0.09, P(Γ_3_) = 0.01. *Let* **ℒ** _***𝒫***,*w,∝*_(*b*, Γ_1_) = 20, **ℒ**_***𝒫***,*w,∝*_(*b*, Γ_2_) = 21, **ℒ** _***𝒫***,*w,∝*_(*b*, Γ_3_) = 10000 *(we measure time in years, and in* Γ_3_ *we take into account the successors too). By 4.4.29* 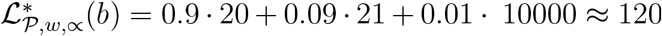

##### Example 4.4.34.

*Suppose that an antelope has high fever causing by foot-and-mouth disease. Consider three sets of development-paths* Γ_1_, Γ_2_ *and* Γ_3_. *The first one leads to death by the disease in a few days. The second one is about that predators can catch it much easier because of its current much weaker conditions, hence it also leads to death. The third one leads to full recovery. Let* P(Γ_1_) = 0.7, P(Γ_2_) = 0.2, P(Γ_3_) = 0.1. *Let* **ℒ** _***𝒫***,*w,∝*_(*b*, Γ_1_) = 4, **ℒ**_*P,w,∝*_(*b*, Γ_2_) = 7, **ℒ** _***𝒫***,*w,∝*_(*b*, Γ_3_) = 3000 *(we measure time in days, and in* Γ_3_ *we do NOT take into account the successors). By 4.4.29* 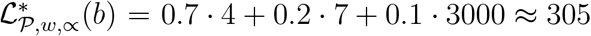. *(cf. example 4.2.6)*

##### Definition 4.4.35.

*Let a life-level-base-system* Λ_*b*_ *be given and b*_1_ = (*O*_1_, *K*_1_, *G*_1_, *t*_1_), *b*_2_ = (*O*_2_, *K*_2_, *G*_2_, *t*_2_) *be two admissible bio-bits such that t*_1_ *< t*_2_. *Then the probability that b*_1_ *can develop into b*_2_ *(i.e. the probability of the event:* ∃*γ* Γ ∈ *γ*(*t*_1_) = (*O*_1_, *K*_1_, *G*_1_), *γ*(*t*_2_) = (*O*_2_, *K*_2_, *G*_2_)*) is denoted by*

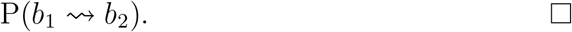

##### Proposition 4.4.36.

*Let a life-level-base-system* Λ_*b*_ *be given and b*_1_ = (*O*_1_, *K*_1_, *G*_1_, *t*_1_), *b*_2_ = (*O*_2_, *K*_2_, *G*_2_, *t*_2_) *be two admissible bio-bits such that t*_1_ *< t*_2_. *Let a* 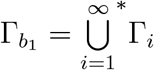 Γ_*i*_ *be a countable disjoint partitioning of* 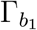 *and let a probability function* P *be given on it. Then*

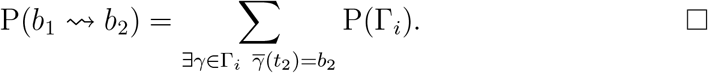

Now we are going to provide formulas how to calculate 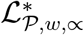 using values at a later point of time i.e. we want a connection between its current value and its values at a future point in time.

##### Lemma 4.4.37.

*Let a propertized life-level-base-system* ⟨Λ_*b*_, **𝒫**, *w*, ∝⟩ *be given. Let p* ∈ **𝒫** *and t*_1_, *t*_2_ ∈ **𝒯**, *t*_1_ *< t*_2_ *and b*_1_ = (*O*_1_, *K*_1_, *G*_1_, *t*_1_) *be an admissible bio-bit. Let a* 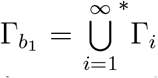 Γ_*i*_ *be a countable disjoint partitioning of* 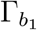 *and let a probability function* P *be given on it. Suppose that if* P(*b*_1_ ⇝ *b*_2_) *>* 0, *γ* ∈Γ, 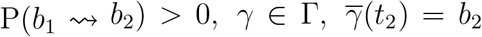 *then* 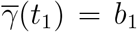. *Also suppose that γ, γ*^*′*^∈ Γ_*i*_ *implies that γ*(*t*_2_) = *γ*^*′*^(*t*_2_). *Then*

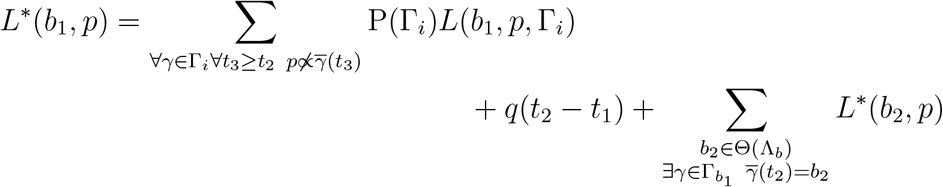

*where*

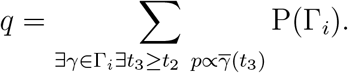

*Proof*. Clearly

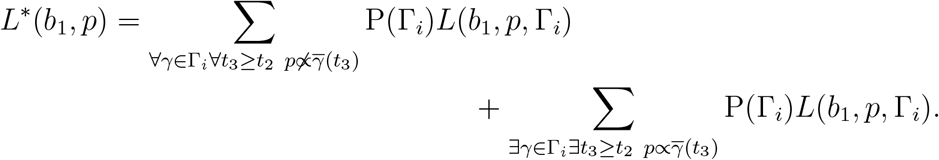

Let us denote the second term by *T*. Then

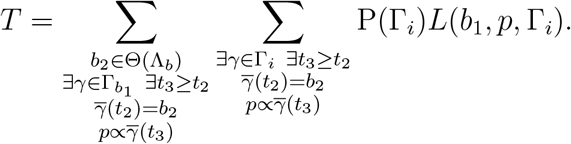

Now using the conditions of the lemma, it can be readily seen that

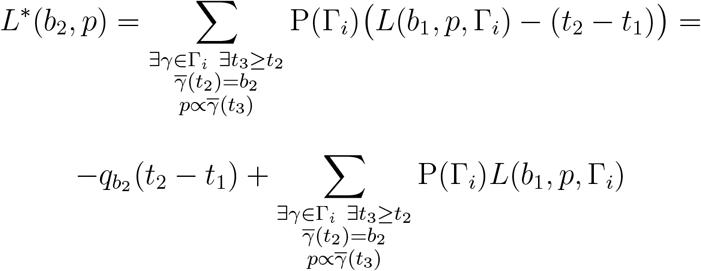

Where

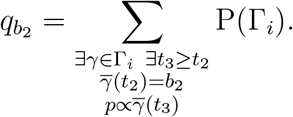

Hence we get that

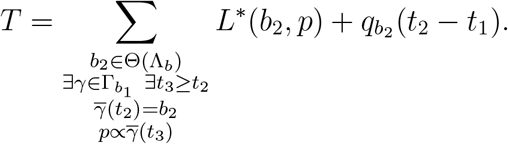

Evidently

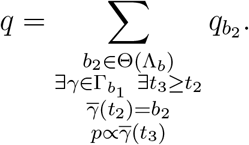

Moreover

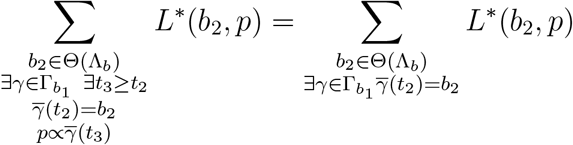

because for the new terms *L*^*\**^(*b*_2_, *p*) = 0.

##### Theorem 4.4.38.

*Let a propertized life-level-base-system* ⟨Λ_*b*_, **𝒫**, *w*, ∝⟩ *be given. Let t*_1_, *t*_2_ ∈**𝒯**, *t*_1_ *< t*_2_ *and b*_1_ = (*O*_1_, *K*_1_, *G*_1_, *t*_1_) *be an admissible bio-bit. Let a* 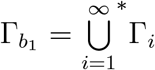 Γ_*i*_ *be a countable disjoint partitioning of* 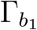 *and let a probability function* P *be given on it. Suppose that if* P(*b*_1_ ⇝ *b*_2_) *>* 0, *γ* ∈Γ, 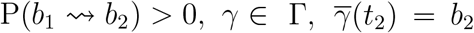 *then* 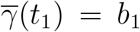. *Also suppose that γ, γ*^*′*^∈ Γ_*i*_ *implies that γ*(*t*_2_) = *γ*^*′*^(*t*_2_). *Then*

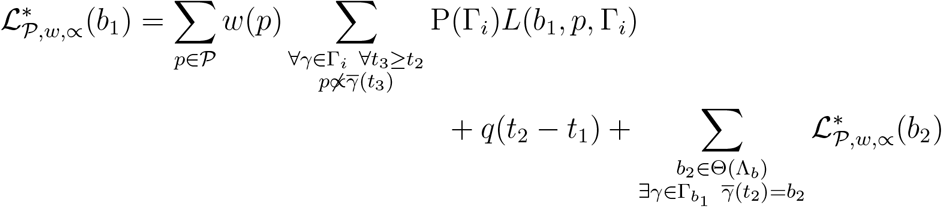

*where*

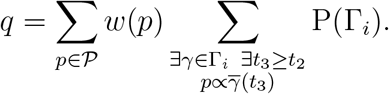

*Proof*. By 4.4.37 we get that

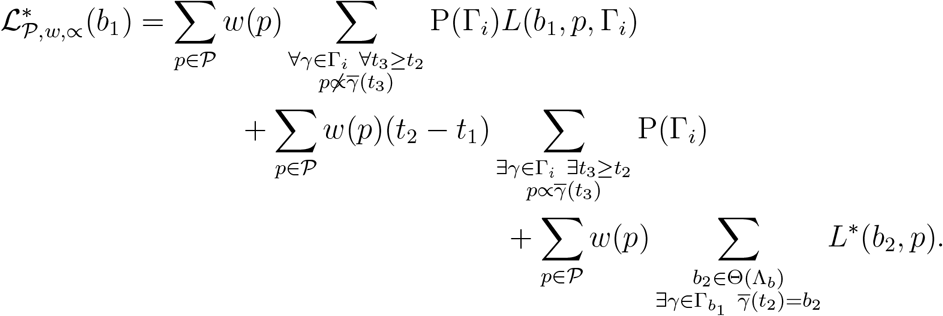

For the last term we get that

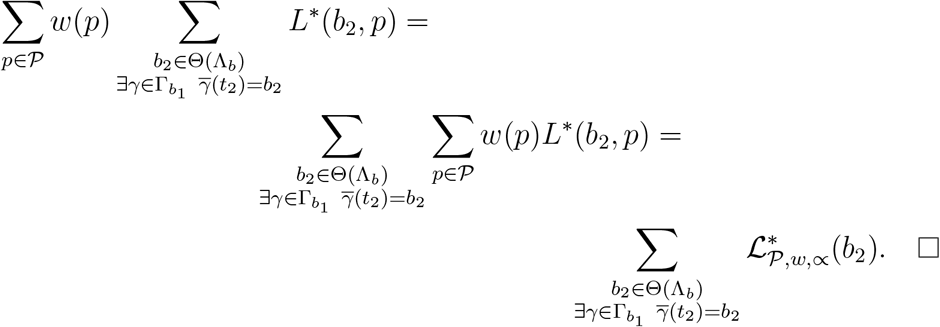

##### Corollary 4.4.39.

*Let a propertized life-level-base-system* ⟨Λ_*b*_, **𝒫**, *w*, ∝⟩ *be given. Let t*_1_, *t*_2_ ∈**𝒯**, *t*_1_ *< t*_2_ *and b*_1_ = (*O*_1_, *K*_1_, *G*_1_, *t*_1_) *be an admissible bio-bit. Let a* 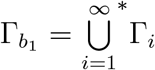 Γ_*i*_ *be a countable disjoint partitioning of* 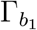 *and let a probability function* P *be given on it. Suppose that if* P(*b*_1_ ⇝ *b*_2_*>* 0, *γ* ∈ Γ, 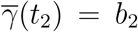 *then γ*(*t*_1_) = *b*_1_. *Also suppose that γ, γ*^*′*^ ∈ Γ_*i*_ *implies that γ*(*t*_2_) = *γ*^*′*^(*t*_2_). *Moreover let* ∀*p* ∈ **𝒫** ∀*i* ∈ ℕ ∃*γ* ∈ Γ_*i*_ ∃*t*_3_ ≥ *t*_2_ *p* ∝ 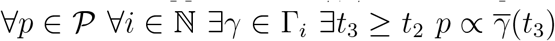 *hold i.e. every property survives till t*_2_ *on each* Γ_*i*_. *Then*

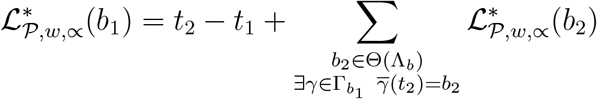

*Proof*. Evidently the first term in 4.4.38 vanishes and

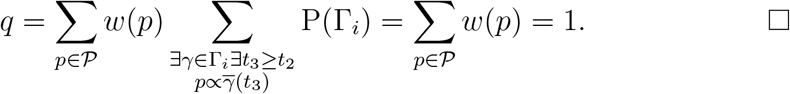

## 5. Algebraic operations on life-level-systems

In the next three sections, we will investigate various operations on life-level functions and systems. First, we are going to examine life-level-systems from some algebraic point of view. Second, we will focus on life-level functions only, and see some basic, technical operations on them, and also some operations that add (restore) nice (expectable) features. And third, we will extend or modify life-level functions / systems according to some biological need and purpose. Hence, while the first two sections are mainly technical, the third section answers biological requirements by partially applying the results of the first two.

This section is about the investigation of algebraic operations on life-level-systems. Here we want to highlight three of them only (actually the first two are the two aspects of the same underlying problem).

1. One of our definitions is about the extension of life-level-systems, i.e. when we extend the domains of our life-level function.
2. We will also examine the merge of two life-level-systems.
3. We will define the equivalence of life-level-systems and also biologically relevant mappings between such systems.

No doubt, they are of utmost interest in biological applications.

### 5.1 Extensions of life-level-systems

First we define the extension of a life-level-system, that is the extended system is greater in every respect, in every component, and also the life-level function is an extension of the original one. In some other sense, we define a natural partial order on life-level-systems at the same time.

#### Definition 5.1.1.

*Let* Λ = ⟨⟨**𝒪** ⊂⟩ ⟨**𝒦** ⊂⟩*ℊ* ***𝒯***Θ, Γ, *ℒbe a life-level-system. Then let* Base(Λ) *denote its underlying life-level-base-system that is* ⟨**𝒪**, ⊂⟩, ⟨**𝒦**, ⊂⟩, ℊ, **𝒯**, Θ, Γ. *ℒ*

#### Definition 5.1.2.

*Let* Λ_1_ = ⟨⟨**𝒪**_1_, ⊂⟩, ⟨**𝒦** _1_, ⊂⟩, ℊ_1_, **𝒯**_1_, Θ_1_, Γ_1_ *and* Λ_2_ ⟨⟨**𝒪**_2_, ⊂⟩, ⟨**𝒦** _2_, ⊂⟩, ℊ_2_, **𝒯**_2_, Θ_2_, Γ_2_ *be two life-level-base-systems. We say that* Λ_2_ *is* ***finer*** *than* Λ_1_, *or* Λ_1_ *is* ***coarser*** *than* Λ_2_, *or* Λ_2_ *is an* ***extension of*** Λ_1_, *or* Λ_1_ *is a* ***restriction of*** Λ_2_, *if*

1. **𝒯**_1_ ⊂ **𝒯**_2_
2. ∀*t* ∈ **𝒯**_1_ **𝒪**_1_(*t*) ⊂ **𝒪**_2_(*t*)
3. ∀*t* ∈ **𝒯**_1_ *the partial order on* **𝒯**_1_(*t*) *equals to the restriction of the partial order on* **𝒪**_2_(*t*) *to* **𝒪**_1_(*t*)
4. ∀*t* ∈ **𝒯**_1_ **𝒦**_1_(*t*) ⊂ **𝒦**_2_(*t*)
5. ∀*t* ∈ **𝒯**_1_ *the partial order on* **𝒦**_1_(*t*) *equals to the restriction of the partial order on* **𝒦**_2_(*t*) *to* **𝒦**_1_(*t*)
6. ∀*t* ∈ **𝒯**_1_ ℊ_1_(*t*) ⊂ ℊ_2_(*t*)
7. Θ_1_ ⊂ Θ_2_
8. ∀*γ*_1_ ∈ Γ_1_ ∃*γ*_2_ ∈ Γ_2_ *such that* ∀*t* ∈ **𝒯**_1_ *γ*_2_(*t*) = *γ*_1_(*t*) *(i.e*. 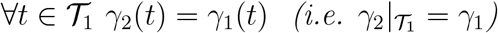). *Furthermore, if* Λ_1_, Λ_2_ *are life-level-systems with associated life-level functions* **ℒ**_1_ *and* **ℒ**_2_ *respectively, then we also require condition*
9. Dom ℒ_1_ ⊂ Dom ℒ_2_ *and* 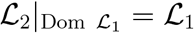.

*In both cases (life-level-base-system and life-level-system), we will also use the notation* Λ_1_ ≤ Λ_2_. □

#### Proposition 5.1.3.

*The extension relation* ≤ *is a partial order on the sets of all life-level-base-systems and life-level-systems*. □

**Remark 5.1.4**. *If* **𝒯**_1_ = **𝒯**_2_, *then the condition on the development-paths is equivalent with* Γ_1_ ⊂ Γ_2_. *While if* **𝒯**_1_ **𝒯**_2_, *then it requires that each* Λ_1_*-development-path has an extension in* Λ_2_ *regarding time*.

#### Example 5.1.5.

*Suppose one has a life-level-system for describing some part of a given ecosystem. We enumerate some possible ways how one can get an extension. One gets an extension if*

- *One adds another ecosystem to one’s model (or just some part of it; but it is a different ecosystem)*.
- *One adds gorganisms to the model that are on greater hierarchical level than already existed in the model, but they live in the same ecosystem*.
- *E.g. until now we just investigated normal organisms, but from now on, we want to examine some groups too*.
- *Similarly for smaller hierarchical level. E.g. we investigated some multi-cell organisms, but now we want to examine the building block cells as well*.
- *One simply adds more gorganisms / sub-environments / supporting-groups to the model*.
- *One adds new development-paths to the model*.
- *One defines life-level on bio-bits where it has not been defined yet*.
- *One refines the time-line by adding more measurement time points to the model (and extends the already existing development-paths according to the new time line)*.

*Of course, one of the above points gives an extension, but they can be combined too*. □

Two natural examples for a coarser life-level-system are the followings.

#### Proposition 5.1.6.

*Let* Λ = ⟨⟨**𝒪**, ⊂⟩, ⟨**𝒦**, ⊂⟩, ℊ, **𝒯**, Θ, Γ, **ℒ** *be a life-level-system in the gene ral con text, and let O* ∈ **𝒪**. *Then* 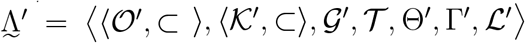 *is a life-level-system too, where*

- **𝒪***′* = {*O*^*′*^ ∈ **𝒪** : *O*^*′*^ ⊂ *O*}, *and the partial order on* **𝒪***′ equals to the restriction of the partial order on* **𝒪** *to* **𝒪***′*
- Θ^*′*^ = {*b* ∈ Θ : *π*_***𝒪***_ (*b*) ∈ **𝒪***′*}
- Γ^*′*^ = {*γ* ∈ Γ : ∀*t* ∈ **𝒯** 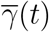 ∈ Θ^*′*^}
- **𝒦***′* = {*π*_***𝒦***_ (*b*) : *b* ∈ Θ^*′*^}, *and the partial order on* **𝒦***′ equals to the restriction of the partial order on* **𝒦** *to* **𝒦***′*
- **𝒢***′* = {*π*_*ℊ*_ (*b*) : *b* ∈ Θ^*′*^}
- Dom ℒ*′* = Dom **ℒ** ∩ Θ^*′*^, ℒ*′* = ℒ|_Dom_ ℒ*′*.

#### Proposition 5.1.7.

*Let* Λ = ⟨**𝒪**, ⊂⟩, ⟨**𝒦**, ⊂⟩, **𝒢, 𝒯**, Θ, Γ, **ℒ** *be a life-level-system, and let γ* ∈ Γ. *Then* Λ^*′*^ = ⟨⟨**𝒪***′*, ⊂⟩, ⟨**𝒦***′*, ⊂⟩, ⟨**𝒢***′*⟩, **𝒯**, Θ^*′*^, Γ^*′*^,**ℒ***′* ⟩is *a life-level-system too, where*

- ∀*t* ∈ **𝒯 𝒪***′*(*t*) = {***𝒪***^*′*^ ∈ **𝒪** (*t*) : ***𝒪***^*′*^ ⊂ *γ*_***𝒪***_ (*t*)}, *and the partial order on* **𝒪***′*(*t*) *equals to the restriction of the partial order on* **𝒪** (*t*) *to* **𝒪***′*(*t*)
- Θ^*′*^ = {*b* ∈ Θ : *π*_***𝒯***_ (*b*) ∈ **𝒯***′*}
- Γ^*′*^ = {*γ*^*′*^ ∈ Γ : ∀*t* ∈ **𝒯** 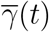 ∈ Θ^*′*^}
- **𝒦***′* = {*π*_***𝒦***_ (*b*) : *b* ∈ Θ^*′*^}, *and* ∀*t* ∈ **𝒯** *the partial order on* **𝒦***′*(*t*) *equals to the restriction of the partial order on* **𝒦** (*t*) *to* **𝒦***′*(*t*)
- **𝒢***′* = {*π*_***𝒢***_ (*b*) : *b* ∈ Θ^*′*^}
- Dom **ℒ***′* = Dom **ℒ** ∩ Θ^*′*^,**ℒ***′* =ℒ|_Dom *ℒ*_*′*.

#### Definition 5.1.8.

*The life-level-systems determined in the previous propositions 5.1.6, 5.1.7 will be denoted by* 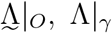 *and called the restriction of* 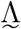 *to the gorganism O, and the restriction of* Λ *to the development-path γ respectively*.

Now we define the infimum of two life-level-systems.

#### Definition 5.1.9.

*Let* Λ_1_ = ⟨⟨**𝒪**_1_, ⊂⟩, ⟨**𝒦**_1_, ⊂⟩, **𝒢**_1_, **𝒯**_1_, Θ_1_, Γ_1_, **ℒ**_1_ *and* Λ_2_ = ⟨**𝒪** _2_, ⊂⟩, ⟨**𝒦**_2_, ⊂⟩, **𝒢**_2_, **𝒯**_2_, Θ_2_, Γ_2_, **ℒ** _2_ *be two life-level-systems. The system* Λ ⟨⟨**𝒪**, ⊂⟩, ⟨**𝒦**, ⊂⟩, **𝒢, 𝒯**, Θ, Γ, **ℒ** *is the* ***infimum*** *of* Λ_1_ *and* Λ_2_, *if*

1. **𝒯** = **𝒯**_1_ ∩ **𝒯**_2_
2. ∀*t* ∈ **𝒯 𝒪** (*t*) = **𝒪**_1_(*t*) ∩ **𝒪**_2_(*t*)
3. ∀*t* ∈ **𝒯** *the partial order on* **𝒪** (*t*) *equals to the restriction of the partial order on* **𝒪**_*i*_(*t*) *to* **𝒪**(*t*) *(i* = 1, 2)
4. ∀*t* ∈ **𝒯 𝒦** = **𝒦**_1_(*t*) ∩ **𝒦**_2_(*t*)
5. ∀*t* ∈ **𝒯** *the partial order on* **𝒦** (*t*) *equals to the restriction of the partial order on* **𝒦**_*i*_(*t*) *to* **𝒦**(*t*) *(i* = 1, 2)
6. ∀*t* ∈ **𝒯 𝒢** (*t*) = **𝒢**_1_(*t*) ∩ **𝒢**_2_(*t*)
7. Θ = Θ_1_ ∩ Θ_2_
8. Dom **ℒ** = {*b* ∈ Dom **ℒ**_1_ ∩ Dom **ℒ**_2_ : **ℒ**_1_(*b*) = **ℒ**_2_(*b*)}
9. Γ = *γ* : **𝒯** → **𝒪** × **𝒦** × **𝒢** | ∃*γ*_1_ ∈ Γ_1_ ∃*γ*_2_ ∈ Γ_2_ *such that* ∀*t* ∈ **𝒯** *γ*(*t*) = *γ*_1_(*t*) = *γ*_2_(*t*) *and* 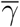(*t*) ∈ Dom **ℒ**

*In this case we will also use the notation* Λ = Λ_1_ ∩ Λ_2_. □

**Remark 5.1.10**. *In definition 5.1.9 actually the only strict requirements are the last two. Criterion (8) requires that* Dom **ℒ** *is the greatest set where* **ℒ**_1_ *and* **ℒ**_2_ *are coincide, i.e*. ℒ_1_|_Dom_ ℒ = ℒ_2_|_Dom_ ℒ. *While criterion (9) identifies all development-paths where* _1_ *and* _2_ *take the same values at every time point. Of course one can easily take the intersection of the sets of gorganisms, sub-environments, supporting groups, time, bio-bits and development-paths, but the last two criterions require more than a simple intersection*.

#### Proposition 5.1.11.

*If* Λ_1_ *and* Λ_2_ *are two life-level-systems, then so is* Λ_1_ ∩ Λ_2_. *Both* Λ_1_ *and* Λ_2_ *are extensions of* Λ_1_ ∩ Λ_2_, *i.e*. Λ_1_ ∩ Λ_2_ ≤ Λ_1_, Λ_2_.

*Proof*. The two requirements on development paths and life-level function properties (es), (do) and (ss) can be easily verified. The statement on extension is obvious. □

#### Proposition 5.1.12.

*If* Λ_1_ *and* Λ_2_ *are two life-level-systems, then their infimum* Λ_1_ ∩ Λ_2_ *always exists, moreover it is the greatest life-level-system* Λ *such that* Λ ≤ Λ_1_, Λ_2_. □

#### Example 5.1.13.

*If two research projects both have a life-level-system, then they can identify the greatest part which is common in both. That is going to be the infimum*.

One can easily define the infimum of arbitrarily many life-level-systems according to definition 5.1.9, and then prove similar statements too.

#### Definition 5.1.14.

*Let* Λ_*i*_ = ⟨⟨**𝒪**_*i*_, ⊂⟩, ⟨**𝒦**_*i*_, ⊂⟩, **𝒢**_*i*_, **𝒯**_*i*_, Θ_*i*_, Γ_*i*_, **ℒ**_*i*_ *be life-level-systems for i* ∈ *I. The system* Λ ⟨⟨**𝒪**, ⊂⟩, ⟨**𝒦**, ⊂⟩, **𝒢, 𝒯**, Θ, Γ, **ℒ** is *the* ***infimum*** *of them, if similar statements hold from points (1) to (8) in definition 5.1.9 with ∩* _*i∈I*_ *instead of intersection for two sets. We formalize (9) only:*

*9*. Γ = *γ* : **𝒯** → **𝒪** × **𝒦** × **𝒢** | ∀*i* ∈ *I* ∃*γ*_*i*_ ∈ Γ_*i*_ *such that* ∀*t* ∈ **𝒯** ∀*i* ∈ *I γ*(*t*) = *γ*_*i*_(*t*) *and γ*(*t*) ∈ Dom **ℒ**

*In this case we will use the notation* 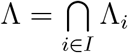 □

#### Proposition 5.1.15.

*Let* Λ_1_, Λ_2_, Λ_3_ *be life-level-systems, such that* Λ_1_ ≤ Λ_3_, Λ_2_ ≤ Λ_3_. *Then there is a coarsest life-level-system that is finer than both* Λ_1_ *and* Λ_2_.

*Proof*. Take the infimum of all life-level-systems that are finer than both. □

#### Definition 5.1.16.

*Let* Λ_1_, Λ_2_ *be life-level-systems, such that there is a finer life-level-system than both. Then the coarsest life-level-system that is finer than both* Λ_1_ *and* Λ_2_ *will be called the* ***supremum of*** Λ_1_ *and* Λ_2_, *and we will use the notation* Λ_1_ ∪ Λ_2_. □

**Remark 5.1.17**. *For two given life-level-systems* Λ_1_, Λ_2_, *it is not necessary that there is a life-level-system that is an extension of both. Clearly a necessary condition for that is that* 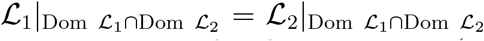. *But it is not sufficient. It is easy to create an example where property (ss) would be violated in any extension*.

**Remark 5.1.18**. *Suppose that two research projects investigate the same ecosystem from life-level perspective, but one examines the organisms and smaller hierarchical levels, while the other examines groups of organisms and higher hierarchical levels. Then they want to merge their life-level-systems. With the terminology built here, it is possible if there is a common extension of both systems*.

*Here the main difficulty can be to guarantee property (ss). One possible way can be to move upwards all life-level values of the second system (which operates on higher levels) by the supremum of all values of the first system*.

### 5.2 Morphisms

Here we are going to investigate mappings between life-level-systems and also the relations that those mappings may express, and relations that can be formalized by properties of some mappings.

#### Definition 5.2.1.

*Let* Λ_1_ = ⟨⟨**𝒪**_1_, ⊂⟩, ⟨**𝒦**_1_, ⊂⟩, **𝒢**_1_, **𝒯**_1_, Θ_1_, Γ_1_⟩ *and* Λ_2_ =⟨⟨**𝒪**_2_, ⊂⟩, ⟨**𝒦**_2_, ⊂⟩, **𝒢**_2_, **𝒯**_2_, Θ_2_, Γ_2_ ⟩*be two life-level-base-systems. A vector of functions F* = ⟨*F*_*𝒪*_, *F*_***𝒦***_, *F*_***𝒢***_, *F*_***𝒯***_ ⟩ *is called a* ***morphism*** *between* Λ_1_ *and* Λ_2_, *if*

1. *F*_**𝒯**_ : **𝒯**_1_ → **𝒯**_2_ *is continuous and strictly increasing*
2. *F*_*𝒪*_ : **𝒪**_1_ → **𝒪**_2_ *is a function such that* ∀*t* ∈ 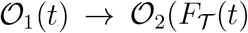 *that keeps the partial order i.e. if O, O*^*′*^ ∈ **𝒪**_1_(*t*) *and O* ⊂ *O*^*′*^, *then F*_***𝒪***_ (*O*) ⊂ *F*_***𝒪***_ (*O*^*′*^).
3. *F*_***𝒦***_ : **𝒦**_1_ → **𝒦**_2_ *is a function such that* ∀*t* ∈ **𝒯**_1_ 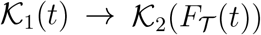 **𝒦**_1_(*t*) → **𝒦**_2_(*F*_***𝒯***_ (*t*)) *that keeps the partial order i.e. if K, K*^*′*^ ∈ **𝒦**_1_(*t*) *and K* ⊂ *K*^*′*^, *then F*_***𝒦***_ (*K*) ⊂ *F*_*𝒦*_(*K*^*′*^).
4. *F* _***𝒢***_ : **𝒢**_1_ → **𝒢**_2_ *is a function such that* ∀*t* ∈ **𝒯**_1_ 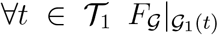 **𝒢**_1_(*t*) → **𝒢**_2_
5. (*F*_*𝒯*_ (*t*)) *if b* = (*O, K, G, t*) ∈ Θ _1_, *then F* (*b*) = *F* (*O, K, G, t*) ∈ Θ_2_, *where F* (*b*) = *F* (*O, K, G, t*) = *F* _***𝒪***_ (*O*), *F* _***𝒦***_ (*K*), *F* _***𝒢***_ (*G*), *F* _***𝒯***_ (*t*) ; *we will use the notation F* : Θ_1_ → Θ_2_
6. *if γ* ∈ Γ_1_, *then F* ° *γ* ° *F* ^*−*1^ ∈ Γ_2_ *(note that* 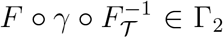 **𝒯**_2_ → Θ_2_ *as expected); we will use the notations F* : Γ_1_ → Γ_2_ *and* 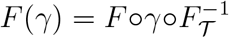

*Furthermore, if* Λ_1_, Λ_2_ *are life-level-systems with associated life-level functions* **ℒ**_1_ *and* **ℒ**_2_ *respectively, then we also require*

7.a. *if b* ∈ Dom **ℒ**_1_, *then F* (*b*) ∈ Dom **ℒ**_2_.
7.b. *We call F* ***id-morphism*** *if* ℒ_1_ *and* ℒ_2_ ? *F are equivalent life-level functions, i.e. there is a strictly increasing function* 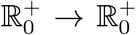 *with f* (0) = 0, *such that b* ∈ Dom **ℒ** _1_ *implies that* **ℒ**_1_(*b*) = *f* **ℒ**_2_ *F* (*b*))). *If instead* **ℒ**_1_(*b*) ≤ *f* **ℒ**_2_ *F* (*b*)))*holds, then F is called* ***increasing-morphism***; *if ℒ*_1_(*b*) *f* **ℒ**_2_ *F* (*b*))) *holds, then F is called* ***decreasing-morphism***.

*We will also use the notations that F* : Λ_1_ → Λ_2_, *and say that F is a morphism*.

*We call F* : Λ_1_ → Λ_2_ *an* ***epimorphism***, *if F is a morphism, F*_***𝒯***_ *is surjective, and* ∀*t* ∈ **𝒯**_1_ 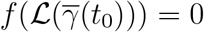 *are surjective mappings, F* : Θ_1_ → Θ_2_ *is surjective and F* : Γ_1_ → Γ_2_ *is also surjective*.

*We call F* : Λ_1_ → Λ_2_ *an* ***isomorphism***, *if F is a morphism, F*_***𝒯***_ *is bijective, and* ∀*t* ∈ **𝒯**_1_ 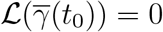 *are bijective mappings, F* : Θ_1_ → Θ_2_ *is bijective and F* : Γ_1_ → Γ_2_ *is also bijective*.

*We call F* : Λ_1_→ Λ_2_ ***strong*** *(id- / increasing- / decreasing-morphism) if instead of condition 7.b*., *the following holds*

*7.b’*. ℒ_1_ = ℒ_2_ ?*F*, ℒ_1_ ≤ ℒ_2_? *F*, ℒ_1_ ≥ ℒ_2_?*F, i.e. if b* ∈ Dom **ℒ**_1_, *then* **ℒ**_1_(*b*) = ℒ_2_ *F* (*b*), ℒ_1_(*b*) ≤ ℒ_2_ *F* (*b*), ℒ_1_(*b*) ≥ ℒ_2_ *F* (*b*) *respectively*.

**Remark 5.2.2**. *In the definition 5.2.1 we used the symbol F in three senses: F* : Λ_1_→ Λ_2_, *F* : Θ_1_→ Θ_2_ *and F* : Γ_1_ → Γ_2_. *This might be ambiguous slightly, but the context should define which one is meant currently. Of course, F* : Λ_1_ → Λ_2_ *is the “main” one and the other two derived from it naturally*.

**Remark 5.2.3**. *If two systems are isomorphic, then it means that they work equivalently from life-level perspective. They may have totally different elements (gorganisms, sub-environments, supporting groups, development-paths), but there is a one-to-one correspondence among the elements that keeps life-level everywhere*.

*If one system is an epimorphic image of another one, then the source system is richer in the sense that there are more elements, but the elements can be grouped in a way such that from life-level perspective it works in the same way*.

**Remark 5.2.4**. *The definition 5.2.1 contains the possibility that there is a strong correspondence and equivalence between two life-level-systems but not on the same time scale, in other words, one system lives “faster” than the other, but apart from that, they work in the same way from life-level perspective. If we speeded up time in the second one, then we would get the same life-level structures and values as we have in the first one*.

#### Example 5.2.5.

*Compare some surface bacteria to similar not-dormant bacteria living in the subsurface which live where there are serious energy limitations constantly, hence the metabolism of the latter bacteria is much much slower than the first one (see [13].)*. *This can be an example of a morphism where only the mapping of time is not a bijection, everything else is*.

#### Example 5.2.6.

*Two distinct ecosystems of two distinct forests located close to each other can be isomorphic in the sense that we can map the elements (gorganisms, sub-environments, supporting groups, development-paths) to the corresponding elements of the other in a way that keeps the life-level values. Here the mapping can be straightforward: we map the elements of similar “types” to each other, e.g. an adult male rabbit from one forest is mapped to an adult male rabbit from the other forest; also the corresponding surrounding of the first rabbit where it lives is mapped to the corresponding surrounding of the other rabbit; similarly for their supporting group; and finally the possible development-paths of the first rabbit are mapped to development-paths of the other rabbit from the other forest*.

#### Example 5.2.7.

*Suppose that our life-level-systems consists of several components of organisms, i.e. there is a given organism and we “look” just inside the organism, all components are part of it. Then two such systems belonging to “very similar” such organisms can be isomorphic. E.g. the life-level-systems belonging to the body of two health adult male gorillas are most probably isomorphic. Or just take two lungs of a man as full life-level-systems, they are most probably isomorphic*.

#### Example 5.2.8.

*Continuing the previous example, isomorphic systems can belong to very different gorganisms. E.g. a kidney of a man can be isomorphic to a kidney of pig. Or a cell of a plant can be isomorphic to a cell of an animal, e.g. in a way, chloroplasts can be mapped to mitochondrions. Or some part of an ecosystem living among tree roots in the ground can be isomorphic to some part of an ecosystem living among water plants (in the water)*.

*We emphasize, that here, isomorphism is strictly meant from life-level perspective and nothing else*.

**Remark 5.2.9**. *The time line* **𝒯** *has important role here. It can happen that F*_*𝒯*_ *is the identity mapping that means that we do our kind of comparison on life-level-systems at the same time interval, same time frame. However we can also compare life-level-systems existing at different point in time. Then we have to apply a simple shift in time, i.e. F*_***𝒯***_ (*t*) = *t* + *t*_0_ *where t*_0_ *is some fixed value*.

#### Example 5.2.10.

*We can have an ecosystem of a given forest 50 years ago and from life-level perspective we can compare it to the currently existing ecosystem, to the ecosystem that evolved from old one. E.g. they can be isomorphic or the old one can be richer in the sense that the new life-level-system is an epimorphic image of the old one, but strictly containing less components*.

#### Definition 5.2.11.

*Let* Λ_1_ = ⟨**𝒪**_1_, ⊂⟩, ⟨**𝒦**_1_, ⊂⟩, **𝒢**_1_, **𝒯**_1_, Θ_1_, Γ_1_, **ℒ**_1_⟩ *and* Λ_2_ =⟨**𝒪**_2_, ⊂⟩, ⟨**𝒦**_2_, ⊂⟩, **𝒢**_2_, **𝒯**_2_, Θ_2_, Γ_2_, **ℒ**_2_ ⟩ *be two life-level-systems. We say the* Λ_2_ *is a* ***quotient*** *of* Λ_1_ *if there is epimorphism F* : Λ_1_ → Λ_2_.

**Remark 5.2.12**. *What does that mean that* Λ_2_ *is a quotient of* Λ_1_*? It means that the elements (gorganisms, sub-environments, supporting groups) of* Λ_1_ *can be grouped, can be organized into groups such that this grouping keeps the development-paths and also the life-level values in a way that they are totally “synchronized” with the corresponding development-paths and life-level values of* Λ_2_.

#### Example 5.2.13.

*Suppose that in* Λ_1_ *we have tissues, organs and organ systems of some organisms. While in* Λ_2_ *we have organs, organ systems and the corresponding organisms. Both* Λ_1_ *and* Λ_2_ *are based on the same organisms. Let F* : Λ_1_ →Λ_2_ *be defined the way that each gorganism is mapped to its container in the hierarchy, i.e. tissues are mapped to the containing organ, organs are mapped to the containing organ system, organ systems are mapped to the containing organism. Assuming that the sub-environments and the supporting-groups follow a similar synchronized hierarchy, then we can extend F similarly to sub-environments and supporting-groups as well such that we end up with F being a strong-increasing-epimorphism*.

Now we going to define a natural necessary requirement on a life-level-system that is roughly the following. If it has sub-systems that work in the same way, then they should provide same life-level values. E.g. if in a system there are two gorganisms with equivalent inner structure and development-paths, then we expect very similar life-level values for both.

#### Definition 5.2.14.

*We call a life-level-system* Λ ***iso-self-synchronized*** *if* Λ_1_ ≤Λ, Λ_2_ ≤Λ *and* Base(Λ_1_), Base(Λ_2_) *are isomorphic, then* Λ_1_, Λ_2_ *are strong-id-isomorphic*.

#### Example 5.2.15.

*An iso-self-synchronized life-level-system provides the same life-level values for two members of the same species living at the same area, under very similar circumstances, being at the same age, having the same sex, etc*.

## 6. Operations on life-level functions

In this section we are going to present several operations which creates new functions from a given life-level function (or from two or many). Actually the methods/operations described here works for any life-level function i.e. it does not matter how we created/defined the original life-level function(s). In most of the cases the new function will be a life-level function too, however it may also happen that it will not satisfy all properties (es), (do) and (ss), mentioned in definition 2.1.10. Because of that we also provide some (technical) operations which restore some of the missing properties for the new function.

We start with some very basic operations, then present some technical ones, and finally some “meaningful” ones.

### 6.1 Basic operations

#### Proposition 6.1.1.

*Let a life-level-system* Λ *be given, let* **ℒ** *denote its associated life-level function. Let f* : 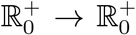 *be a strictly increasing function and f* (0) = 0. *Then f* (**ℒ**) *is also an associated life-level function*.

*Proof*. (es): If **ℒ** (*b*) = 0, then *f* (**ℒ**(*b*)) = 0 (*b* ∈ Dom **ℒ**).

(do): If *γ* ∈ Γ, 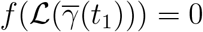, *t*_0_ *< t*_1_, then 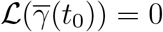 which yields that 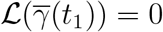 which gives that 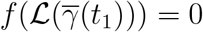.

(ss): If *b*_1_, *b*_2_ ∈ Dom **ℒ**, *b*_1_ ⊆_*𝒪*_ *b*_2_, then **ℒ**(*b*_1_) ≤ **ℒ**(*b*_2_). Then *f* (**ℒ**(*b*_1_)) ≤ *f* (**ℒ**(*b*_2_)) holds. □

#### Proposition 6.1.2.

*Let a life-level-system* Λ *be given, let* **ℒ** *denote its associated life-level function. Let c* ∈ ℝ^+^. *Then c***ℒ**, lg(1 + **ℒ**), 10^*ℒ*^ − 1 *are all associated life-level functions that are equivalent to* **ℒ**.

*If* **ℒ** *<* 1, *then* 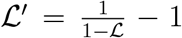 *is an equivalent life-level function whose domain is extended to* [0, +∞).

*If* Dom **ℒ** ⊂ [0, +∞), *then* 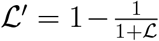 *is an equivalent life-level function such that* **ℒ** *<* 1.

*Proof*. The equivalences are ensured by the functions *f* (*x*) = *cx, f* (*x*) = lg(1 + *x*), 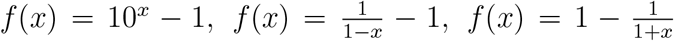 respectively. Showing that they are associated, apply 6.1.1. In the last two cases, the change of the corresponding domain can be checked easily. □

#### Proposition 6.1.3.

*Let a life-level-base-system* Λ_*b*_ *be given together with two associated life-level functions* **ℒ**_1_, **ℒ**_2_ *on a common domain. Let* 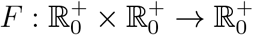 *be a 2-variable function such that*

1. *F* (*x, y*) = 0 ⇔ (*x, y*) = (0, 0) *and*
2. *x* ≤ *x*^*′*^, *y* ≤ *y*^*′*^ *implies that F* (*x, y*) ≤ *F* (*x*^*′*^, *y*^*′*^).

*Then F* (**ℒ**_1_, **ℒ**_2_) *is also an associated life-level function*.

*Proof*. Set **ℒ** = *F* (**ℒ**_1_, **ℒ**_2_).

(es) **ℒ** (∅, *K, G, t*)= 0 obviously holds by the first condition.

(do) If *γ* ∈ Γ, *γ*_*𝒪*_ (*t*_0_)≠∅, **ℒ**_*γ*_(*t*_0_) = 0, *t*_0_ *< t*_1_ then 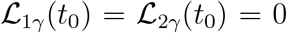 again by the first condition. This implies that 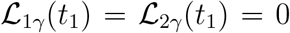 which yields that **ℒ**_*γ*_(*t*_1_) = 0.

(ss) If *b*_1_, *b*_2_ ∈ Dom **ℒ**, *b*_1_ ⊆_*𝒪*_ *b*_2_ then **ℒ**_1_(*b*_1_) ≤ **ℒ**_1_(*b*_2_), **ℒ**_2_(*b*_1_) ≤ **ℒ**_2_(*b*_2_) hold which gives that **ℒ**(*b*_1_) ≤ **ℒ**(*b*_2_) by the second condition. □

#### Proposition 6.1.4.

*Let a life-level-base-system* Λ_*b*_ *be given together with two equivalent associated life-level functions* **ℒ**_1_, **ℒ**_2_ *on a common domain. Let F:* 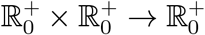 *be a 2-variable function such that*

1. *F* (0, 0) = 0 *and*
2. *x < x*^*′*^, *y < y*^*′*^ *implies that F* (*x, y*) *< F* (*x*^*′*^, *y*^*′*^).

*Then F* (**ℒ**_1_, **ℒ**_2_) *is also an associated life-level function that is equivalent to* **ℒ**_1_ *(and then to* **ℒ**_2_ *too)*.

*Proof*. Let 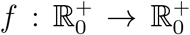 be a strictly increasing function with *f* (0) = 0, such that **ℒ**_2_ = *f* (**ℒ**_1_). To show the equivalence, set 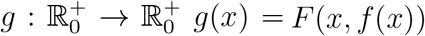. Clearly *g*(**ℒ**_1_) = *F* (ℒ_1_,ℒ_2_). We have to show that *g* is a strictly increasing function and *g*(0) = 0. If *x < x*^*′*^, then *f* (*x*) *< f* (*x*^*′*^) and the second condition gives the first claim. Obviously *g*(0) = *F* (0, *f* (0)) = *F* (0, 0) = 0.

By 6.1.1 *F* (**ℒ**_1_, **ℒ**_2_) is an associated life-level function. □

#### Corollary 6.1.5.

*Let a life-level-base-system* Λ_*b*_ *be given together with two associated life-level functions* **ℒ**_1_, **ℒ**_2_ *on a common domain. Then* **ℒ**_1_ +**ℒ**_2_ *and* **ℒ**_1_ + **ℒ**_1_**ℒ**_2_ *are both associated life-level functions*.

*If* **ℒ**_1_ *and* **ℒ**_2_ *are equivalent then* **ℒ**_1_+**ℒ**_2_ *and* **ℒ**_1_+**ℒ**_1_**ℒ**_2_ *both are equivalent to* **ℒ**_1_ *(and then* **ℒ**_2_ *too)*.

*Proof*. Regarding equivalence, apply 6.1.4 for the functions *F* (*x, y*) = *x* + *y* and *G*(*x, y*) = *x* + *xy* respectively.

Regarding association, *F* (*x, y*) = *x* + *y* works for **ℒ**_1_ + **ℒ**_2_ by 6.1.3. But *G*(*x, y*) = *x* + *xy* does not satisfy the necessity part of condition (1) in 6.1.3, however that part is used in the verification of (do) only, and it holds because **ℒ**_*γ*_(*t*_0_) = 0 implies that 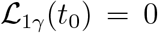which is enough to get that **ℒ**_*γ*_(*t*_1_) = 0. □

Now we formulate a weakening of 6.1.1.

#### Proposition 6.1.6.

*Let a life-level-system* Λ *be given, let* **ℒ** *denote its associated life-level function. Let* 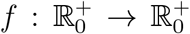*be an increasing function such that f* (*x*) = 0 ⇔ *x* = 0. *Then f* (**ℒ**) *is also an associated life-level function that is alive-equivalent with* **ℒ**.

*Proof*. (es) and (ss) can be similarly verified as in the proof of 6.1.1. In (do), one has to apply that *f* (*x*) = 0 ⇒ *x* = 0. Alive-equivalence follows from *f* (*x*) *>* 0 ⇔ *x >* 0.

#### Proposition 6.1.7.

*Let a life-level-base-system* Λ_*b*_ *be given together with associated life-level functions* **ℒ**_1_, …, **ℒ**_*n*_ *on a common domain* (*n* ∈ N). *Then* max{**ℒ**_1_, …, **ℒ**_*n*_} *is also an associated life-level function*.

*Proof*. Set **ℒ** = max{**ℒ**_1_, …, **ℒ**_*n*_}.

(es) **ℒ** (∅, *K, G, t)* = 0 obviously holds.

(do) If *γ* ∈ Γ, *γ*_*𝒪*_(*t*_0_) ∅, **ℒ**_*γ*_(*t*_0_) = 0, *t*_0_ *< t*_1_ then 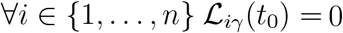 which implies that 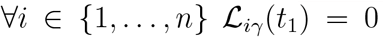 which yields that **ℒ**_*γ*_(*t*_1_) = 0.

(ss) If *b*_1_, *b*_2_ ∈ Dom **ℒ**, *b*_1_ ⊆_*𝒪*_ *b*_2_ then ∀*i* ∈ {1, …, *n*} **ℒ**_*i*_(*b*_1_) ≤ **ℒ**_*i*_(*b*_2_) holds. Let *i*_0_ ∈ {1, …, *n*} such that 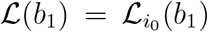. But then 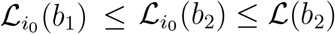 which gives the claim. □

**Remark 6.1.8**. *A similar statement is true for countably many associated life-level functions with* sup *instead of* max.

#### Proposition 6.1.9.

*Let a life-level-base-system* Λ_*b*_ *be given together with a sequence of associated life-level functions* (**ℒ**_*n*_) *on a common domain. If* **ℒ**_*n*_ *→* **ℒ** *(pointwise), then* **ℒ** *is also an associated life-level function, provided that there is δ >* 0 *such that* ∀*n* Ran **ℒ**_*n*_ − {0} ⊂ (*δ*, +∞).

*Proof*. Ran 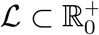, and properties (es) and (ss) are all obvious.

The condition is needed for verifying property (do): If *γ* ∈ Γ, **ℒ**_*γ*_(*t*) = 0, then there is *N* such that *n > N* implies that **ℒ**_*n,γ*_(*t*) = 0. Now if *t*^*′*^ *> t, n > N*, then **ℒ**_*n,γ*_(*t*^*′*^) = 0 which yields that **ℒ**_*γ*_(*t*^*′*^) = 0.

**Remark 6.1.10**. *The condition in the previous proposition can be roughly rephrased that there is “minimal” positive life-level value*. □

### 6.2 Guaranteeing property (do)

We present a simple technical operation that makes a function satisfy property (do) if originally it did not satisfy it.

#### Definition 6.2.1.

*Let a life-level-base system* Λ_*b*_ *be given together with a* Λ_*b*_*-base-function*

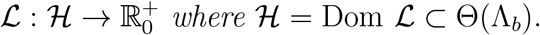

*If b* = (*O, K, G, t*) ∈ Dom **ℒ** *then set*

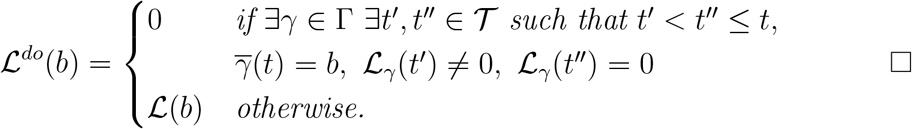

#### Proposition 6.2.2.

*By the notation of the previous definition, the following statements hold*.

1. **ℒ**^*do*^ ≤ **ℒ**,
2. **ℒ**^*do*^ *satisfies property (do)*,
3. *If* **ℒ** *satisfies property (es), then so does* **ℒ**^*do*^,
4. *If* **ℒ**_1_ ∼ **ℒ**_2_, *then* 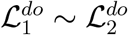.

*Proof*. We only show 4, and it is a consequence of the following. If *γ* ∈Γ, *t′, t′′* ∈ **𝒯**, then the followings are equivalent by 2.2.22.

a) 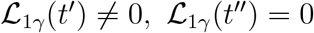

b) 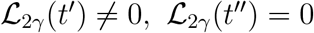. □

#### Proposition 6.2.3.

*Let a life-level-base system* Λ_*b*_ *be given together with a* Λ_*b*_*-base-function* **ℒ** *which satisfies property (ss). Suppose that*

1. *b, b′* ∈ Dom **ℒ**, *b* ⊆_*𝒪*_ *b′*, **ℒ** (*b′*) *>* 0 *implies that* **ℒ** (*b*) *>* 0 *and*
2. *b, b′* ∈ Dom **ℒ**, 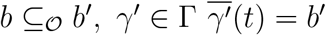 *implies that* ∃*γ* ∈ Γ *such that* 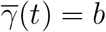 *and* 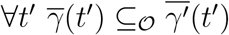.

*Then* **ℒ**^*do*^ *also satisfies property (ss)*.

*Proof*. Let *b, b′* ∈ Dom **ℒ**, *b* ⊆_*𝒪*_ *b′*.

If **ℒ**^*do*^(*b′*) *>* 0, then if **ℒ**^*do*^(*b*) = 0, then **ℒ**^*do*^(*b*) ≤ **ℒ**^*do*^(*b′*) obviously holds.

If **ℒ**^*do*^(*b*) *>* 0 then it holds by property (ss) for **ℒ**.

If **ℒ**^*do*^(*b′*) = 0, then if **ℒ** (*b′*) = 0, then **ℒ** (*b*) = 0 and **ℒ**^*do*^(*b*) = 0. If **ℒ** (*b′*) *>* 0, then ∃*γ′* ∈ Γ ∃*t′, t′′* ∈ **𝒯** such that 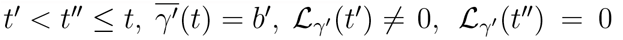. Let *γ* ∈ Γ be chosen according to the condition. Then **ℒ**_*γ*_(*t′*)≠ 0, **ℒ**_*γ*_(*t′′*) = 0. Hence **ℒ**^*do*^(*b*) = 0. □

### 6.3 Smoothening the life-level curve

Here our original aim is to create a little more sophisticated version of 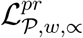. The reason for that is that we want a kind of solution for the problem mentioned in remark 4.2.3. In order to handle that we take averages in the future by a weight function. In this sense, we bring the future values back to the present in a way, that the further in the future the value is, the less we take that into account. Of course we define this method generally as a new operation.

More importantly, this operation can also be considered as a kind of general method of how one can transform a life-level function whose values are based on information in the present only to a function that takes into account the near future as well. See example 6.3.8.

#### Definition 6.3.1.

*Let a life-level-system* Λ *be given in the less general context. Let a property-system* ⟨**𝒫**, *w*⟩ *be also given together with the* ∝ *relation. Let t*_0_ ∈ **𝒯** *and* (*O, K, G, t*_0_) *be an admissible bio-bit. Set*

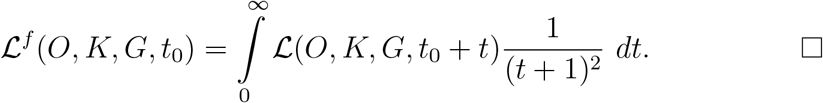

#### Proposition 6.3.2.

*Let* **ℒ**^*f*^ *be defined as in 6.3.1*. *Then* Ran **ℒ**^*f*^ ⊂[0, +∞) *and* **ℒ**^*f*^ *has properties (es), (do), (ss). Proof*. **ℒ** ≥ 0 gives that Ran **ℒ**^*f*^ ⊂ [0, +∞).

(es): If **ℒ** (∅, *K, G, t*) = 0 then so does **ℒ**^*f*^ (∅, *K, G, t*).

(do): Suppose that **ℒ**^*f*^ (*O, K, G, t*_0_) = 0. It gives that the function *t* 1→ **ℒ**^*f*^ (*O, K, G, t*) is 0 on [*t*_0_, +∞) expect on a set of measure 0. Then evidently **ℒ**^*f*^ (*O, K, G, t*_1_) = 0 for *t*_1_ *> t*_0_.

(ss): If *O* ⊂ *O′, O, O′* ∈ **𝒪**, then ∀*t′* **ℒ** (*O, K, G, t′*) ≤ **ℒ** (*O′, K, G, t′*) which yields that **ℒ**^*f*^ (*O, K, G, t′*) ≤ **ℒ**^*f*^ (*O′, K, G, t′*).

**Remark 6.3.3**. *Note that in the proof of property (do), we have not used property (do) of* **ℒ**. □

#### Corollary 6.3.4.

*Let* 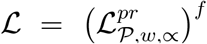 *(see definition 4.2.1)*. *Then* Ran **ℒ** ⊂ [0, 1] *and* **ℒ** *has properties (es),(do). Moreover if* ∝ *is increasing, then* **ℒ** *has property (ss) as well*.

*Proof*. Only **ℒ** ≤ 1 needs explanation and it is because 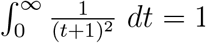. □

Unfortunately we cannot state that the ^*f*^ operation keeps the ordering relation among bio-bits, however it keeps it in a straightforward case.

#### Proposition 6.3.5.

*Let* ℒ^*f*^ *be defined as in 6.3.1*. *Let t* ∈***𝒯****and* (*O*_1_, *K*_1_, *G*_1_, *t*), (*O*_2_, *K*_2_, *G*_2_, *t*) ∈Dom ℒ*such that ∀t′*≥ *t* ℒ(*O*_1_, *K*_1_, *G*_1_, *t′*)≤ℒ (*O*_2_, *K*_2_, *G*_2_, *t′*) *holds. Then* ∀*t′*≥ *t* ℒ^*f*^ (*O*_1_, *K*_1_, *G*_1_, *t′*) ≤ ℒ^*f*^ (*O*_2_, *K*_2_, *G*_2_, *t′*) *also holds*.

**Remark 6.3.6**. *Evidently the* ^*f*^ *operation does not necessarily keep equivalence:*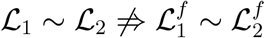.

**Remark 6.3.7**. *Certainly one may use other weight functions instead*

*of* 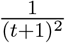 *according one’s need. E.g. one may consider to use one function from the family* 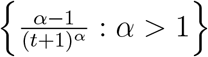. *The smaller α one takes, the more* **ℒ**^*f*^ *takes into account later values of* ℒ. *In the contrary, for big α*, ℒ^*f*^ *mainly takes into account the values close to the present*.

#### Example 6.3.8.

*Suppose that we have a life-level function whose values are based on information in the present only. Now if an organism lives close to a just erupting volcano, then if we apply the current operation to the original life-level function, the eruption itself can decrease the present life-level of the organism*.

### 6.4 Moving averages

One can examine the usual moving average of a life-level function. However we need some small restriction for that.

#### Definition 6.4.1.

*Let a life-level-system* Λ *be given. We call* Γ ***unique at each point*** *if* ∀ (*O, K, G, t*) ∈ Dom ℒ *there is a unique γ* ∈ Γ *such that* 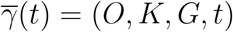.

#### Definition 6.4.2.

*Let a life-level-system* Λ *be given such that* Γ *is unique at each point. Let l* ∈ ℝ^+^. *Let* (*O, K, G, t*_0_) ∈ Dom **ℒ** *γ* ∈ Γ *such that* 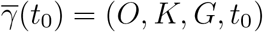. *Then the* ***l-length moving average through γ*** *is*

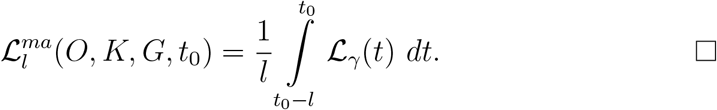

#### Proposition 6.4.3.

*Let a life-level-system* Λ *be given such that* Γ *is unique at each point. Let l* ∈ ℝ^+^. *Then* Ran 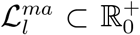 *and* 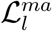 *satisfies property (do)*.

*Proof*. **ℒ**_*γ*_ (*t*) ≥ 0 which gives that Ran 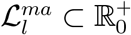.

If 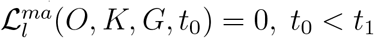, then ∃*t′* ∈ [*t*_0_ −*l, t*_0_] such that **ℒ**_*γ*_ (*t′*) = 0. Which yields that ∀*t′′ > t′* **ℒ**_*γ*_ (*t′′*) = 0 by property (do) for **ℒ**. Which implies

that

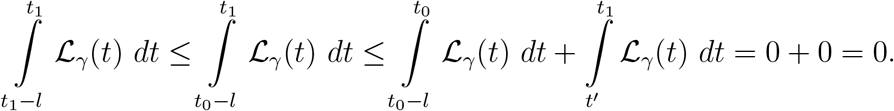

Therefore 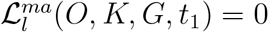. □

#### Proposition 6.4.4.

*Let a life-level-system* Λ *be given such that* Γ *is unique at each point. Let l* ∈ ℝ^+^. *Assume that O, O*^*′*^ ∈ **𝒪**, *O* ⊂ *O*^*′*^, (*O, K, G, t*_0_) ∈ Dom ℒ *and* 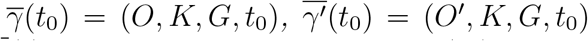*implies that* 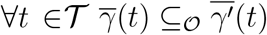. *Then* 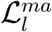 *satisfies property (ss)*.

*Proof*. Trivial consequence of property (ss) of ℒ.

**Remark 6.4.5**. *Property (es) usually does not hold for* 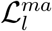, *especially after the right end point of the interval* {*t* ∈ **𝒯** : γ_*O*_(*t*) ≠ ∅}. *If one wants property (es) to be held, then the definition of* 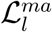 *has to be modified by adding that condition explicitly*. □

This operation does not necessarily keep equivalence, however the following holds.

#### Proposition 6.4.6.

*Let a life-level-system* Λ *be given such that* Γ *is unique at each point and let* ***ℒ***_1_, *ℒ*_2_ *be two associated life-level functions. Let l* ∈ ℝ^+^. *If ℒ*_1_ ≤ **ℒ**_2_, *then* 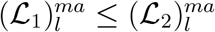. □

Now we are going to omit the condition that Γ is unique. Instead, we use probability on Γ_*b*_ (see 2.2.31). We also change to “future” window from “past” window.

#### Definition 6.4.7.

*Let a countable life-level-system* Λ *be given. Let l* ∈ ℝ^+^. *Let b* = (*O, K, G, t*_0_) ∈ Dom ℒ *and* P *be a probability function given on* Γ_*b*_. *Then set*

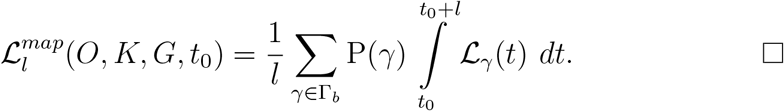

Similarly to 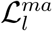 one can show the following.

#### Proposition 6.4.8.

*By the notations and assumptions of 6.4.7*, Ran 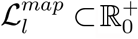 *and* 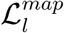 *satisfies property (do)*.

### 6.5 Discretizing

In some cases, one may want discrete values on one’s life-level system instead of some “continuous” ones. We will investigate two such cases: on life-level values and time: One may want discrete life-level values, e.g. integers, or one may prefer discrete time points, e.g. measurement at every day.

#### Proposition 6.5.1.

*Let a life-level-system* Λ *be given. Let* 0 *< ε* ≤ 1. *Then* 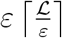 *is an associated life-level function that is alive-equivalent with ℒ*.

*Proof*. Obviously the function 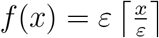 is increasing and *f* (*x*) = 0 ⇔*x* = 0. Then apply 6.1.6.

**Remark 6.5.2**. *If instead, one tries* ⌊**ℒ**⌋ *or just the usual rounding, then one may loose alive-equivalence, however it may result an associated life-level function*.

There may be a given small threshold at 0, under which an gorganism cannot be considered alive, and one may want to force it in one’s modified model in the following way.

#### Proposition 6.5.3.

*Let a life-level-system* Λ *be given, with a threshold* 0 *< ε <* 1 *such that γ* ∈ Γ, 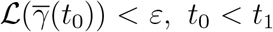 *implies that* 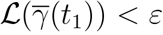. *Then* 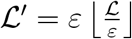 *is an associated life-level function such that* **ℒ**^*′*^(*b*) = 0 ⇔ **ℒ**(*b*) *< ε*.

*Proof*. (es) and (ss) can be similarly verified as in the proof of 6.1.1 for function 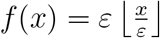.

(do): If 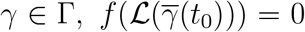, *t*_0_ *< t*_1_, then 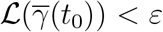which yields that 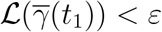which gives that 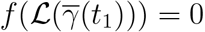.

**Remark 6.5.4**. *This method (6.5.3) can also be applied when one wants to suppress small fluctuations in the values*. □

Now we describe a discretizing method by averages on time. Roughly, we take the average of the life-level around every integer.

#### Proposition 6.5.5.

*Let a life-level-system* Λ = ⟨ **𝒪**, ⊂⟩, ⟨ κ, ⊂⟩, **𝒢, 𝒯**, Θ, *ℒ be given in the less general context for* 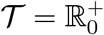. *For z* ∈ ℕ*set*

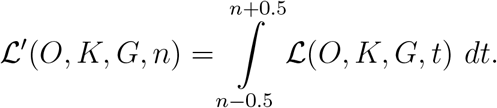

*Then* Λ^*′*^ = ⟨⟨ **𝒪**, ⊂⟩, ⟨ κ, ⊂⟩, **𝒢, 𝒯** *′*, Θ, *ℒ*^*′*^ *is a life-level-system where* ***𝒯*** ^*′*^ = ℕ_0_ □

*Proof*. Property (es) is trivial.

If **ℒ**^*′*^(*O, K, G, n*) = 0, then **ℒ** (*O, K, G, n*) = 0, hence by (do) on **ℒ**, we get that ∀*t > n* **ℒ** (*O, K, G, t*) = 0, therefore ∀*m > n* **ℒ**^*′*^(*O, K, G, m*) = 0. □

If *O*_1_ ⊂ *O*_2_, then 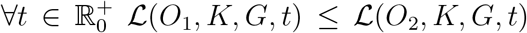, thus ∀*n* ∈ ℕ_0_ **ℒ**^*′*^(*O*_1_, *K, G, n*) ≤ **ℒ**^*′*^(*O*_2_, *K, G, n*).

### 6.6 Distance between life-level functions

Now we want to compare how close or how far the values of two life-level functions are from each other. We define three such distances.

#### Definition 6.6.1.

*Let a life-level-base-system* Λ *be given together with two associated life-level functions* **ℒ**_1_, **ℒ**_2_ *such that* Dom **ℒ**_1_ = Dom **ℒ** _2_. *Set*

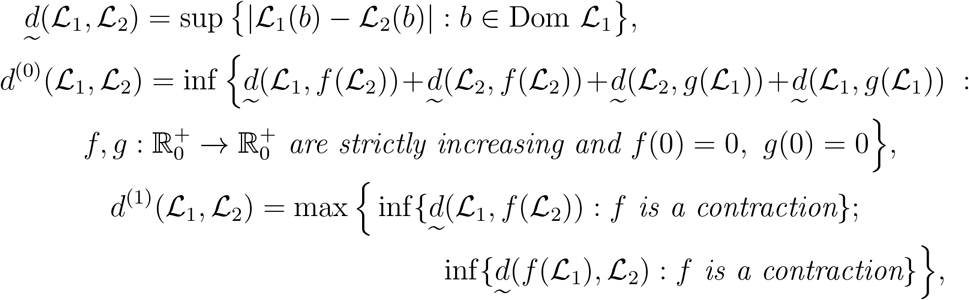

*where* 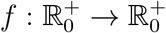 *is called a contraction if f is strictly increasing, f* (0) = 0 *and* 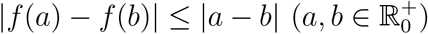. ◻

#### Proposition 6.6.2.

*Let a life-level-base-system* Λ *be given together with associated life-level functions on common domain. Then* 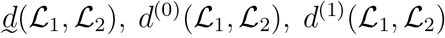 *are pseudo-metrics on the set of associated life-level functions*.

*Proof*. It can be proved in the usual way that 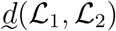 is a pseudo-metric.

Now we prove it for *d*^(0)^ (**ℒ**_1_, **ℒ**_2_) and *d*^(1)^ (**ℒ**_1_, **ℒ**_2_).

Clearly *d*^(0)^(**ℒ**_1_,**ℒ**_1_) = *d*^(1)^(**ℒ**_1_, **ℒ**_1_) = 0, since take *f* (*x*) = *g*(*x*) = *x* in both cases.

Obviously *d*^(0)^(**ℒ**_1_, **ℒ**_2_) = *d*^(0)^(**ℒ**_2_,**ℒ**_1_) and *d*^(1)^(**ℒ**_1_, **ℒ**_2_) = *d*^(1)^(**ℒ**_2_, **ℒ**_1_) both hold because both defining formulas are symmetric (for *d*^(0)^ just swap *f* and *g*).

To see the triangle inequality for *d*^(0)^, let *ε >* 0 and take corresponding functions *f, g, h, j* such that

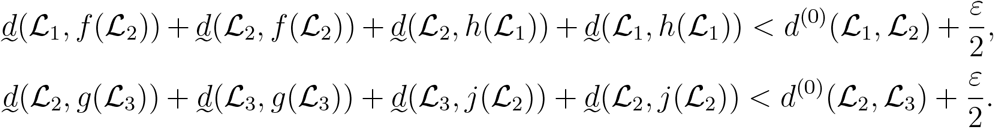

By

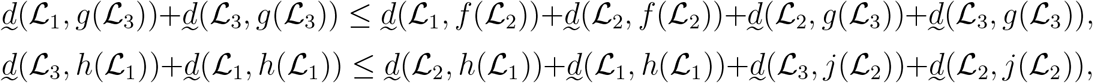

we get that

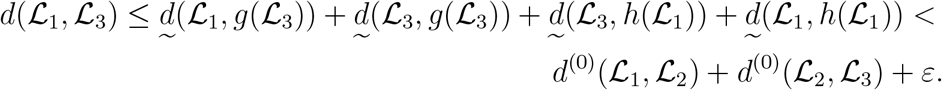

When *ε* tends to 0, then we get the claim.

To see the triangle inequality for *d*^(1)^, let *ε >* 0 and take corresponding functions *f, g* such that

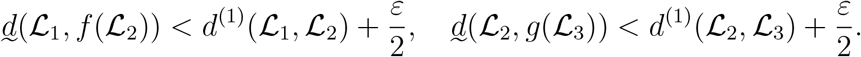

But *f* being a contraction gives that

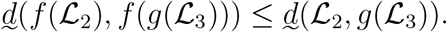

By the triangle inequality for 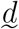 we get that

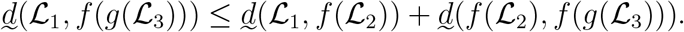

If *f, g* are contractions, then so is *f* о *g*, hence we get that

inf{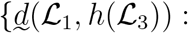 is a contraction} 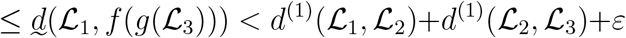.

It holds for all *ε >* 0, therefore

inf{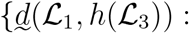 is a contraction} ≤ *d*^(1)^(**ℒ**_1_, **ℒ**_2_) + *d*^(1)^(**ℒ**_2_, **ℒ**_3_).

Now apply the same argument for 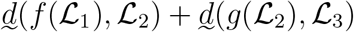 and similarly we end up with

inf{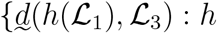 is a contraction} ≤ *d*^(1)^(**ℒ**_1_, **ℒ**_2_) + *d*^(1)^(**ℒ**_2_, **ℒ**_3_),

which altogether yield that *d*^(1)^(**ℒ** _1_, **ℒ** _3_) ≤ *d*^(1)^(**ℒ**_1_,**ℒ**_2_) + *d*^(1)^(**ℒ**_2_, **ℒ**_3_). ◻

**Remark 6.6.3**. *Unfortunately neither d*^(0)^ *nor d*^(1)^ *is not good enough to force d*(**ℒ**_1_, **ℒ**_2_) = 0 *for equivalent life-level functions* **ℒ**_1_, **ℒ**_2_.

### 6.7 Convergence of life-level functions

Suppose that a new species identified, but first, only one representative is found, moreover it is hard to categorize it taxonomically at first sight. Furthermore its life cycle is not fully clear, nor its relation to its surroundings. In such a case, we can assign a rough estimate value to it as a life-level. As information grows, we can refine our estimations gradually, and get a better and better values for its life-level. Hence in this case we have constructed a sequence of life-level functions which is hopefully converges to some fixed value.

#### Definition 6.7.1.

*Let a life-level-base-system* Λ *be given together with a sequence of associated life-level functions* (**ℒ**_*n*_) *on the same domain. We say that* (**ℒ**_*n*_) ***converges uniformly*** *to* **ℒ**, *if d*(**ℒ**_*n*_, **ℒ**) → 0. *In this case, we will use the notation:* **ℒ** _*n*_ → **ℒ**.

#### Definition 6.7.2.

*Let a life-level-base-system* Λ *be given together with a sequence of associated life-level functions* (**ℒ**_*n*_) *on the same domain. We say that* (**ℒ**_*n*_) ***converges pointwise*** *to* **ℒ**, *if* ∀*b* ∈ Dom **ℒ ℒ**_*n*_(*b*) → **ℒ**(*b*) *holds*.

#### Proposition 6.7.3.

*Let a life-level-base-system* Λ *be given together with a sequence of associated life-level functions* (**ℒ**_*n*_) *on the same domain. If* (**ℒ**_*n*_) *converges to* **ℒ** *uniformly, then it converges pointwise to* **ℒ**. *If* Λ *is finite*, (**ℒ**_*n*_) *converges to* **ℒ** *pointwise, then it converges uniformly*. ◻

#### Proposition 6.7.4.

*Let a life-level-base-system* Λ *be given together with a sequence of associated life-level functions* (**ℒ**_*n*_) *on the same domain, and let* (**ℒ**_*n*_) *converge to* **ℒ** *pointwise. Then* **ℒ** *satisfies properties (es) and (ss)*.

*If there is an ε >* 0 *such that n* ∈ ℕ, *b* ∈ Dom **ℒ**_*n*_, **ℒ**_*n*_(*b*) *< ε implies that* **ℒ**_*n*_(*b*) = 0, *then (do) holds as well*. ◻

**Remark 6.7.5**. *Clearly, alone even uniform convergence does not guarantee property (do)*.

#### Proposition 6.7.6.

*Let a life-level-base-system* Λ *be given together with a sequence of associated life-level functions* (**ℒ**_*n*_) *on the same domain, and let* (**ℒ**_*n*_) *converge to* **ℒ** *pointwise. Then the following holds*.

1. *If* ∀*n* **ℒ**_*n*_ *is subaddtive (additive), then so is* **ℒ**.
2. *If the convergence is uniform and* ∀*n* **ℒ**_*n*_ *is continuous, then so is* **ℒ**. ◻

## 7 Extension and fine tuning of life-level systems

In this section we want to provide some general extension methods for already existing life-level-systems. In all cases, the need for extension will be some biological necessity, a kind of natural requirement for getting a greater, broader system. Usually we will provide many solutions, not only one.

The second group of modifications are about overriding some of the original, already given life-level values in order to get better models. Generally speaking, in those cases, we want to take into account the life-levels of some biologically “close” other gorganisms.

In some cases, the extension and overriding may happen at the same time.

### 7.1 Life-level of a group-type gorganisms

Let us expose the not too precise title: Suppose that we have a life-level function defined on bio-bits that are based on non group-type gorganisms only. Our aim is to extend this function somehow to bio-bits that based on group-type gorganism too. Actually, all methods here will be more general: we will not only apply them for group-type gorganisms, but also any gorganisms with smaller components.

Operations described here can also be considered as technical ones for adding the missing property (ss) to a Λ-base-function.

One more comment: it would be tempting to think that we are just going to assign life-level values to species, but we are still far from that currently, and our aim is different. (See subsection 8.3 and section 9.)

We present three such operations.

#### Definition 7.1.1.

*Let a life-level-system* Λ *be given. Let* (*O, K, G, t*) *be an admissible bio-bit. Then set*

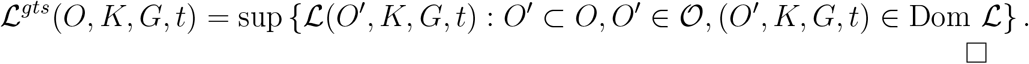

#### Proposition 7.1.2.

*Let a life-level-system* Λ *be given. If* (*O, K, G, t*) ∈ Dom **ℒ** *then* **ℒ**(*O, K, G, t*) = **ℒ**^*gts*^(*O, K, G, t*) *i.e*. **ℒ**^*gts*^ *is an extension of* **ℒ**.

*Proof*. Property (ss) verifies the claim. ◻

#### Proposition 7.1.3.

*Let a life-level-system* Λ *be given. Then* ℒ^*gts*^ *satisfies properties (es) and (ss)*.

*Proof*. (es): Clearly *O* = ∅ has ∅ as an only subset.

(ss): If *O*_1_ ⊂ *O*_2_ then *O*^*′*^ ⊂ *O*_1_ implies that *O*^*′*^ ⊂ *O*_2_ which gives immediately that **ℒ**^*gts*^(*O*_1_, *K, G, t*) ≤ **ℒ**^*gts*^(*O*_2_, *K, G, t*). □

**Remark 7.1.4**. *When showing property (ss) for* **ℒ**^*gts*^, *we have not used property (ss) for* **ℒ**. *I.e. this operation “adds” property (ss) to a* Λ*-base-function*.

If new gorganisms do not migrate into the group, then property (do) holds as well.

#### Proposition 7.1.5.

*Let a life-level-system* Λ *be given such that* Λ *is* Γ*-below-synchronized-backward. Then* **ℒ**^*gts*^ *satisfies properties (do)*.

*Proof*. Let *γ* ∈ Γ, *γ***𝒪** (*t*_0_) ≠∅, **ℒ**^*gts*^ *γ*_*𝒪*_(*t*_0_), *γ*_*𝒦*_(*t*_0_), *γ*_*𝒢*_(*t*_0_), *t*_0_ = 0, and *t*_0_ *< t*_1_. If *O*^*′*^ ⊂ *γ*_*𝒪*_(*t*_1_), then 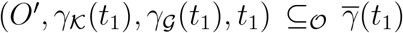, and choose *γ*^*′*^ ∈ Γ according to the Γ-below-synchronized-backward condition: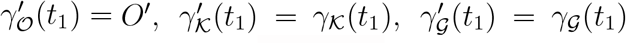 and 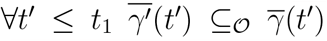. But then by (ss) for **ℒ**, 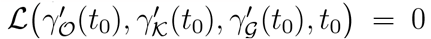 holds. Then by (do) for **ℒ**, we get that 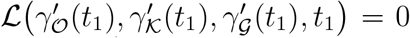 holds. Which gives that **ℒ**^*gts*^ ((*γ*_*𝒪*_(*t*_1_), *γ*_𝒦_(*t*_1_), *γ*_𝒢_(*t*_1_), *t*_1_)= 0. ◻

#### Proposition 7.1.6.

*Let a life-level-system* Λ *be given. If* **ℒ**_1_ ≤ **ℒ** _2_, *then* 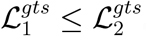.

#### Proposition 7.1.7.

*Let a finite life-level-system* Λ *be given. Then* **ℒ**_1_ ∼ **ℒ**_2_ *implies that* 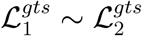.

*Proof*. By 2.2.19 we have to show that 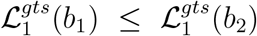 implies that 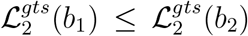 (*b*_1_, *b*_2_ ∈ Dom **ℒ**_1_). Then by finiteness of **𝒪** there are 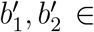 Dom **ℒ**_1_ such that 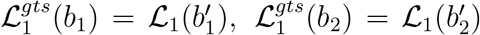 and 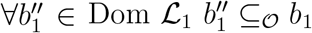 implies that 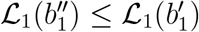, and 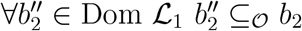 implies that 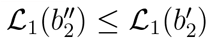 hold. By **ℒ** _1_ ∼ **ℒ** _2_ we get that 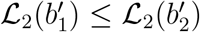 and 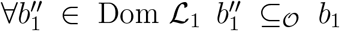 [implies that 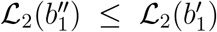, and 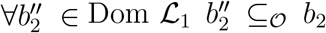 implies that 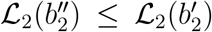. These clearly give that 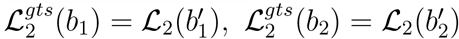 and then 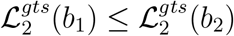 holds. ◻

#### Definition 7.1.8.

*Let a finite life-level-system* Λ *be given such that* ⟨𝒪, ⊂⟩, *is a lattice (let* ∪ *denote the supremum, while* ∩ *the infimum of two elements). Let*

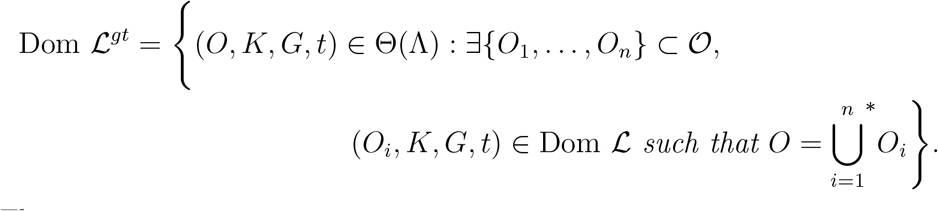

*Then set*

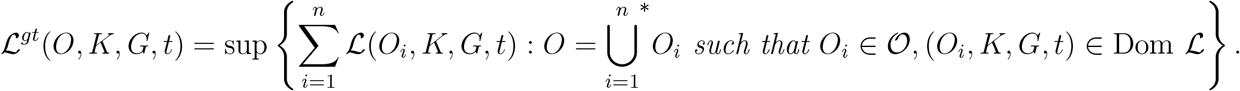

□
**Remark 7.1.9**. *The* sup *regards for a set because a group-type gorganism can have many partitioning and we wanted to take all into account*.

#### Proposition 7.1.10.

1. Dom **ℒ** ⊂ Dom **ℒ**^*gt*^,
2. *If* (*O, K, G, t*) ∈ Dom **ℒ** *then* **ℒ**(*O, K, G, t*) ≤ **ℒ**^*gt*^(*O, K, G, t*).

*Proof*. Both statements follow from the fact that for *O*, {*O*} is a one element admissible partition. ◻

**Remark 7.1.11**. *One can apply that method to extend a life-level function with an original domain containing bio-bits* (*O, K, G, t*) *such that O is non group-type*.

**Remark 7.1.12**. *With some care, the definition 7.1.8 could be formulated for countable life-level-system with bounded life-level function as well*.

#### Proposition 7.1.13.

*Let a finite life-level-system* Λ *be given. Then* **ℒ**^*gt*^ *satisfies property (es). If* Λ *is* Γ*-below-synchronized-backward, then* **ℒ**^*gt*^ *satisfies property (do) as well*.

*Proof*. (es): Clearly *O* = ∅ has an only partition.

(do): Let *γ* ∈ Γ, *γ*_*𝒪*_(*t*_0_)≠∅, **ℒ**^*gt 0*^*(γ*_***𝒪***_ (*t*_0_), *γ*_***𝒦***_ (*t*_0_), *γ*_***𝒢***_ (*t*_0_), *t*_0_)= 0, and *t*_0_ *< t*_1_. If *O*^*′*^⊂ *γ*_***𝒪***_ (*t*_1_), then choose *γ*^*′*^ ∈ Γ according to the Γ-below-synchronized-backward condition: 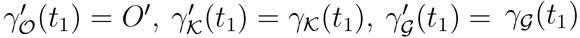 and 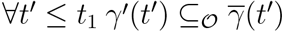. But then by (ss) for **ℒ**, 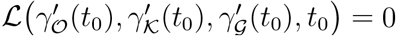 holds. Then by (do) for **ℒ**,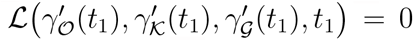 holds. Which gives that **ℒ**^*gt*^ *γ*_*𝒪*_(*t*_1_), *γ*_***𝒦***_ (*t*_1_) *γ****𝒢***(*t*_1_), *t*_1_ = 0. ◻

#### Proposition 7.1.14.

*Let us use the notations of definition 7.1.8*. *Suppose that* **𝒪** *is closed for subtraction, i.e. O*_1_, *O*_2_ ∈ **𝒪** *implies that O*_2_ − *O*_1_ ∈ 𝒪. *Then* **ℒ**^*gt*^ *satisfies properties (ss)*.

*Proof*. If *O* ⊂ *O*^*′*^ and 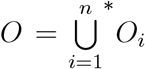 such that *O*_*i*_ ∈ **𝒪**, (*O*_*i*_, *K, G, t*) ∈ Dom **ℒ** then 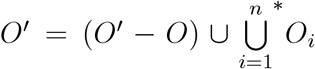 is an admissible partition of *O*^*′*^. Obviously 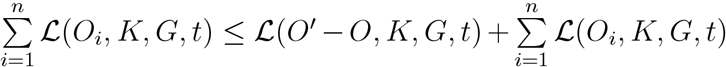. Hence we get that **ℒ**^*gt*^(*O, K, G, t*) ≤ **ℒ**^*gt*^(*O*^*′*^, *K, G, t*).

**Remark 7.1.15**. *We just sketch a third such operation that is a slight modification of* **ℒ**^*gt*^.

*Modify the definition 7.1.8 in a way that replace* 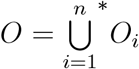*with* 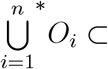 *everywhere. Let us denote the new function by* **ℒ**^*gtp*^. *One can show similarly that it satisfies properties (es) and (do), and (ss) as well, and here we do not need any additional condition for that*. □

### 7.2 Life-level determined by the supporting-group

This subsection is about the possible options that it may be worth considering that the life-level of a gorganism can also be determined by its supporting-group. More precisely if a life-level function is given, then an operation can be defined that creates a new life-level-base function from it using original values on the elements of the supporting-group (it does not necessarily satisfy all properties (es),(do) and (ss)).

In some sense this kind of operations are the opposite of the ones described in the previous subsection. Here we pull the life-levels back, where there we pushed them up.

We will provide two such operations.

Let us outline the underlying motivation of the first one by the following example.

#### Example 7.2.1.

*Consider a not fertile (young) man with a supporting-group of fertile men and women. Suppose that the relationship between the man and the group is extremely strong in the sense that in the short run the survival of the whole group depends on the man, without the man the whole group would extinct in a short time. The man is incapable to inherit any of its properties, when he dies none of his properties survives on any of his development-paths. However almost all of his properties exist in at least one member of the group. Hence those properties will survive with great certainty however in a different way*.

*In this example, the man had a crucial role in the survival of the group but this also suggests that it is worth considering the survival of the properties of the gorganism by its fellow members in its supporting-group*.

First we can stay on a more general level than we outlined in the previous example, namely we do not need to refer to properties, we can just say that the life-level of our gorganism should be at least as high as any of its fellow members’ in its supporting-group.

#### Definition 7.2.2.

*A life-level-system* Λ *is called* ***group dissolvable*** *if* (*O, K, G, t*) ∈ Dom **ℒ**, *O*^*′*^ ∈ *G* ∩ **𝒪** *implies that* (*O*^*′*^, *K, G, t*) ∈ Dom **ℒ**.

#### Definition 7.2.3.

*Let a group dissolvable life-level-system* Λ *be given. Let* (*O, K, G, t*) ∈ Dom **ℒ**. *Then set*

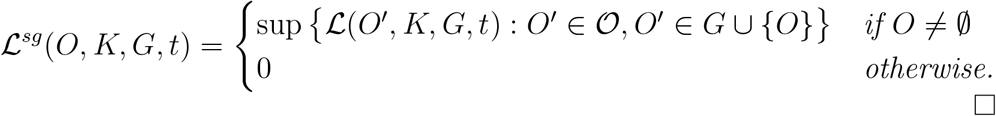

**Remark 7.2.4**. *The term “*∪{*O*}*” was needed because it may happen that O* ∉*G and without that* **ℒ**^*sg*^ (*O*) *would be (partially) independent from* **ℒ** (*O*).

#### Proposition 7.2.5.

*Let a group dissolvable life-level-system* Λ *be given. Let* (*O, K, G, t*) ∈ Dom **ℒ**. *Then the following statements hold*.

1. **ℒ** (*O, K, G, t*) ≤ **ℒ***sg* (*O, K, G, t*)
2. *If O, O*^*′*^ ∈ *G* ∩ **𝒪** *then* **ℒ**^*sg*^ (*O, K, G, t*) = **ℒ**^*sg*^ (*O*^*′*^, *K, G, t*).

#### Proposition 7.2.6.

*Let a group dissolvable life-level-system* Λ *be given. Then the operation* ^*sg*^ *is idempotent i.e*. **ℒ**^*sg*^ = (**ℒ**^*sg*^)^*sg*^. *Proof*. Clearly

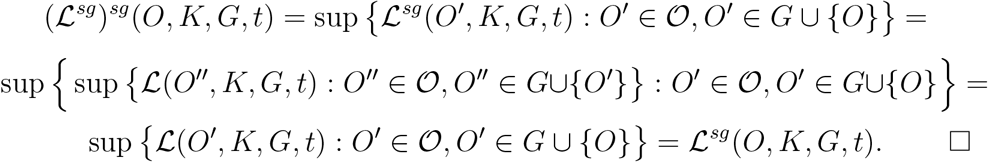

#### Proposition 7.2.7.

*Let a group dissolvable life-level-system* Λ *be given. Then* Ran 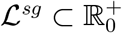 *and* **ℒ**^*sg*^ *satisfies properties (es) and (ss)*.

*Proof*. (ss) is a consequence of (ss) for **ℒ**^*sg*^. The other statements are (also) trivial. □

#### Proposition 7.2.8.

*Let a group dissolvable life-level-system* Λ *be given*.

*Suppose that any of the following holds for* **ℒ** = **ℒ**^*sg*^.

1. *If* **ℒ**(*O, K, G, t*) = 0 *then O* = ∅.
2. 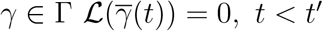, *t < t*^*′*^ *implies that γ*_*𝒪*_(*t*^*′*^) = ∅ (Λ *is* Γ*-vanishing). In this case* **ℒ**^*sg*^ *satisfies properties (do)*. □

**Remark 7.2.9**. *One may recognize similarities between* **ℒ**^*gts*^ *and* **ℒ**^*sg*^, *however using sub-gorganisms of O or sub-gorganisms of the supporting-group G makes an essential difference*.

Roughly speaking we are now going to define how long after *t*_0_ property *p* survives in the group *G*.

#### Definition 7.2.10.

*Let a propertized life-level-base-system* ⟨Λ_*b*_, **𝒫**, *w*, ∝⟩ *be given. Let p* ∈ **𝒫** *and t* ∈ **𝒯**. *Let G* ∈ **𝒢**(*t*), *K* ∈ **𝒦**(*t*). *Set*

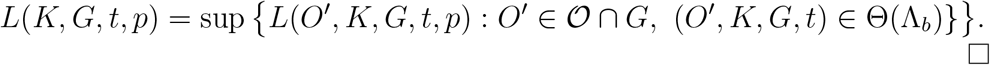

#### Proposition 7.2.11.

*Let a propertized life-level-base-system* ⟨Λ_*b*_, **𝒫**, *w*,∝⟩ *be given. Then*

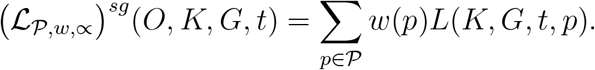

*Proof*. By the definition of **ℒ**_*P,w,∝*_, it can be readily seen that

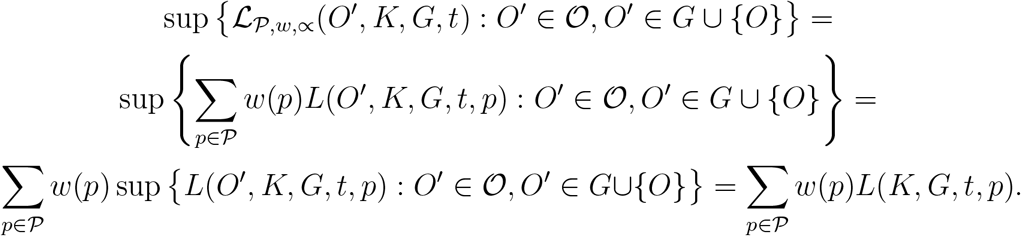

□

#### Example 7.2.12.

*Let us apply this method for one of two identical twins with his brother being alive or not. In the first case, we put his brother into his supporting-group. Then we clearly end up with a higher value when the brother is alive than not*.

#### 7.2.1 Using a weight

We are going to provide a more sophisticated model comparing to the previous one (7.2). Here we assume that a weight function is also given on the elements of the supporting-group that is supposed to express how much an element affects the life-level of the bio-bit.

##### Definition 7.2.13.

*Let a group dissolvable life-level-system* Λ *be given. Let* (*O, K, G, t*) ∈ Dom **ℒ**. *A weight function on the supporting-group of* (*O, K, G, t*) *is a function w such that*

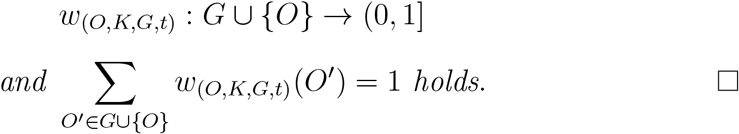

##### Definition 7.2.14.

*Let a group dissolvable life-level-system* Λ *be given together with weight functions on the supporting-groups. Let* (*O, K, G, t*) ∈ Dom **ℒ**. *Then set*

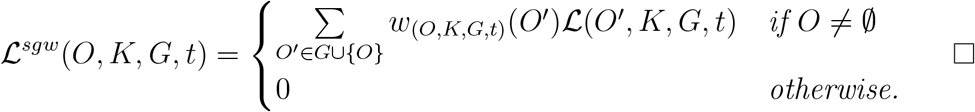

##### Definition 7.2.15.

*Let a group dissolvable life-level-system* Λ *be given together with weight functions on the supporting-groups. If* **ℒ** = **ℒ**^*sgw*^ *holds, then we call* **ℒ *sg-balanced*** *regarding the given weight functions*.

##### Proposition 7.2.16.

*Let a group dissolvable life-level-system* Λ *be given together with weight functions on the supporting-groups. If* **ℒ** *sg-balanced, then*

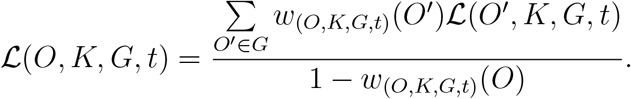

*Proof*. This simply follows from

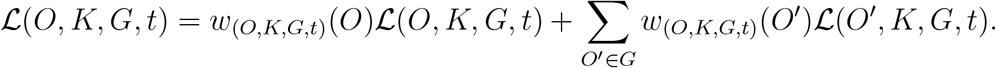

□

**Remark 7.2.17**. *This notion “sg-balanced” can be simply used to calculate how emigration from/immigration to the supporting-group affects the life-level of the underlying gorganism*.

##### Proposition 7.2.18.

*Let* **ℒ**^*sgw*^ *be defined as in 7.2.14*. *Then* Ran **ℒ**^*sgw*^ ⊂ [0, +∞) *and* **ℒ**^*sgw*^ *satisfies property (es)*.

##### Proposition 7.2.19.

*Let a group dissolvable life-level-system* <u>Λ</u> *be given in the less general context together with weight functions on the supporting-groups. Then* **ℒ**^*sgw*^ *satisfies property (do)*.

*Proof*. Let **ℒ**^*sgw*^ (*O, K, G, t*) = 0 and *t < t*^*′*^. Then ∀*O*^*′*^ ∈ *G*∪{*O*} **ℒ**(*O*^*′*^, *K, G, t*) = 0 which gives that ∀*O*^*′*^ ∈ *G* ∪ {*O*} **ℒ**(*O*^*′*^, *K, G, t*^*′*^) = 0 by property (do) for **ℒ**. This obviously yields that **ℒ**^*sgw*^ (*O, K, G, t*^*′*^) = 0.

Now we recursively define the compositions of this operation.

##### Definition 7.2.20.

*Let a group dissolvable finite life-level-system* Λ *be given together with weight functions on the supporting-groups. Let*

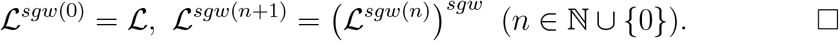

##### Proposition 7.2.21.

*Let a group dissolvable finite life-level-system* Λ *be given together with weight functions on the supporting-groups. Suppose that for all* (*O, K, G, t*) ∈ Dom **ℒ, ℒ**^*sgw*(*n*)^(*O, K, G, t*) *converges and let us denote the limit by* **ℒ**^*sgw∞*^(*O, K, G, t*). *Then* **ℒ**^*sgw∞*^ *is sg-balanced*.

*Proof*. Let (*O, K, G, t*) ∈ Dom **ℒ** and *ε >* 0. As Λ is finite, then *G* is finite hence there is *N* ∈ N such that *O*^*′*^ ∈ *G* ∪ {*O*}, *n* ≥ *N* implies that

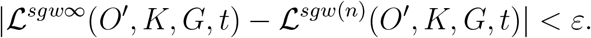

Then

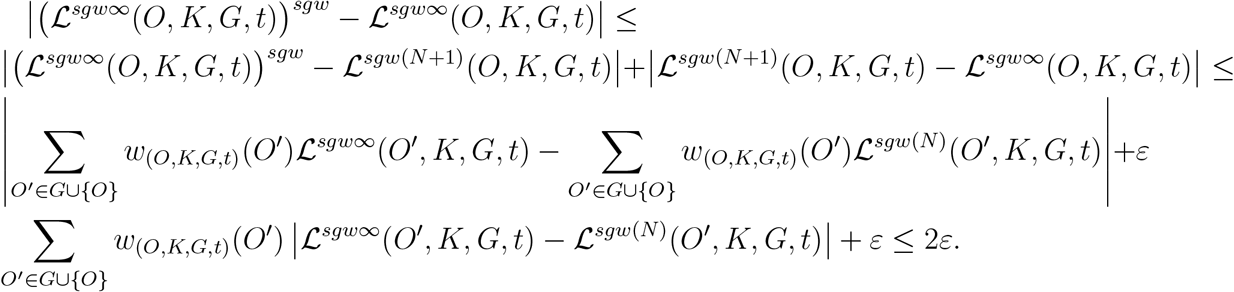

As it holds for every *ε >* 0, **ℒ**^*sgw∞*^ is sg-balanced. □

#### 7.2.2 Using survival time of properties

Based on the result of proposition 7.2.11, we provide a partially modified version of the definition 7.2.3 combined with the survival time of properties. In this case there is no initial life-level function, i.e. it is not an operation in the original sense, rather it is a new enriched type life-level derivation (cf. subsection 4.4). Roughly, here we just take into account the survival time of properties only that the gorganism owns.

Again let us start with an example which shows the basic motivation.

##### Example 7.2.22.

*Let us modify example 7.2.1 slightly*.

*The man is still not fertile but his supporting-group consists of many healthy young dogs. They live far from any other groups of people or dogs. For the survival of the dogs the existence of the man is crucial in the short run. The difference here is that there are many properties of the man that do not exist in any of the dogs. However many exist. We can examine the survival time of the common properties and use it as a value of a generalized life-level function*.

##### Definition 7.2.23.

*Let a propertized life-level-base-system* ⟨Λ_*b*_, **𝒫**,∝⟩ *be given. Let b* = (*O, K, G, t*) *be an admissible bio-bit. Set*

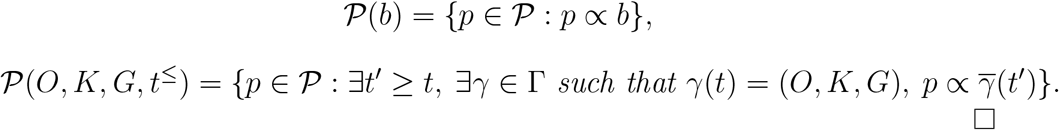

**Remark 7.2.24**. *Let a life-level-system* Λ *and a property set* **𝒫** *be given together with the* ∝ *relation. The property that* **ℒ** *is normal regarding* **𝒫** *is equivalent with* **ℒ**(*b*) *>* 0 *implies that* **𝒫**(*b*)≠ ∅.

##### Definition 7.2.25.

*Let a propertized life-level-system* ⟨Λ, **𝒫**, *w*, ∝⟩ *be given. Let* (*O, K, G, t*) *be an admissible bio-bit and O* ∈ *G*.

*Set* **ℒ**^*p*^(*O, K, G, t*) = 0 *if* **𝒫**(*O, K, G, t*) = ∅, *otherwise set*

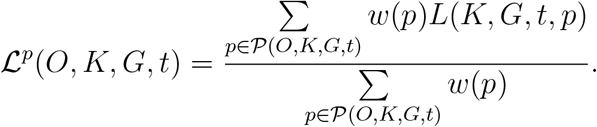

*Set*

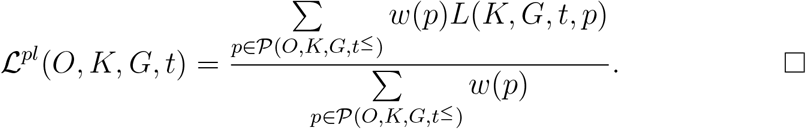

##### Proposition 7.2.26.

*With the assumptions and notations of 7.2.25*, **ℒ**^*p*^(*O, K, G, t*) *and* **ℒ**^*pl*^(*O, K, G, t*) *are the conditional expected values of the function X* : **𝒫** → ℝ∪{0}, *X*(*p*) = *L*(*K, G, t, p*) *regarding the set* **𝒫**(*O, K, G, t*) *and* **𝒫**(*O, K, G, t*^≤^) *respectively*.

*Proof*. Obviously *w* can be considered as a probability distribution on **𝒫**. □

##### Proposition 7.2.27.

*With the assumptions and notations of 7.2.25*, Ran 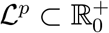 *and* **ℒ**^*p*^ *satisfies property (es). Similar statements are true for* **ℒ**^*pl*^.

##### Proposition 7.2.28.

*Let O, O*^*′*^ ∈ **𝒪**, *O* ⊂ *O*^*′*^ *and* **𝒫**(*O, K, G, t*) = **𝒫**(*O*^*′*^, *K, G, t*). *Then for this O, O*^*′*^, **ℒ**^*p*^ *satisfies property (ss)*. □

##### Example 7.2.29.

*Obviously* **ℒ**^*p*^, **ℒ**^*pl*^ *do not necessarily satisfy property (do), e.g. consider example 7.2.22*. □

##### Lemma 7.2.30.

*With the assumptions and notations of 7.2.25, let O, O*^*′*^ ∈ **𝒪**, *O* ⊂ *O*^*′*^ *and* **𝒫**(*O*^*′*^, *K, G, t*) − **𝒫**(*O, K, G, t*) = {*p*^*′*^}. *Then* **ℒ**^*p*^(*O*^*′*^, *K, G, t*) *<* **ℒ**^*p*^(*O, K, G, t*) ⇔ *L*(*K, G, t, p*^*′*^) *<* **ℒ**^*p*^(*O, K, G, t*).

*Similar statement is true for* **ℒ**^*pl*^.

*Proof*. Clearly

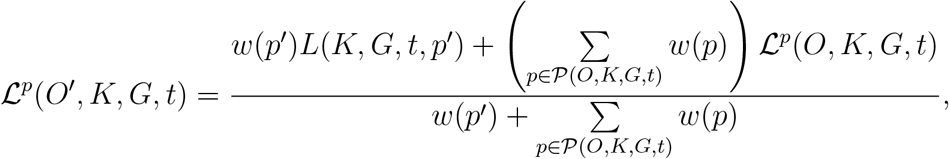

that is the weighted average of *L*(*K, G, t, p*^*′*^) and **ℒ**^*p*^(*O, K, G, t*), proving the claim.

##### Example 7.2.31.

**ℒ**^*p*^, **ℒ**^*pl*^ *do not satisfy property (ss) in general. Let O, O*^*′*^ ∈ **𝒪**, *O* ⊂ *O*^*′*^. *Let O*^*′*^ *own an only property p*^*′*^ *that O does not. If the survival time of p*^*′*^ *on the group G is smaller than* **ℒ**^*p*^(*O, K, G, t*) *then* **ℒ**^*p*^(*O*^*′*^, *K, G, t*) *<* **ℒ**^*p*^(*O, K, G, t*) *holds by 7.2.30*.

### 7.3 Life-level determined by the reverse-supporting-group

If we examine examples 7.2.1 and 7.2.22 more carefully, then it gets clear that we used a special subset of the supporting-group of the man. Moreover the support works the other way around too, or it is even “stronger” in the opposite way: the man is in the supporting-group of those other organisms. Therefore we can refine our model into that direction: the life-level of a gorganism can be determined by its reverse-supporting-group, that roughly consists of organisms that the given organism supports.

#### Definition 7.3.1.

*Let a life-level-system* Λ *be given and let b* = (*O, K, G, t*) ∈ Dom **ℒ**. *The* ***reverse-supporting-group*** *of* (*O, K, G, t*) *is*

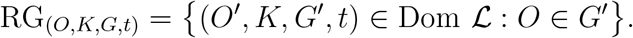

*Also set*

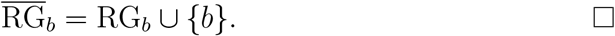

#### Definition 7.3.2.

*Let a life-level-system* Λ *be given. Let* (*O, K, G, t*) ∈ Dom **ℒ**. *Then set*

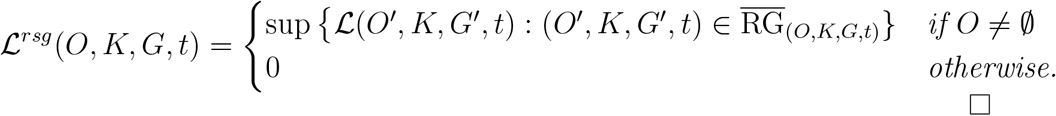

**Remark 7.3.3**. *In the previous definition we needed* 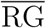 *and not* RG, *because it may happen that O* ∉ *G, and without that* **ℒ**^*rsg*^ (*O*) *would be (partially) independent from* **ℒ**(*O*).

#### Proposition 7.3.4.

*Let a group dissolvable life-level-system* Λ *be given. Let* (*O, K, G, t*) ∈ Dom **ℒ**. *Then* **ℒ**(*O, K, G, t*) ≤ **ℒ**^*rsg*^ (*O, K, G, t*). □

#### Proposition 7.3.5.

Ran 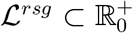 *and* **ℒ**^*rsg*^ *satisfies properties (es)*. □

#### Proposition 7.3.6.

*Let a life-level-system* Λ *be given. Suppose that* (*O, K, G, t*) ∈ Dom **ℒ** *and O*^*′*^ ⊂ *O*^*′′*^, *O*^*′*^, *O*^*′′*^ ∈ **𝒪**, *O*^*′*^ ∈ *G implies that O*^*′′*^ ∈ *G. Then* **ℒ**^*rsg*^ *satisfies properties (ss)*.

*Proof*. The condition gives that *O*^*′*^ ⊂ *O*^*′′*^ implies that 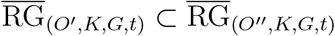. Then also using (ss) for **ℒ**, we get that claim. □

#### Proposition 7.3.7.

*Let a life-level-system* Λ *be given. Suppose that any of the following holds for* Λ.

1. *If* **ℒ**(*O, K, G, t*) = 0 *then O* = ∅.
2. Λ *is* Γ*-vanishing*.

*In this case* **ℒ**^*rsg*^ *satisfies properties (do)*.

Roughly speaking we are now going to define how long after *t*_0_ property *p* survives in the reverse-supporting-group of (*O, K, G, t*).

#### Definition 7.3.8.

*Let a propertized life-level-base-system* ⟨Λ_*b*_, **𝒫**, *w*, ∝⟩

*be given. Let* (*O, K, G, t*) ∈ Dom **ℒ** *and let p* ∈ **𝒫**. *Set*

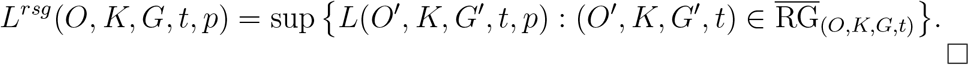

#### Proposition 7.3.9.

*Let a propertized life-level-base-system* ⟨Λ_*b*_, **𝒫**, *w*, ∝ ⟩ *be given. Then*

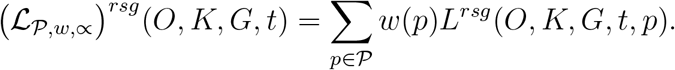

*Proof*. It can be readily seen that

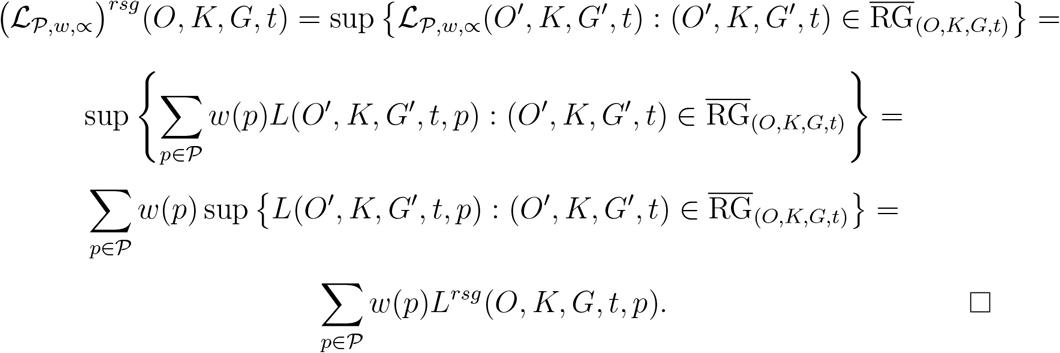

Now we provide a partially modified version of the definition 7.3.2 combined with the survival time of properties (cf. proposition 7.3.9).

#### Definition 7.3.10.

*Let a propertized life-level-system* ⟨Λ, **𝒫**, *w*, ∝⟩ *be given. Let b* = (*O, K, G, t*) *be an admissible bio-bit and O* ∈ *G*.

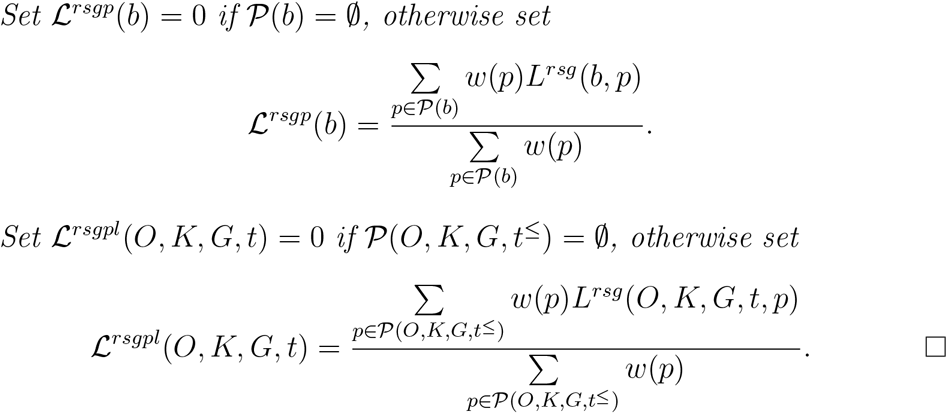

#### Proposition 7.3.11.

*With the assumptions and notations of 7.3.10*, **ℒ**^*rsgp*^(*O, K, G, t*) *and* **ℒ**^*rsgpl*^(*O, K, G, t*) *are the conditional expected values of the function X* : **𝒫** → ℝ ∪ {0}, *X*(*p*) = *L*^*rsg*^(*O, K, G, t, p*) *regarding the set* **𝒫**(*O, K, G, t*) *and* **𝒫**(*O, K, G, t*^≤^) *respectively*.

*Proof*. Obviously *w* can be considered as a probability distribution on **𝒫**.

#### Proposition 7.3.12.

*With the assumptions and notations of 7.3.10*, Ran 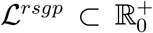 *and* **ℒ**^*rsgp*^ *satisfies property (es). Similar statements are true for* **ℒ**^*rsgpl*^. □

#### Proposition 7.3.13.

*With the assumptions and notations of 7.3.10, let O, O*^*′*^ ∈ **𝒪**, *O* ⊂ *O*^*′*^ *and* **𝒫**(*O, K, G, t*) = **𝒫**(*O*^*′*^, *K, G, t*). *Then for this O, O*^*′*^, **ℒ**^*rsgp*^ *satisfies property (ss)*. □

#### Example 7.3.14.

**ℒ**^*rsgp*^, **ℒ**^*rsgpl*^ *do not necessarily satisfy property (do), e.g. consider example 7.2.22*.

#### Lemma 7.3.15.

*With the assumptions and notations of 7.3.10, let O, O*^*′*^ ∈ **𝒪**, *O* ⊂ *O*^*′*^ *and* **𝒫**(*O*^*′*^, *K, G, t*) − **𝒫**(*O, K, G, t*) = {*p*^*′*^}. *Then* **ℒ**^*rsgp*^(*O*^*′*^, *K, G, t*) *<* **ℒ**^*rsgp*^(*O, K, G, t*) ⇔ *L*^*rsg*^(*O, K, G, t, p*^*′*^) *<* **ℒ**^*rsgp*^(*O, K, G, t*).

*Similar statement is true for* **ℒ**^*rsgpl*^.

*Proof*. Similar argument can be applied then in 7.2.30. □

#### Example 7.3.16.

**ℒ**^*rsgp*^, **ℒ**^*rsgpl*^ *do not satisfy property (ss) in general. Apply 7.3.15*.

One can also apply a similar approach for reverse-supporting-groups as we have in subsection 7.2.1. Here we just sketch the basis of that.

#### Definition 7.3.17.

*Let a life-level-system* Λ *be given together with weight functions w on the supporting-groups. Let* (*O, K, G, t*) ∈ Dom **ℒ**. *Then set*

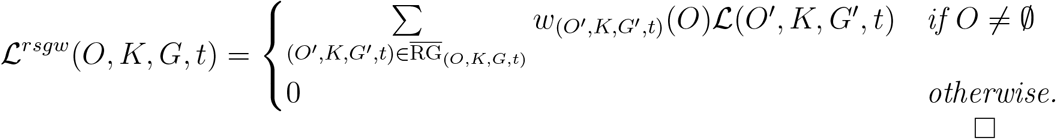

### 7.4 Life-level by development-path branching in the past

If we have an organism that has offsprings (descendants) then we have two options if we want to take into consideration that they may increase the life-level of the original organism. One option is to put them into the supporting-group and apply 7.2. The second option is to find the relatives and then identify the factor of relation and then combine those values into a new life-level function. That is what we are going to describe here.

First we need to handle when a descendant appeared in the past and we need to connect all of those to the current development-path. In this new sense, a descendant is represented by its development-path.

#### Definition 7.4.1.

*Let a life-level-system* Λ *be given and let γ* ∈ Γ, *t* ∈ **𝒯**. *We call γ*^*′*^ ∈ Γ *a* ***branch of γ at time t*** *(or t-branch of γ) if there is t*^*′*^ ≤ *t such that* 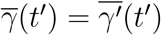. *All branches of γ is denoted by* **ℬ**_*γ*_(*t*).

We now want to categorize those linked descendants’ development-paths, or better to say, we want to assign a factor to each such path that expresses its relation to the original gorganism. □

#### Definition 7.4.2.

*Let a life-level-system* Λ *be given and let γ* ∈ Γ, *t* ∈ **𝒯**. *A* ***relation-factor*** *is a function on each development-path branch:*

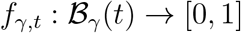

*such that f*_*γ,t*_(*γ*) = 1. □

Roughly speaking, we define a new operation in a way that we take all descendants whose relation factor is higher than a fixed value, and take the maximum life-level decreased by the factor of those.

#### Definition 7.4.3.

*Let a life-level-system* Λ *be given together with a relation-factor f*_*γ,t*_ *on each development-path branch. Let q* ∈ (0, 1). *Let b* = (*O, K, G, t*) ∈ Dom **ℒ**. *Then set* 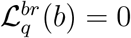 *when O* = ∅, *otherwise set*

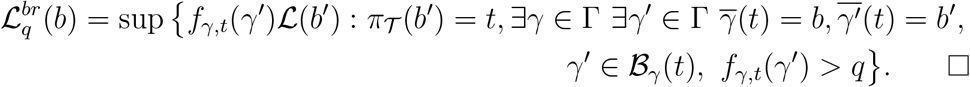

#### Proposition 7.4.4.

*Let a life-level-system* Λ *be given together with a relation-factor f*_*γ,t*_ *on each development-path branch. Let q* ∈ (0, 1). *Then* 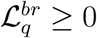 *and* 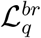 *satisfies property (es)*. □

**Remark 7.4.5**. *Clearly property (do) is not satisfied necessarily. However this phenomena was expected in this model*.

#### Proposition 7.4.6.

*Let a life-level-system* Λ *be given together with a relation-factor f*_*γ,t*_ *on each development-path branch. Let q, q*^*′*^ ∈ (0, 1). *If q < q*^*′*^ *then* 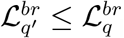.

#### Proposition 7.4.7.

*Let a life-level-system* Λ *be given together with a relation-factor f*_*γ,t*_ *on each development-path branch. Let q* ∈ (0, 1). *Suppose that b, b*^*′*^ ∈ Dom **ℒ**, *b* ⊆_*𝒪*_ *b*^*′*^, 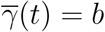 *implies that* ∃*γ*^*′*^ ∈ Γ *such that*

1. 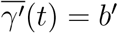 *and* 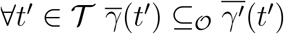 *and*
2. *γ*^*′′*^ ∈ **ℬ**_*γ*_(*t*) *implies that γ*^*′′*^ ∈ **ℬ**_*γ*_*′* (*t*) *and* 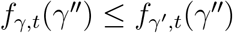.

*Then* 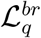 *satisfies property (ss)*.

**Remark 7.4.8**. *In condition (1) the requirement* 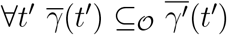 *is not needed in the proof, however it is quite natural to be added. Actually, in this form, condition (1) is that* Λ *is* Γ*-above-synchronized*.

*In condition (2), normally, one cannot expect more that f*_*γ,t*_(*γ*^*′′*^) = *f*_*γ*_*′*_,*t*_(*γ*^*′′*^), *however we formulated this more generally*.

## 8 Averages

We have already investigated average type operations e.g. when we smoothed the life-level curve (see 6.3) or when we considered moving averages (see 6.4). Now we are going to introduce and investigate some more.

### 8.1 Estimating future life-levels

Until now we thought of the life-level function which can describe the past or the present. Or also the future in the sense that we know exactly how the future looks like. But it is never the case. Here we present a more sophisticated method to assign more reasonable values to bio-bits in the future.

Throughout the paper we always emphasized that in order to determine life-level one needs all 4 components: the gorganism, the sub-environment, the supporting-group and time. None of them can be missing. It is still true for the past, the present and the future as well. Nevertheless we can get more precise values, or better to say, better estimations for the future if we calculate averages on probabilistic bases. In the general context the estimation provides future values for a gorganism, not a bio-bit. In the most general context the estimation provides future values for the current bio-bit.

#### Definition 8.1.1.

*Let a life-level-system* Λ *be given. Let t* ∈ **𝒯**, *O ∈* **𝒪**(*t*), *K* ∈ **𝒦** (*t*), *G* ∈ **𝒢** (*t*). *Let t be a future point in time*.

*If there is a non-zero probability that O can exists in the sub-environment K at time t, then we denote this fact by O* ≺_*t*_ *K and its probability by* P(*O* ≺_*t*_ *K*).

*If there is a non-zero probability that the group G acts as a supporting-group for O at time t, then we denote this fact by O* ≺_*t*_ *G and its probability by* P(*O* ≺_*t*_ *G*).

*Here of course we understand probability according to our current knowledge at time t*_0_ *that denotes the current moment. If we want to include that in the notation we use* 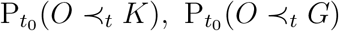. □

Let a life-level system 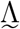 be given in the general context. Let *t*_0_ denote the current moment and let *t*_1_ *> t*_0_ be some future point in time. We do not know the future, however we want an estimate for the life-level of our gorganism *O* ∈ **𝒪** at time *t*_1_. To achieve that goal we can calculate the expected value of 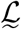 for *O* at *t*_1_.

We also provide two more specific versions where besides *O*, either *K* or *G* is fixed as well. In other words we want to estimate the life-level of our gorganism *O* ∈ **𝒪** at time *t*_1_ when the sub-environment *K* (or the supporting-group *G*) is intact through that period of time.

#### Definition 8.1.2.

*Let a countable life-level system* Λ *be given in the general context and let t*_0_, *t*_1_ ∈ **𝒯**, *t*_1_ *> t*_0_. *Set*

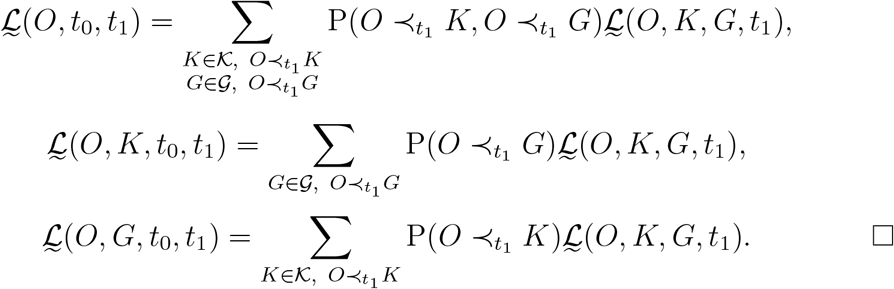

**Remark 8.1.3**. *We deliberately used the same symbol* 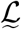 *for these functions hoping that it will not cause confusion because the parameter list uniquely identifies the corresponding function*.

**Remark 8.1.4**. *Instead of the countablity condition, one may suppose that there are finitely or countably many sub-environments such that* 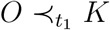 *holds and similarly there are finitely or countably many groups such that* 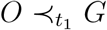 *holds*.

Now we provide estimation in the most general case.

#### Definition 8.1.5.

*Let a life-level-system* Λ *be given. If γ* ∈ Γ, *t*_0_, *t*_1_ ∈ **𝒯**, *t*_0_ *< t*_1_ *then* P(*γ, t*_0_, *t*_1_) *will denote the probability that γ gives correctly how the bio-bit γ*(*t*_0_) = *γ*_*𝒪*_(*t*_0_), *γ*_**𝒦**_(*t*_0_), *γ*_***𝒢***_(*t*_0_) *evolves to the bio-bit γ*(*t*_1_) = *γ*_*𝒪*_(*t*_1_), *γ*_**𝒦**_(*t*_1_), *γ*_***𝒢***_(*t*_1_) *from time t*_0_ *to t*_1_.

*Again we understand probability according to our current knowledge at time t*_0_ *that denotes the current moment. If we want to include that in the notation we use* 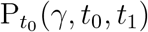.

Suppose now that we have a life-level system Λ in the most general context and we have a bio-bit (*O, K, G, t*_0_) ∈ Dom **ℒ** where *t*_0_ denotes the current moment and let *t*_1_ *> t*_0_. We want to estimate the life-level of the bio-bit that evolves from our original bio-bit at time *t*_1_. Again we calculate the expected value of **ℒ** for (*O, K, G*) at *t*_1_.

#### Definition 8.1.6.

*Let a countable life-level system* Λ *be given and let* (*O, K, G, t*_0_) ∈ Dom **ℒ**, *t*_0_, *t*_1_ ∈ **𝒯**, *t*_1_ *> t*_0_. *Set*

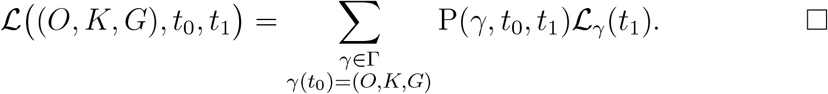

#### 8.1.1 Operation type averages

We have just defined some functions that can be considered as operations on life-level functions to some extent.

##### Definition 8.1.7.

*Let a countable life-level system* 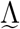 *be given in the general context. Using the notation of definition 8.1.2 set*

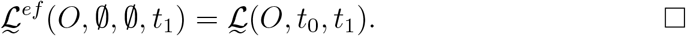

##### Proposition 8.1.8.

*Let a countable life-level system* 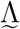 *be given in the general context. Then* 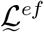 *satisfies property (es)*.

*Moreover suppose t hat O* ⊂ *O*^*′*^ *implies that* 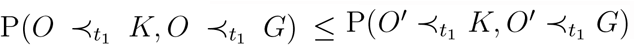. *Then* 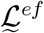 *satisfies property (ss)*.

*Proof*. Property (es) trivially holds. For property (ss) apply property (ss) on **ℒ**.

**Remark 8.1.9**. *Evidently the condition* 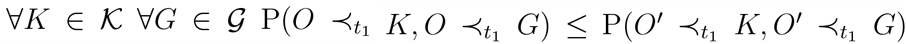 *is equivalent with* 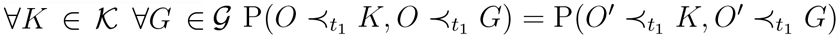.

**ℒ**

**Remark 8.1.10**. *Without any additional condition one cannot expect that* 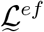 *satisfies property (do). In order to force that one can apply e.g. the operation* ^*do*^ *(see 6.2)*.

### 8.2 Averages on time

First we can simply calculate the average of a life-level function on a given time interval.

#### Definition 8.2.1.

*Let a life-level-system* Λ *be given. Let* [*a, b*] ⊂ **𝒯**, *γ* ∈ Γ. *Then the average of* **ℒ** *through γ on the interval* [*a, b*] *is*

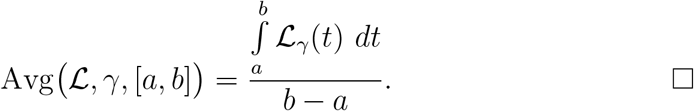

It may be more interesting to examine the average life-level of a gorganism on its whole life-span.

#### Definition 8.2.2.

*Let a life-level-system* Λ *be given such that* ***𝒯*** *is a closed interval of ℝ. Let γ* ∈ Γ. *Then the average of* **ℒ** *on the whole life-span determined by γ is*

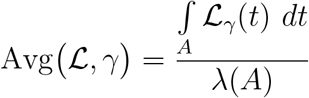

*where A* = {*t* ∈ **𝒯** : **ℒ**_*γ*_(*t*) *>* 0} *and λ*(*A*) *is the length of A*.

**Remark 8.2.3**. *In 2.2.2 we showed that {t* ∈ ***𝒯***: **ℒ**_*γ*_(*t*) *>* 0} *is always an interval*.

We now define average for a given bio-bit.

#### Definition 8.2.4.

*Let a life-level-system* Λ *be given such that* ***𝒯*** *is a closed interval of ℝ. Let b* ∈ Dom **ℒ**. *Let a probability function* P *be given on* Γ_*b*_. *Then set*

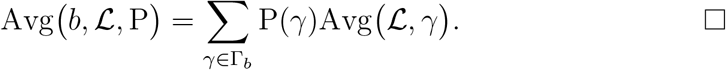

### 8.3 Assigning values to groups

In this subsection we will assign life-level type values to groups of gorganisms, e.g. species. But the usual warnings are valid here as well. The warnings are valid on two levels, an old and a new one. First, there are some other parameters that cannot be switched off completely, e.g. at least parameter “time” should remain. Second, we want to keep the possibility that we do not just assign one single value to a group, instead we assign a (finite) multiset of numbers to the group.

So one should not think that we are now going to assign values to species. However we are getting closer to that, but only closer. We will do that in section 9, not here.

#### Definition 8.3.1.

*A life-level-system* Λ *is called* ***single-valued*** *if* ∀*t* ∈ **𝒯** ∀*O* ∈ **𝒪**(*t*) (*O, K, G, t*), (*O, K*^*′*^, *G*^*′*^, *t*) ∈ Dom **ℒ** ⇒ *K* = *K*^*′*^, *G* = *G*^*′*^.

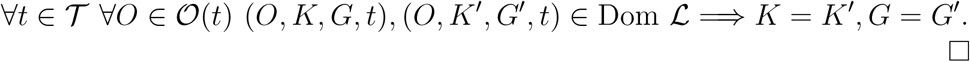

**Remark 8.3.2**. *The previous condition can also be formalized in a way that* ∀*t* ∈ **𝒯** ∀*O* ∈ **𝒪**(*t*) *there is at most one K, G pair such that* **ℒ** *assigns value to* (*O, K, G, t*). *I.e. for a given gorganism the model does not want to take in account different sub-environments / supporting groups at the same time*.

#### Definition 8.3.3.

*Let a single-valued finite life-level-system* Λ *be given. Let t* ∈ **𝒯**, *F* ⊂ **𝒪**(*t*). *Then set*

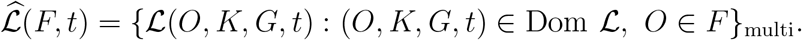

*If K* ∈ **𝒦**, *G* ∈ **𝒢** *are fixed, then set*

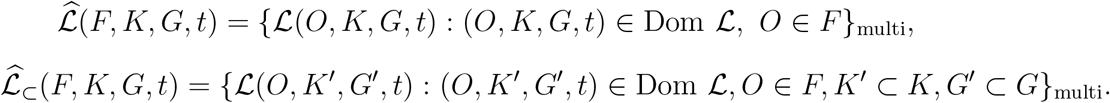

*We call these multisets the* ***life-level mset****s of F*. □

**Remark 8.3.4**. *Let f be any statistical “average”-type function that acts on finite multisets. If* Λ *is a single -valued finite life-level-system, t* ∈ **𝒯**, *F* ⊂ **𝒪**(*t*), *then one can calculate* 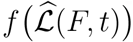, *and use it as one single value that somehow expresses the life-level of the whole group*.

*It is important to note that* 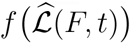 *is a two-variable function i.e. this generalized life-level value depends on F and t. But if we think that in the short term F is fix, then the value is still time dependent. I.e. if our aim is to assign a value to all members of a species (i.e. one value to the group), then the value may vary through time*.

We can get a partially different type of average if we have a kind of weight on the elements of the group.

#### Definition 8.3.5.

*Let a single-valued finite life-level-system* Λ *be given. Let t* ∈ **𝒯**, *F* ⊂ **𝒪**(*t*). *Let μ* : *F* → [0, 1] *be a weight function. Then set*

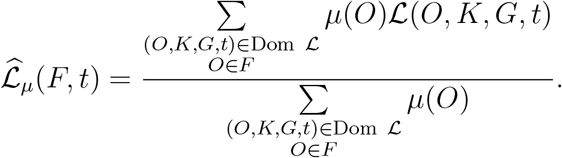

*If K* ∈ **𝒦**, *G* ∈ **𝒢** *are fixed, then* 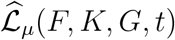 *is defined similarly*. □

If we have a (finitely additive) measure on (*t*), then we can get another type of average. We define it in the simplest case only.

#### Definition 8.3.6.

*Let a life-level-system* Λ *be given in the less general context. Let t* ∈ **𝒯**, *F* ⊂ **𝒪**, *K* ∈ **𝒦**, *G* ∈ **𝒢**. *Let μ be a (finitely additive) measure on* **𝒪**. *Then set*

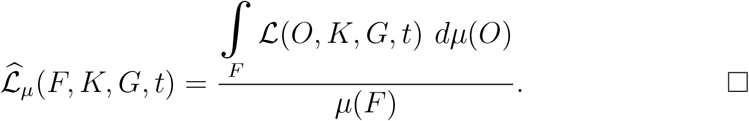

**Remark 8.3.7**. *Let* dist(.,.) *be any metric on finite multi-subsets of ℝ (see e.g. [5])*. *If* Λ *is a single-valued finite life-level-system, t* ∈ **𝒯**, *F*_1_, *F*_2_ ⊂ **𝒪** (*t*), *then one can calculate* dist 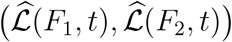, *and consider it as the life-level distance between the two groups*.

*Another option is that if a statistical “average”-type function f is given, then* 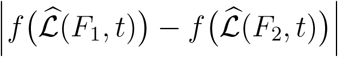 *can serve as a distance*.

Now we compare the life-level msets of groups. Roughly, we say that one is smaller than the other, if most of the elements are smaller.

#### Definition 8.3.8.

*Let a single-valued finite life-level-system* Λ *be given. Let t* ∈ **𝒯**, *F*_1_, *F*_2_ ⊂ **𝒪**(*t*). *We say that the life-level msets of F*_1_ *is smaller than the life-level msets of F*_2_, *if* 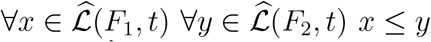 *holds. In this case we use the notation* 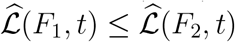.

*Let q* ∈ (0, 1). *We say that the life-level msets of F*_1_ *is smaller than the life-level msets of F*_2_ *by factor q, if* 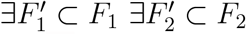 *such that* 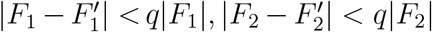 *and* 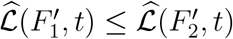 *holds. In this case we use the notation* 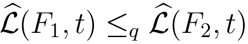.

## 9 Assigning life-level value to species

In this section, we are going to make some effort to add life-level value to species. Better to say, we will outline methods to produce, derive life-level values from already existing life-level functions (systems). We will provide two such ways and sketch a third one. The first one simply takes the maximum life-level of certain bio-bits, while the second one is based on selecting the most viable organisms from a species and use the average of their life-level values. In both methods, the time parameter cannot be excluded fully.

It is important to mention that here we are not going to identify, define what a species is. Instead, we assume that the given application already has a categorization on certain organisms, i.e. some distinct groups of organisms are considered to be species by the application, but how it does that, it is completely up to the application.

The very basic version simply takes the maximum life-level in the group. However it is not enough to take that at a given time point, since e.g. the whole species can be at a dormant state (e.g. in winter or in an ice age) and that is not what we are interested in. Therefore we have to take into account a certain time interval around the given time point and take the maximum there. That interval length can be dependent on the species and the base time point as well, so it cannot be chosen independently.

### Definition 9.0.1.

*Let a life-level-system* <u>Λ</u> *be given at the less general context and let G* ∈ **𝒢**, *t*_0_ ∈ **𝒯**, 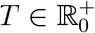. *Set*

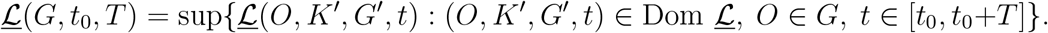

**Remark 9.0.2**. *In some sense we can interpret the previous definition that we take the best sub-environments and the best supporting-groups from those which are applicable for the members of the group at the given time interval, and make investigation for those bio-bits*.

In the most general context, we have to take into account the development-paths of the group.

### Definition 9.0.3.

*Let a life-level-system* Λ *be given and let G* ∈ **𝒢**(*t*_0_) ∩ **𝒪**(*t*_0_), *t*_0_ ∈ **𝒯**, 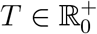 *Set*

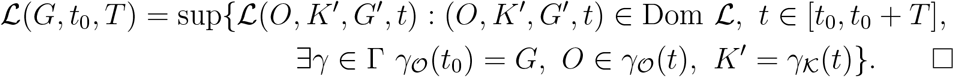

**Remark 9.0.4**. *In definition 9.0.3 we did not require the existence of a γ*^*′*^ ∈ Γ *such that γ*^*′*^ (*t*) = *O and* 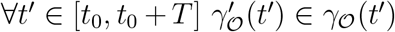, *i.e. we do not require that the organism stays in the group at every time point. Instead, we just take a development-path of the group G at a given time point t, and find all possible elements*.

For the second method, we need a concept that also covers the notion of “mainstream” gorganism, that is we want to select the most viable organisms from a species which can “truly” represent the species itself. We will define a much more general concept here.

### Definition 9.0.5.

*Let a life-level-subbase-system* Λ_*sb*_ *be given and let* **𝒫** *be a property set. We call b* ∈ Θ *a* ***𝒫-bio-bit*** *if* ∀*p* ∈ **𝒫** *p* ∝ *b holds. We also use the notation* **𝒫** ∝ *b in this case*. □

### Example 9.0.6.

*Of course, the “mainstream” organisms of a species can be defined several ways by an application. Here we provide a sketch of one possible way. Let* **𝒫** *consists of the following properties:*

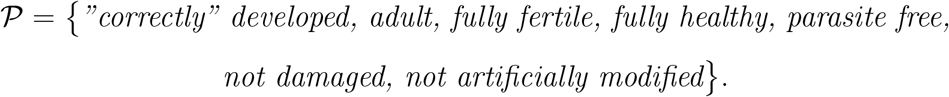

*Then the set of all* **𝒫***-bio-bits can be considered as the set of “mainstream” organisms*.

Now we define the second type of life-level assignment to species. It is based on the average life-level value of the “mainstream” gorganisms of the group in the time interval. We define it in the less general context only; one can handle the most general context similarly to definition 9.0.3.

### Definition 9.0.7.

*Let a finite life-level-system* Λ *be given at the less general context and let* **𝒫** *be a property set, let G* ∈ **𝒢**, *t*_0_ ∈ **𝒯**, 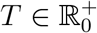 *Set*

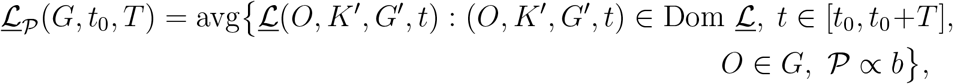

*where* avg *is the usual average of some set of numbers*.

Now we sketch a third possible way of life-level assignment to species, but we can just describe it very roughly because it is not clear how one could formalize this one. We remark that this is the only place in the paper, where we consider abstract organisms, everywhere else we refrain from doing so.

### Definition 9.0.8.

*Suppose that we have a life-level-system where the life-level function is based on some principles, i.e. the life-level is always calculated from some attributes of a bio-bit, there are no ad hoc values. If we have a group of organisms, e.g. species, then one can imagine an idealized organism from that group, an organism that owns all possible positive properties of those organisms. Moreover one can add the best possible sub-environment and the best possible supporting group that one can imagine. And then apply the life-level calculation method for that idealized bio-bit. What one gets can serve as a life-level value for the given species*.

**Remark 9.0.9**. *The question arises if one single such “super” idealized member is enough to determine the life-level, because in some cases more than one member may be needed, e.g. one may need an idealized male and an idealized female as well, because they may have different life-level values*.

**Remark 9.0.10**. *We have said that we do not define here what a species is, however we make an important comment*.

*A given ant species consists of colonies and not individual ants. An individual ant belongs to something (a colony) from which one can ask that which species it is. It is a wrong question to ask which species a single individual ant belongs to. A single ant does not belong to any species, its colony does. A single ant belongs to something (its colony) which belongs to a species*.

*Similarly, one would not ask about a cell of a mouse or an organ of a mouse that which species does it belong to. A mouse belongs to some species, but it is a wrong question to ask that about one of its cells*.

*Therefore it is important to choose carefully the gorganisms for which we apply our methods*.

## 10 Metrics in accordance with a life-level function

### 10.1 Synchronized distances

A research project might have a kind of distance on the set of all gorganisms (and/or sub-environments and/or supporting-groups) that somehow measures how far two such objects (gorganisms / sub-environments / supporting-groups) are from each other in some biological sense, e.g. some type of genetic/phenotypic distance between individuals/groups (see e.g. [25], [24]). One can have separate (pseudo-)metrics on **𝒪, 𝒦, 𝒢** and also on **𝒯**, but instead of that, or in addition, one can have a metric on the set of all bio-bits i.e. on Θ(Λ) or Dom **ℒ**. Of course one can create a metric on Θ(Λ) from metrics on **𝒪, 𝒦, 𝒢, 𝒯** and vice versa (by some usual method).

If the project also maintains a life-level function, then it is worth asking if the distance and the life-level function are synchronized, meaning that closer objects provide closer life-level values. More precisely the closer the objects are, the closer the associated life-level values are. Here we are going to formalize this notion.

#### Definition 10.1.1.

*Let a life-level-system* Λ *be given together with a pseudo-metric d on* Dom **ℒ**. *We say that the associated life-level function* **ℒ** *and the pseudo-metric d are* ***synchronized*** *if*

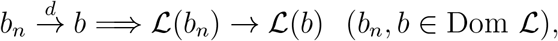

*they are* ***uniformly-synchronized*** *if*

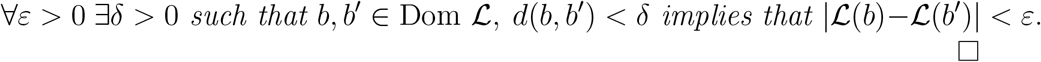

□

**Remark 10.1.2**. *An equivalent form of synchronized* **ℒ** *and d is*

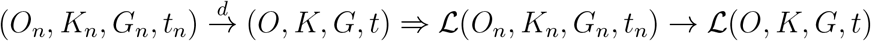

*where* (*O*_*n*_, *K*_*n*_, *G*_*n*_, *t*_*n*_), (*O, K, G, t*) ∈ Dom **ℒ**.

*Or if d* = *d*_𝒪_ + *d*_**𝒦**_ + *d*_**𝒢**_ + *d*_**𝒯**_, *where d*_𝒪_, *d*_**𝒦**_, *d*_**𝒢**_, *d*_**𝒯**_ *are pseudo-metrics on* **𝒪, 𝒦, 𝒢, 𝒯** *respectively, then the equivalent form is*

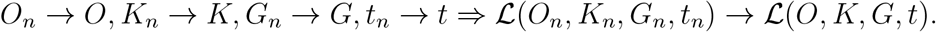

#### Proposition 10.1.3.

*Let a life-level-system* Λ *be given together with a pseudo-metric d on* Dom **ℒ**. *Then* **ℒ** *and d are synchronized iff* **ℒ** *is a continuous function (regarding the pseudo-metric space* ⟨Dom **ℒ**, *d*⟩).□

We define some weaker forms where we have a pseudo-metric on **𝒪** only (or **𝒦** or **𝒢**).

#### Definition 10.1.4.

*Let a life-level-system* Λ *be given together with a pseudo-metric d*_*𝒪*_ *on* **𝒪**. *We say that the associated life-level function* **ℒ** *and the pseudo-metric d*_*𝒪*_ *are* ***synchronized*** *if*

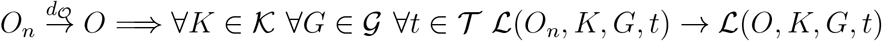

*where* (*O*_*n*_, *K, G, t*), (*O, K, G, t*) ∈ Dom **ℒ**.

*We say that* **ℒ** *and d*_*𝒪*_ *are* ***uniformly-synchronized*** *if*

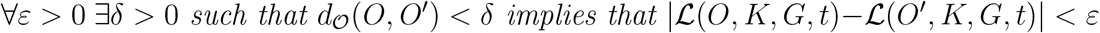

*where* (*O, K, G, t*), (*O*^*′*^, *K, G, t*) ∈ Dom **ℒ**.

*We similarly define when* **ℒ** *and d*_***𝒦***_ *(or d*_***𝒢***_ *) are (uniformly-)synchronized where d*_***𝒦***_, *d*_***𝒢***_ *are pseudo-metrics on* **𝒦, 𝒢** *respectively*. □

#### Proposition 10.1.5.

*Let a life-level-system* Λ *be given together with pseudo-metrics d*_𝒪_ *on* **𝒪**, *and d on* Dom **ℒ**. *If* **ℒ** *and d*_𝒪_ *are uniformly-synchronized, then they are synchronized. If* **ℒ** *and d are uniformly-synchronized, then they are synchronized as well*. □

### 10.2 Distances derived from a life-level function

If no such pseudo-metric(s) are given on these sets, then we can also investigate how a given life-level function can produce a (synchronized) pseudo-metric on these sets as well. We will present several such pseudo-metrics.

#### Definition 10.2.1.

*Let a life-level-system* Λ *be given. For the associated life-level function* **ℒ** *and two bio-bits b*_1_, *b*_2_ ∈Dom **ℒ** *set*

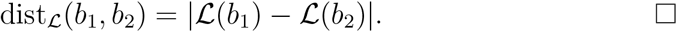

#### Proposition 10.2.2.

*The function* dist_*ℒ*_ (*b*_1_, *b*_2_) *is a synchronized pseudometric on* Dom **ℒ**. □

#### Definition 10.2.3.

*Let a life-level-system* Λ *be given. For O*_1_, *O*_2_ ∈ **𝒪**

*Set*

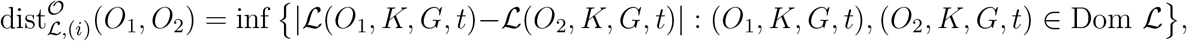

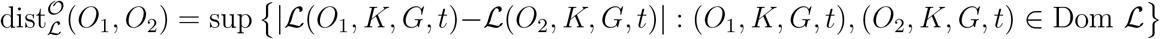

*if the above sets are not empty*.

**Remark 10.2.4**. *One can similarly define distance between sub-environments or supporting-groups starting from a life-level function*.

#### Proposition 10.2.5.

*Let a life-level-system* Λ *be given. Then* 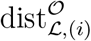*and* 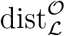 *are pseudo-metrics, moreover* 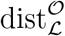 *is also uniformly-synchronized*.

*Proof*. We just show the triangle inequality for 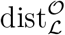(for 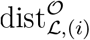it is similar).

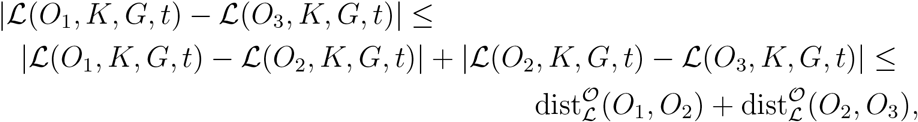

and taking the supremum on the left hand side gives the claim. □

**Remark 10.2.6**. *In some sense* 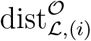 *is more natural for measuring distance between two gorganisms, however it is not necessarily synchronized*.

#### Proposition 10.2.7.

*Let a life-level-system* Λ *be given with a pseudometric d*_𝒪_ *on* **𝒪**. *Then d*_*𝒪*_ *and* **ℒ** *are uniformly-synchronized if* 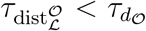 *i.e. the generated topology by d*_*𝒪*_ *is finer than the generated topology by* 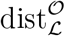.

*Proof*. It is an obvious consequence of the fact that

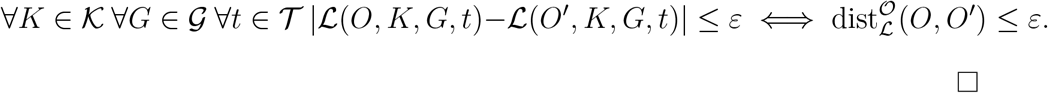

**Remark 10.2.8**. *We make a comment on time regarding distances derived from a life-level function*.

*If* **ℒ** *is a life-level function in the most general context, then in the definition of* 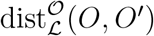, *O, O’* ∈ **𝒪** (*t*_0_) *for some t*_0_ ∈ **𝒯**, *i.e. they exist at the same moment t*_0_ *(and similarly K and G exist at t*_0_*). In other words our measurement restricted to time points, by that method we cannot compare objects existing further in time. (The reason for that is that in* 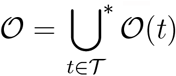

*there is a disjoint union, similarly for* ***𝒦, 𝒢***.)

*In the general context, of course there is no such restriction*.

*The distance* dist_ℒ_ *can measure distance between bio-bits that exist at different time points*.

## 11 Measuring vitality of sub-organisms

Here we build a small, basic theory about how to measure the vitality of sub-organisms of an organism, in other words we want to measure how much a given (generalized) sub-organism is important for the life-level of a containing (generalized) organism. As an example, one can think of how to characterize the vital organs of a body, or more generally, how one can measure the different such vitality levels of organ systems, organs, tissues, etc. regarding the body, or also one sub-system regarding a containing system. Another example is when one examines the vitality of a member of a group of wolfs regarding to the whole group (e.g. the alpha male).

First we will build some generic notions related to such type of measurements, and then we show a way how one can create such type of measurement from a given life-level function (better to say life-level-system). We will also investigate the connections between the properties of the vitality-measurement and the corresponding properties of the life-level function.

In the sequel, if a lattice **𝒪** is given with a least element, then we will denote the infimum, supremum in **𝒪**, the order relation on **𝒪** and the least element by ∩, ∪, ⊂, ∅ respectively.

### Definition 11.0.1.

*Let a lattice* ⟨ **𝒪**, ∩, ∪, ⊂⟩ *be given with a least element* ∅. *We call a function*

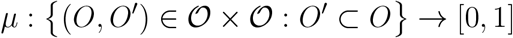

*a* ***vitality-measurement*** *on* **𝒪** *if it satisfies the following properties*.

1. *μ*_*𝒪*_(∅) = 0, *μ*_*𝒪*_(*O*) = 1
2. *O*_1_ ⊂ *O*_2_ ⊂ *O* =⇒ *μ*_*𝒪*_(*O*_1_) ≤ *μ*_*𝒪*_(*O*_2_)
3. *μ*_*𝒪*_(*O*_1_ ∪ *O*_2_) ≤ *μ*_*𝒪*_(*O*_1_) + *μ*_*𝒪*_(*O*_2_) (*O*_1_, *O*_2_ ⊂ *O*)
4. 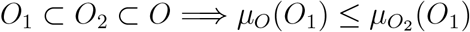

*where we use the notation μ*_*𝒪*_(*O′*) *for μ*(*O, O′*), *and it is called the vitality of O′ regarding O*.

*We say that O′* ⊂ *O is* ***vital*** *for O if μ*_*𝒪*_(*O′*) = 1. □

**Remark 11.0.2**. *We make some technical remarks:*

- *As we mentioned in the definition, we divided μ into sub-functions μ*_*𝒪*_ *by the first component of μ. In other words, it can be considered that we have a vitality-measurement for each O* ∈ **𝒪**. *The only required connection between distinct μ*_*𝒪*_*-s is presented in condition (4)*.
- *It can be seen by induction that condition 11.0.1 (3) is equivalent with the subadditivity of* 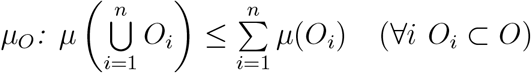.
- *Condition 11.0.1 (3) implies that μ*_*𝒪*_ ≥ 0 *(Set O′* = *O*_1_ = *O*_2_ ⊂ *O)*.

**Remark 11.0.3**. *The function μ is supposed to express how much an “inner” part gorganism is important for the life-level of a containing gorganism. Clearly 11.0.1 (1), (2) and (3) are straightforward expectations. Condition (4) says that the same component is more important for a smaller gorganism than for a larger one*.

*Obviously one cannot await more than subadditivity i.e. additivity is not an expectable property of such a function*.

### 11.1 Deriving vitality-measurement from life-level function

If a life-level function is added with some additional assumptions, then one can create a vitality-measurement with the help of that.

First we will need the general notion of subtraction in a lattice.

#### Definition 11.1.1.

*Let a lattice* ⟨**𝒪**, ∩, ∪, ⊂⟩ *be given with a least element. We say that the binary operation* − *is a subtraction on* **𝒪** *if it is function like* **𝒪** × **𝒪** → **𝒪** *such that it satisfies*

1. (*O*_1_ − *O*_2_) ∩ *O*_2_ = ∅
2. (*O*_1_ − *O*_2_) ∪ (*O*_1_ ∩ *O*_2_) = *O*_1_. □

**Remark 11.1.2**. *We just make three technical remarks*.

1. *Condition 2 implies that O*_1_−*O*_2_⊂*O*_1_ *and* (*O*_1_−*O*_2_) ∪*O*_2_ = *O*_1_∪*O*_2_. *Moreover these two properties together are equivalent with condition 2*.
2. *Condition 2 yields that if O*_1_ ∩ *O*_2_ = ∅, *then O*_1_ − *O*_2_ = *O*_1_.
3. *If the lattice is modular, then the subtraction is unique (if exists)*.

**Remark 11.1.3**. *One may wonder why we need such general notion of subtraction here, because if every gorganism is represented by a set, then the subtraction can be simply the set theoretical subtraction and that is it. In some cases, yes. However there can be cases where removing a sub-gorganism from a host gorganism cannot be that simple that we just cut it off, because cutting would destroy immediately the host gorganism; instead removing means some “intelligent” way of cutting, that usually means that one has to modify some remaining connected parts in order to the host remain functioning*.

*Furthermore, in some models the gorganisms are not represented by sets (subsets of a bigger set), and in those cases, we have to specify which properties we expect from a subtraction operation*.

#### Definition 11.1.4.

*Let* Λ *be a life-level-system in the general context such that* ⟨⟨**𝒪**, ∩,∪,⊂,−⟩ *is a lattice with a least element and with subtraction operation such that*

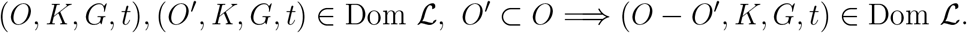

*If* (*O, K, G, t*) ∈ Dom **ℒ, ℒ** (*O, K, G, t*) ≠ 0 *and O′* ⊂ *O, then set*

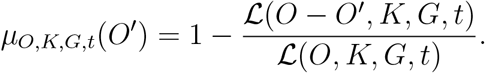

*When K, G, t are clear from the context, then we will use μ*_*𝒪*_(*O′*) *simply*. □

**Remark 11.1.5**. *We remark that μ*_*𝒪*_ *is not defined when* **ℒ** (*O, K, G, t*) = 0, *but it is obviously not needed in applications*.

#### Proposition 11.1.6.

*The function μ*_*𝒪*_ *defined in 11.1.4 satisfies condition (1) and (2) in the definition of vitality-measurement (11.0.1)*.

*Proof*. (1) *μ*_*𝒪*_(*O*) = 1 comes from **ℒ** (∅, *K, G, t*) = 0. *μ*_*𝒪*_(∅) = 0 is obvious.

(2) If *O*_1_ ⊂ *O*_2_ ⊂ *O*, then *O* − *O*_2_ ⊂ *O* − *O*_1_ which yields that

**ℒ** (*O* − *O*_2_, *K, G, t*) ≤ **ℒ** (*O* − *O*_1_, *K, G, t*) by property (ss) of **ℒ**. Which gives that *μ*_*𝒪*_(*O*_1_) ≤ *μ*_*𝒪*_(*O*_2_). □

#### Proposition 11.1.7.

*The condition that*

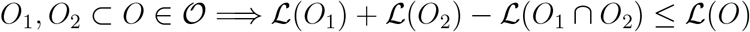

*is equivalent with condition 11.0.1 (3) for μ*_*𝒪*_ *defined in 11.1.4*.

*Proof*. For *O*_1_, *O*_2_ ⊂ *O* ∈ **𝒪** and fixed *K* ∈ **𝒦**, *G* ∈ **𝒢**, *t* ∈ **𝒯** clearly

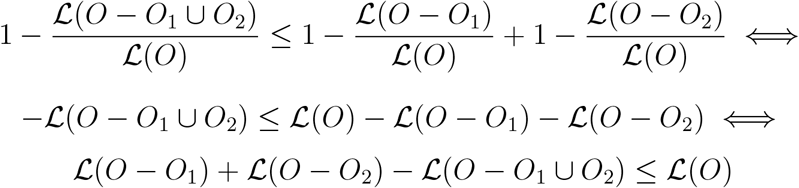

and that is equivalent with the condition here. □

#### Proposition 11.1.8.

*The condition that*

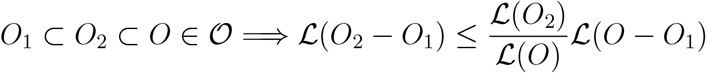

*is equivalent with condition 11.0.1 (4) for μ*_*𝒪*_ *defined in 11.1.4*.

*Proof*. For *O*_1_ ⊂ *O*_2_ ⊂ *O* ∈ **𝒪** and fixed *K* ∈ **𝒦**, *G* ∈ **𝒢**, *t* ∈ **𝒯** clearly

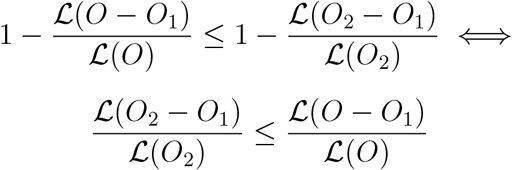

and that is equivalent with the condition here. □

#### Proposition 11.1.9.

*If subadditivity (condition 11.0.1 (3)) holds for functions μ*_*𝒪*_ *defined in 11.1.4 then*

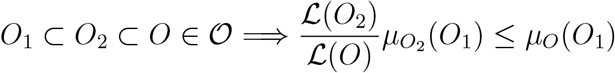

*holds*.

*Proof*. Let *O*_1_⊂*O*_2_⊂*O* ∈ **𝒪**. Applying 11.0.1 (3) for *O*_1_ and *O*−*O*_2_ we get that

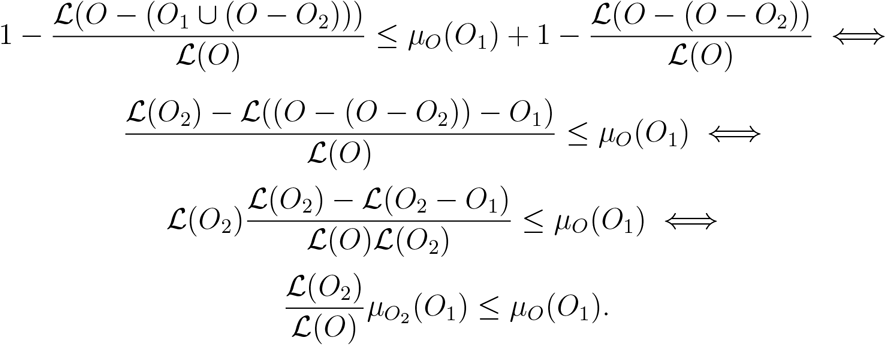

□

#### Corollary 11.1.10.

*If μ defined in 11.1.4 is a vitality-measurement*,

*O*_2_ ⊂ *O* ∈ **𝒪** *and* **ℒ** (*O*_2_) = **ℒ** (*O*), *then*

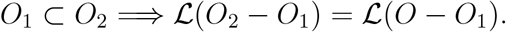

*Proof*. By 11.1.9 we get that 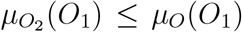, while condition (4) gives the opposite 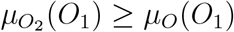, hence 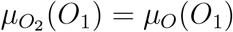, that is

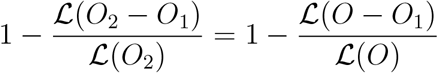

which yields the claim.□

#### Corollary 11.1.11.

*With the assumption of the previous corollary 11.1.10*,

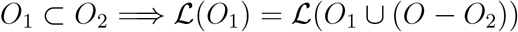

*also holds. Moreover this is an equivalent condition*.□

#### Proposition 11.1.12.

*Let* Λ *be a life-level-system in the general context such that*, ⟨**𝒪**,∩, ∪, ⊂, −⟩*is a lattice with a least element and with subtraction operation such that*

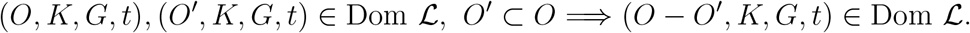

*If* **ℒ** *is additive, then μ*_*𝒪*_ *defined in 11.1.4 is a vitality-measurement*.

*Proof*. Condition (1) and (2) are established in 11.1.6.

Condition (3) comes from 11.1.7, because additivity gives that

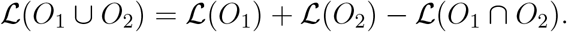

And *O*_1_ ∪ *O*_2_ ⊂ *O* implies that **ℒ** (*O*_1_ ∪ *O*_2_) ≤ **ℒ** (*O*) by property (ss) of **ℒ**. For condition (4), let *O*_1_ ⊂ *O*_2_ ⊂ *O* ∈ **𝒪**. Set *A* = *O*_1_, *B* = *O*_2_ − *O*_1_, *C* =*O* − *O*_2_. By 11.1.8 and using additivity of **ℒ**, we have to show that

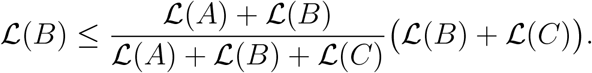

That is

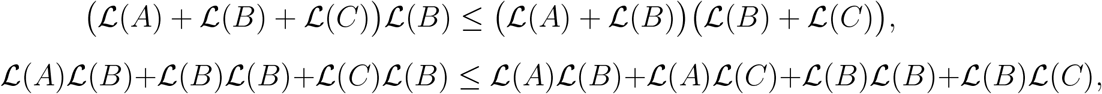

which holds since 0 ≤ **ℒ** (*A*) **ℒ** (*C*).

**Remark 11.1.13**. *In contrast with definition 11.1.4, an alternative way of creating a vitality-measurement from a life-level function would be the following (under similar assumptions on* **𝒪** *and* **ℒ**).

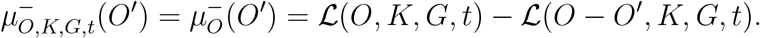

*However it has some drawbacks. E.g. subadditivity (condition 11.0.1 (3)) would be equivalent with* 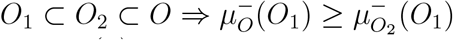 *which is a kind of opposite of condition 11.0.1 (4), and sounds quite unnatural and against our expectation*.

*Compare identity in proposition 11.1.9 which can be acceptable*.

**Remark 11.1.14**. *One can define functions μ*_*𝒪*_ *like in definition 11.1.4 if* Λ *is more general, i.e. it is a life-level-system in the most general context. In that case, we need assumptions for***𝒪**(*t*) *for each t* ∈ **𝒯** *like in definition 11.1.4, and one can have functions μ*^(*t*)^ *defined on* **𝒪** × (*t*) **𝒪** (*t*) *for each t∈* ***𝒯***.

*In such a case, one can examine the change of vitality through development-path. But here two development-paths are needed, one for O and one for O′ (O* ⊂ *O). Let γ, γ′* ∈ **Γ** *such that γ* (*t*_0_) = (*O, K, G*), *γ′*(*t*_0_) = (*O′, K, G*), *and* ∀*t > t*_0_ *π*_𝒪_(*γ* (*t*)) ⊂ *π*_𝒪_(*γ′*(*t*)), *π* _***𝒦***_ (*γ* (*t*)) = *π* **𝒦**(*γ′*(*t*)), *π* **𝒢** (*γ* (*t*)) = *π* _***𝒢***_ (*γ′*(*t*)). *Then the change of vitality in time of O′ regarding O through γ, γ ′ is described by*

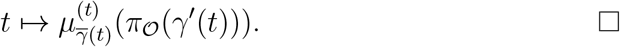

#### 11.2 Deriving life-level function from vitality-measurement

Now we deal with the opposite: How to create a life-level function from a given vitality-measurement. We will provide one such way (it will be easy to see that many can exist).

Here we deal with partially ordered sets only instead of lattices. For those, we also need the notion of subtraction.

##### Definition 11.2.1.

*Let a partially ordered set* ⟨ **𝒪**, ⊂⟩ *be given. We say that the binary operation* − *is a subtraction on* **𝒪** *if it is function like* **𝒪** × **𝒪** → **𝒪** *such that it satisfies*

1. (*O*_1_ − *O*_2_) ⊂ *O*_1_
2. *O* ⊂ (*O*_1_ − *O*_2_) =⇒ *O* ⊄*O*_2_. □

##### Proposition 11.2.2.

*Let a partially ordered set*⟨**𝒪**,⊂,∅⟩ *be given with a least element which satisfies the followings*.

1. ∀*O* ∈ **𝒪** ∃*M* ∈ **𝒪** *maximal element such that O* ⊂ *M*.
2. *If for O*_1_, *O*_2_ ∈ **𝒪** ∃*O* ≠ ∅ *such that O* ⊂ *O*_1_, *O* ⊂ *O*_2_, *then* ∃*O*_3_ ∈ **𝒪** *such that O*_1_ ⊂ *O*_3_, *O*_2_ ⊂ *O*_3_.

*Then* ∀*O* ∈**𝒪**, *O* ≠∅ *there is a unique M* ∈ **𝒪** *maximal element such that O* ⊂ *M*.

*Proof*. Indirect, using condition 2.

##### Definition 11.2.3.

*Let* Λ_*b*_ *be a life-level-base-system in the general context such that* ⟨**𝒪**, ⊂, ∅, −⟩ *is a partially ordered set with a least element, and with subtraction operation, and* ∀*O* ∈ **𝒪**, *O* ≠ ∅ *there is a unique M* ∈ **𝒪** *maximal element such that O* ⊂ *M. Let us denote the maximal element belonging to O by M*^*O*^. *Let a vitality-measurement μ be given on* **𝒪**.

*If* (*O, K, G, t*) ∈ Θ (Λ_*b*_), *then set* **ℒ** (∅, *K, G, t*) = 0, *and for O* ≠ ∅ *set*

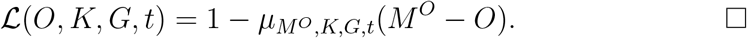

##### Proposition 11.2.4.

*Function* **ℒ** *defined in 11.2.3 is well-defined, and satisfies* 0 ≤ **ℒ** ≤ 1, *and properties (es) and (ss). If M* ∈ **𝒪** *is maximal, then* **ℒ** (*M*) = 1.

*Proof*. **ℒ** is well-defined, because *MO* is unique.

Inequality 0 ≤ **ℒ** ≤ 1 is a consequence of 0 ≤ *μ* ≤ 1.

Property (ss): If *O*_1_ ⊂ *O*_2_ ∈ **𝒪**, then 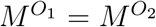 by uniqueness. Let us denote it by *M*. Hence **ℒ**(*O*_1_) = 1−*μ*_*M*_ (*M*−*O*_1_), **ℒ**(*O*_2_) = 1−*μ*_*M*_ (*M*−*O*_2_). Then 11.0.1 (2) gives the claim.

**ℒ** (*M*) = 1 is a consequence of *μ*(∅) = 0.

**Remark 11.2.5**. *Evidently, from this method, one cannot expect property (do) to be satisfied*. □

##### Proposition 11.2.6.

*If* **ℒ** *is defined as in 11.2.3, M* ∈ **𝒪** *is maximal, O*_1_, *O*_2_ ⊂ *M, then*

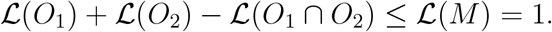

*Proof*. Set *μ* = *μ*_*M*_. Clearly *M* − *O*_1_, *M* − *O*_2_ ⊂ *M*, hence by 11.0.1 (3)

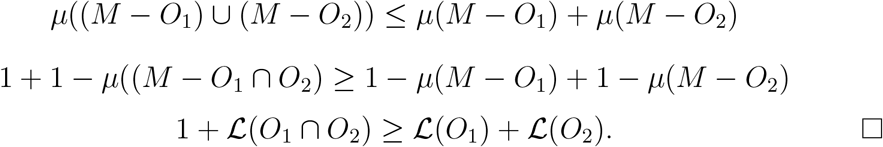

In some sense, the operation in 11.2.3 is partially the opposite of operation described in 11.1.4.

##### Proposition 11.2.7.

*Let μ*_*𝒪*_ *be defined from* **ℒ** *as in 11.1.4, and let* **ℒ***′ be defined from μ*_*𝒪*_ *as in 11.2.3*. *If O* ∈ **𝒪** *is maximal with respect to* ⊂, **ℒ** (*O*) = 1 *and O′* ⊂ *O, then* **ℒ** (*O′*) = **ℒ***′*(*O′*). □

## 12 Some final remarks on possible generalizations / modifications / usages of the model

**Remark 12.0.1**. *The presented model is comparison based i.e. the life-level of every two bio-bits can be compared. In that sense, one may say that a comparison could be enough, we would not need* 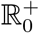 *as a range of life-level values. But the current method has some advantage. There can be changes in life-level that would not alter the order among bio-bits, but it has definite advantage if one can simply follow the change*.

**Remark 12.0.2**. *The range of* **ℒ** *is* 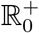 *where* 0 *is used for “not-alive”*.

*What else can be used as a possible range instead?*

1. *One could use the whole* ℝ∪{− ∞}. *In that case the value* −∞*would represent non-living. However the* log *function would give a natural isomorphism between the original concept and this one showing that they are the same in a technical point of view*.
2. *One could also try* ℕ∪{0}, *getting a discrete system of life-level values in this way*.
3. *In the presented model, the value of a life-level function is a number in* 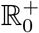, *hence every two values can be compared. One may wonder if this is a necessary requirement, i.e. a different model can be imagined with non comparable values. If one thinks that there may be bio-bits whose life-level is not comparable (i.e. none is smaller/greater than the other) then* 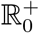 *can be replaced with a lattice (with a minimal element)*.

*E.g. the (finite) direct product of several copies of* 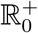 *can be used:*

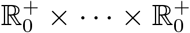 *that can be considered as a partially ordered set using the product (partial) order. Here the minimal element* (0, …, 0) *may represent if a bio-bit is not alive*.

*E.g. the direct product of some (usual) life-level functions can attain this scenario*.

1. *One may also wonder if we need any (usual) operation on the range of a life-level function, e.g. addition or multiplication on* 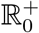, *because so far we have seldom used any of those. (Addition is used in section 11 with connection to vitality.)*
2. *The synchronized distance of two bio-bits (see section 10) may be important, hence the metric on* 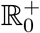 *is essential. However, for that purpose, an ordered metric space may suit as well*.

**Remark 12.0.3**. *One might consider to add the stronger condition to the definition schemas that* ⟨**𝒪**, ⊂⟩*is not only a partially ordered set but also a lattice. Similarly for* **⟨𝒦**, ⊂⟩. *It does not mean that* Dom **ℒ** *also has to be lattice type for its components, because that requirement seems to be too strong*.

**Remark 12.0.4**. *One can try to model the various decisions and/or behavior of a gorganism such that one supposes that the gorganism has its own life-level function that it evaluates in every situation, more precisely it evaluates it for all possible choices, and it chooses the one which produces the highest value*.

*One can also do it in the other way around, i.e. one can do some reverse engineering in a way that using the decisions/choices of a gorganism, one can try to find out what life-level function the gorganism uses*.

**Remark 12.0.5**. *According to section 9, one can build a similar, more specific theory on species only, where the elements of* **𝒪** *are species and the elements of* **𝒢***are group of species. In this case the elements of* **𝒦***should be some generic sub-environments too*.

## Acknowledgment

The author is grateful to professor Ivo Siekmann for his extensive support, valuable comments, advice and helpful criticism.

